# Genome-wide association study identifies seven novel loci associating with circulating cytokines and cell adhesion molecules in Finns

**DOI:** 10.1101/491852

**Authors:** Eeva Sliz, Marita Kalaoja, Ari Ahola-Olli, Olli Raitakari, Markus Perola, Veikko Salomaa, Terho Lehtimäki, Toni Karhu, Heimo Viinamäki, Marko Salmi, Kristiina Santalahti, Sirpa Jalkanen, Jari Jokelainen, Sirkka Keinänen-Kiukaanniemi, Minna Männikkö, Karl-Heinz Herzig, Marjo-Riitta Järvelin, Sylvain Sebert, Johannes Kettunen

## Abstract

**Background:** Inflammatory processes contribute to the pathophysiology of multiple chronic conditions. Genetic factors play a crucial role in modulating the inflammatory load, but the exact mechanisms are incompletely understood.

**Methods:** To add understanding to the molecular mechanisms in inflammation, we performed a genome-wide association study (GWAS) on 16 circulating cytokines and cell adhesion molecules (inflammatory phenotypes) in Northern Finland Birth Cohort 1966 (NFBC1966, N=5,284). A subsequent meta-analysis was completed for 10 phenotypes available in a GWAS of three other Finnish population cohorts adding up to 13,577 individuals in the study. Complementary association tests were performed to study the effect of the ABO blood types on soluble adhesion molecule levels.

**Results:** We identified seven novel and confirmed six previously reported loci associating with at least one of the studied inflammatory phenotypes (p<3.1×10^−9^). We observed three loci associating with the concentration of soluble vascular cell adhesion molecule-1 (sVCAM-1), one of which is the *ABO* locus that has been previously associated with soluble E-selectin (sE-selectin) and intercellular adhesion molecule-1 (sICAM-1) levels. Results from the complementary analyses suggest that the blood type B associates primarily with the concentration of sVCAM-1 while the A1 subtype shows a robust effect on sE-selectin and sICAM-1 levels. Furthermore, the genotypes in the *ABO* locus associating with higher soluble adhesion molecule levels tend to associate with lower low-density lipoprotein cholesterol level and lower cardiovascular disease risk.

**Conclusion:** The present results extend the knowledge about genetic factors contributing to the inflammatory load. Our findings suggest that two distinct mechanisms contribute to the soluble adhesion molecule levels at the *ABO* locus. The negative correlation between the genetic effects on soluble adhesion molecule levels and cardiovascular traits in this locus further suggests that increased soluble adhesion molecule levels per se may not be a risk factor for cardiovascular disease.

## Introduction

It is currently established that inflammatory load may play a role in the etiology of autoimmune and infectious diseases, but also in a broad range of other diseases, such as chronic cardio-metabolic disorders (1), neurodegenerative diseases (2), and cancer (3). The risk for these diseases increases with age (4), and due to the world’s aging population (5), their prevalence is likely to expand. Moreover, these diseases often co-occur which is likely due to shared inflammation related pathophysiology (6).

Inflammation is the body’s physiological response to harmful stimuli involving multiple molecular and cellular interactions attempting to restore disturbances in tissue or systemic homeostasis (7). Circulating cytokines, growth factors, chemokines, and cell adhesion molecules (hereafter inflammatory phenotypes) are fundamental mediators of inflammatory responses. Genes encoding these molecules and their receptors play a crucial role in mediating the related functions. Previous studies have identified loci associating with levels of inflammatory phenotypes (8–10), but the understanding of the exact regulatory mechanisms is still incomplete.

To add insights to the genetic mechanisms contributing to the inflammatory load, we performed a genome-wide association study (GWAS) of 16 circulating inflammatory phenotypes in 5,284 individuals from Northern Finland Birth Cohort 1966 (NFBC1966), and a subsequent meta-analysis of 10 phenotypes in three other Finnish population cohsorts (8) adding up to a total of 13,577 individuals in the study. We report identification of seven novel and replication of six loci associating with levels of the circulating inflammatory markers.

## Methods

### Study populations, genotyping and inflammatory phenotype quantification

#### Northern Finland Birth Cohort 1966

The Northern Finland Birth Cohort 1966 (NFBC1966) was initiated to study factors affecting preterm birth, low birth weight, and subsequent morbidity and mortality (*www.oulu.fi/nfbc*). It comprises 96% of all births during 1966 in the two northernmost provinces in Finland; altogether 12,058 children were live-born into the cohort, and the follow-ups occurred at the ages of 1, 14, 31, and 46 years (11, 12). The data analyzed in the present study is from the 31-years follow-up when clinical examinations and blood sampling was completed for altogether 6,033 individuals, 5,284 of whom had body mass index, inflammatory phenotypes and genotype data available (a maximum number of individuals per inflammatory marker 5,100). Genotyping of the samples was completed using 370k Illumina HumanHap arrays (Illumina Inc., CA, USA) and subsequent imputation was performed based on the 1000 Genome reference panel. A total of 16 inflammatory phenotypes were quantified from overnight fasting plasma samples using Bio-Rad’s Bio-Plex 200 system (BioRad Laboratories Inc., CA, USA) with Milliplex human chemokine/cytokine and CVD/cytokine kits (Cat# HCYTOMAG-60K-12 and Cat# SPR349; Millipore, St Charles, MO, USA) and Bio-Plex Manager Software version 4.3 as previously described (13). The 16 inflammatory phenotypes studied in the NFBC1966 were interleukin (IL) 1-alpha (IL1α), IL1-beta (IL1β), IL4, IL6, IL8, IL17, IL1 receptor antagonist (IL1ra), interferon gamma-induced protein 10 (IP10), monocyte chemoattractant protein 1 (MCP1), tumor necrosis factor alpha (TNFα), vascular endothelial growth factor (VEGF), plasminogen activator inhibitor 1 (PAI-1), soluble CD40 ligand (sCD40L), soluble E-selectin (sE-selectin), soluble intercellular adhesion molecule 1 (sICAM-1) and soluble vascular cell adhesion molecule 1 (sVCAM-1).

#### GWAS summary statistics from three Finnish population cohorts

Meta-analyses were conducted for 10 phenotypes available in a previous GWAS (8). The study included up to 8,293 Finnish individuals from The Cardiovascular Risk in Young Finns Study (YFS) (14) and FINRISK (*www.thl.fi/finriski*) (15) adding up to 13,577 individuals studied in the present meta-analyses. Shortly, YFS is a population-based follow-up study started in 1980 comprising randomly chosen individuals from Finnish cities Helsinki, Kuopio, Tampere, Oulu, and Turku. The YFS data included in the previous GWAS is from 2,019 individuals who participated in the follow-up in 2007 and who had both inflammatory phenotype and genotype data available. FINRISK is a Finnish population survey conducted every five years to monitor chronic diseases and their risk factors. The surveys use independent, random, and representative samples from different geographical areas of Finland. The data included in the present meta-analyses were from participants of the 1997 and 2002 surveys. Genotypes were obtained using 670k Illumina HumanHap arrays (Illumina Inc., CA, USA) and imputed based on 1000 Genome reference panel. Inflammatory markers were quantified using Bio-Rad’s premixed Bio-Plex Pro Human Cytokine 27-plex Assay and 21-plex Assay, and Bio-Plex 200 reader with Bio-Plex 6.0 software (Bio-Rad Laboratories Inc., CA, USA) as previously described (16, 17). Samples were serum in YFS, EDTA plasma in FINRISK1997, and heparin plasma in FINRISK2002.

### Statistical analyses

#### GWAS and meta-analysis

To allow meta-analysis between the present results and the previous GWAS, the data processing and analysis model were done according to Ahola-Olli *et al.* (8): Preceding the analyses, linear regression models were fitted to adjust the inflammatory phenotypes for age, sex, BMI, and ten first genetic principal components to control for population stratification. The resulting residuals were normalized with inverse-rank based transformation, and the adjusted and transformed residuals were used as phenotypes in the analyses. Genome-wide association tests were performed using snptest_v2.5.1 software (18, 19). Allele effects were estimated using an additive model (-frequentist 1) and the option to center and scale the phenotypes was disabled (-use_raw_phenotypes). The GWAS results were filtered by including markers with model fit info > 0.8 and minor allele count > 10. Filtered data was used to perform meta-analyses by METAL software (v.2011-03-25) (20) for the 10 phenotypes (IL1β, IL1ra, IL17, IL4, IL6, IL8, IP10, MCP1, TNFα, and VEGF) available in the previous GWAS (8). Genomic control correction was enabled to account for population stratification and cryptic relatedness.

#### Supplemental genome-wide tests in NFBC1966

Individuals showing symptoms of an acute infection were omitted from the supplemental genome-wide tests performed in the NFBC1966 population. Here, individuals reported having fever at the time of the blood sampling and individuals having C-reactive protein (CRP) level > 10 mg/l were excluded. Otherwise the analysis models were as above.

#### Conditional analyses and variance explained

To assess whether the identified loci harbor multiple independent association signals, we conducted conditional analyses by further adjusting the models with the locus-specific lead variants. The association tests were repeated within a 2Mb window around the lead SNP for the phenotypes studied in the NFBC1966 population only. For the meta-analyzed phenotypes, we applied a method proposed by Yang *et al.* that enables conditional analyses of GWAS summary statistics (21, 22). NFBC1966 was used as a reference sample to estimate linkage disequilibrium (LD) corrections in these analyses. The proportion of variance explained was calculated using all independent variants using the following formula:

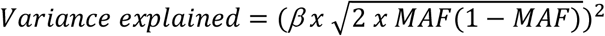

Here *β* is the variant’s effect estimate on the inflammatory phenotype and MAF denotes minor allele frequency.

#### Complementary association tests on soluble adhesion molecule levels

Complementary association tests within a 2Mb window were conducted to better evaluate the effect of the ABO blood type on the association with soluble adhesion molecule levels at the *ABO* locus. For sE-selectin, sICAM-1 and sVCAM-1 levels, linear models were fitted by further adjusting for the ABO blood type or rs507666 genotype tagging the A1 subtype (23).

In addition, we determined the effect estimates of ABO blood types and ABO blood types stratified by rs507666 genotype on sE-selectin, sICAM-1 and sVCAM-1 levels in linear models. Here, adjusted and transformed soluble adhesion molecule concentrations were as outcomes and ABO blood types as categorical variables (blood type A versus non-A, etc.); corresponding models were fitted for the stratified blood types (blood type A with rs507666 G/G versus others, etc.)

#### Correlations of the genetic effects

As previous evidence suggests that the elevated concentrations of circulating markers of inflammation increase the risk of cardiovascular diseases (CVD) (24, 25), we further evaluated how variants at the loci associating with inflammatory phenotypes may relate to other cardiovascular traits. We extracted SNP effects on coronary artery disease (CAD) risk, stroke risk, and low-density lipoprotein cholesterol (LDL-C) or high-density lipoprotein cholesterol (HDL-C) levels from open-access data provided by CARDIoGRAM (26), Stroke Consortia (27), and a metabolomics GWAS (28). First, data were filtered to include only the SNPs available in all the three data sets within a 1Mb window around each lead variant. Next, subsets of representative SNPs at the each of the significant loci were extracted using clumping function in PLINK 1.90b4.1 (29). Here, NFBC1966 was used as the reference sample to construct LD structures and *r*^2^=0.2 was used as the LD threshold while other parameters were as by default. The subsets of SNPs were used to determine the linear relationships of the genetic effects (Z-scores) on inflammatory phenotypes versus other traits for each significant loci identified in the present GWAS.

## Results

Basic characteristics of the NFBC1966 study population is provided in Table 1. Inflammatory phenotype distributions are tabulated in Table S1 and their correlation structure is shown in Figure S1. Using a threshold of p<3.1×10^−9^ for statistical significance (standard genome-wide significance level p<5×10^−8^ corrected for 16 phenotypes tested), we identified seven novel and six previously reported loci associating with one or more of the inflammatory phenotypes. The results are summarized in Table 1 and combined Manhattan plots are shown in Figure 1. Manhattan plots and Q-Q plots for each inflammatory phenotype are provided in the supplement (Figure S2 A-Z). Genomic inflation factor values range between 0.99-1.02 suggesting no inflation in the test statistics (Table S2). Table S3 lists traits associated previously with the loci showing novel associations with inflammatory phenotypes in the present study.

**Table 1.** Basic charcteristics of the NFBC1966 study population.

| Characteristics |  |
| --- | --- |
| Total number of individuals | 5284 |
| Number of males (%) | 2543 (48.1) |
| Age, years | 31.1 ± 0.4 |
| BMI, kg/m<sup>2</sup> | 24.4 ± 4.0 |
| Glucose, mmol/l | 5.1 ± 0.7 |
| LDL-cholesterol, mmol/l | 3.0 ± 0.9 |
| HDL-cholesterol, mmol/l | 1.6 ± 0.4 |
| Systolic blood pressure, mmHg | 124.2 ± 13.6 |
| Diastolic blood pressure, mmHg | 76.8 ± 11.7 |
| Values are mean ± standard deviation. |  |

**Table 2.**
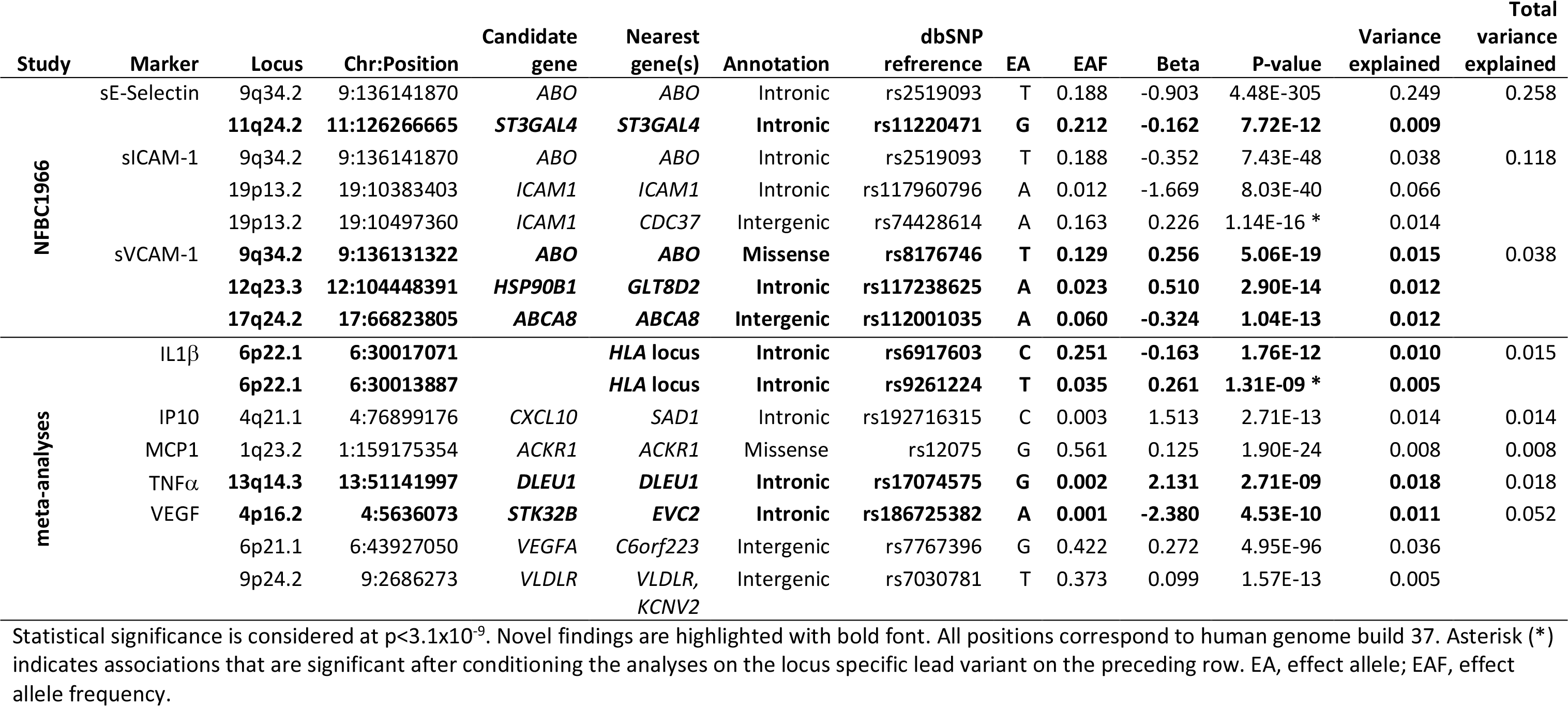
Significant loci associating with the circulating inflammatory phenotypes.

**Figure 1.**
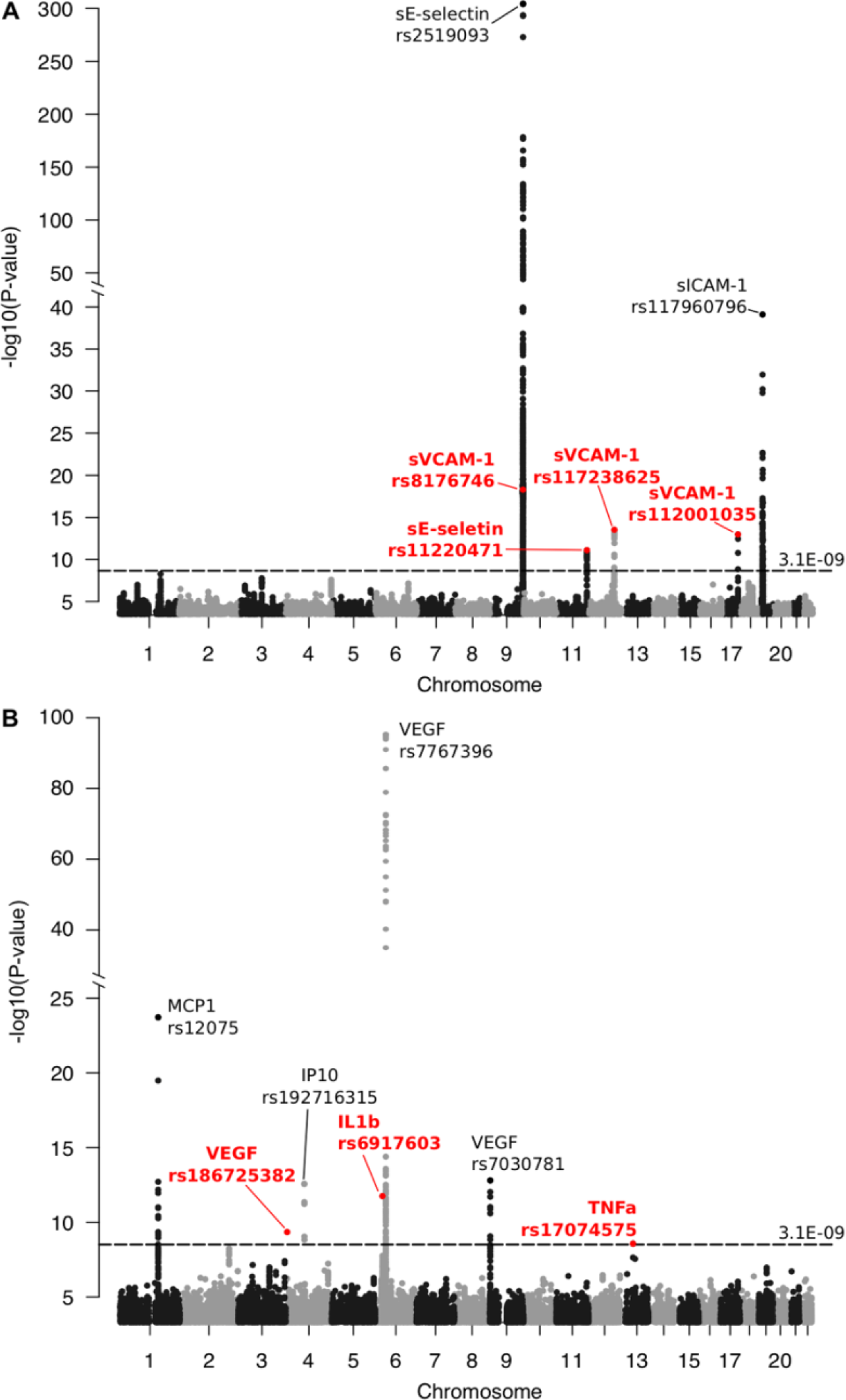
The combined Manhattan plots for significant associations with inflammatory markers studied in (A) NFBC1966 and in (B) meta-analyses with three other Finnish population cohorts. Significance threshold p<3.1×10^−9^ derives from the standard p-value limit for genome-wide significance p<5×10^−8^ corrected for 16 markers examined in the present study. Novel association signals are highlighted with red font and replicated loci are marked with black font.

### Cell adhesion molecules

#### The ABO locus shows large effects on sE-selectin, sICAM-1, and sVCAM-1 levels

We observed a novel effect on sVCAM-1 concentration in 9q34.2 near *ABO* (ABO, alpha 1-3-N-acetylgalactosaminyltransferase and alpha 1-3-galactosyltransferase) in the NFBC1966 population. This locus showed a robust association also with sE-selectin and sICAM-1 concentrations as previously reported (23, 30, 31). Noteworthy, the lead variant for sE-selectin and sICAM-1 associations (rs2519093) was different from the lead variant for sVCAM-1 association (rs8176746). The former variant is in LD (*r*^*2*^=1 in NFBC1966) with rs507666 tagging the ABO blood type A subtype A1 whereas the latter variant tags the blood type B (23).

To better evaluate if the ABO blood types constitute the molecular mechanism explaining the association between the *ABO* locus and soluble adhesion molecule levels, we completed supplementary association tests further adjusted for ABO blood type or rs507666 genotype indicative of the A1 subtype. The results of the supplementary tests suggested that the association of the rs8176746 with sVCAM-1 concentration is independent of the A1 subtype (p=4.98×10^−15^ for the rs507666-adjusted association). On the contrary, the associations of the rs2519093 with concentrations of sE-selectin and sVCAM-1 remained highly significant when adjusted for ABO blood type (p=3.40×10^−123^ and p=3.43×10^−17^, respectively). Statistical significances were abolished when rs8176746 association with sVCAM-1 was adjusted for ABO blood type and rs2519093 association with sE-selectin or sICAM-1 was adjusted for rs507666.

We further determined the effect estimates of the ABO blood types and ABO blood types stratified by rs507666 genotype on soluble adhesion molecule levels. The blood type A showed negative associations with the levels of all the three adhesion molecules and the effect was the most robust on the sE-selectin level (Figure 2, Panel 1). However, major discrepancies in the effect directions were seen when the analyses were stratified by the rs507666 genotype (Figure 2, Panel 2). Congruent with previous reports (23, 30), the present results suggest that the A1 subtype/rs507666 genotype influences sE-selectin or sVICAM-1 levels. In contrast, the blood type B seems to attribute predominantly to sVCAM-1 level while the A1 subtype/rs507666 shows only a modest effect on sVCAM-1.

**Figure 2.**
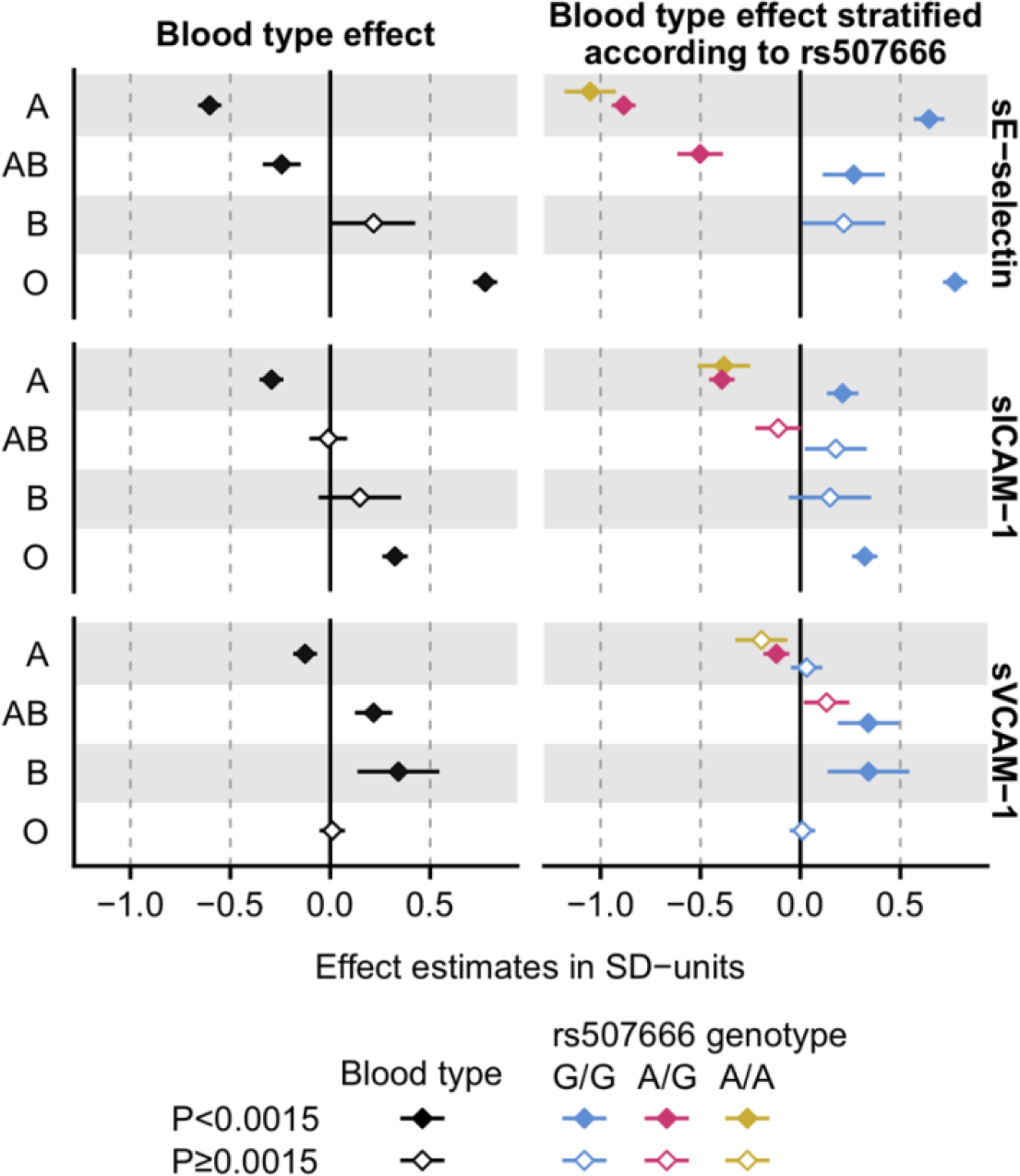
The effects of the ABO blood types and the A1 subtype on soluble adhesion molecule levels. The effects of the ABO blood types on sE-selectin, sICAM-1 and sVCAM-1 levels were evaluated in linear models, where adjusted (sex, age, BMI, and the ten first genetic principal components) and transformed soluble adhesion molecule concentrations were used as outcomes and the ABO blood type served as categorical variable (A versus non-A, etc.). Corresponding models were fitted for the ABO blood types stratified by the rs507666-A allele count (0, 1 or 2), where the A allele tags the ABO subtype A1 having enhanced glycosyltransferase activity (25). No individuals were found to have B or O blood type and one or more copies of the rs507666-A allele and, thus, it was not possible to perform stratification within these blood types.

#### HSP90B1 and ABCA8 loci associate with sVCAM-1 levels

We identified two other novel loci for sVCAM-1 (12q23.3 and 17q24.2) in the NFBC1966 population. In chr12 the lead variant rs117238625 is in LD (*r*^*2*^=1 in NFBC1966) with rs117468318 locating in the 5’ UTR region of *HSP90B1* (heat shock protein 90kDa beta member 1) and, according to RegulomeDB (32), is likely to affect transcription factor binding providing evidence for a possible regulatory mechanism. Variants in this locus have been previously associated with stem cell growth factor beta levels (8) and corneal structure (33). The association signal in chr17 locates near *ABCA8* (ATP binding cassette subfamily A member 8) encoding one of the ATP binding cassette transporters. Other studies have identified associations in this locus with HDL and LDL cholesterol levels (34–37), breast cancer risk (38), heart’s electrical cycle related traits (QT interval, QRS duration) (39–41).

#### Variations in sialyltransferase encoding genes show an effect on sE-selectin level

For sE-selectin level, we identified a novel association in 11q24.2 in the region of *ST3GAL4* (ST3 beta-galactoside alpha-2,3-sialyltransferase 4). Other studies have associated other variants in this region with mean platelet volume and platelet count (42), LDL cholesterol levels (34, 36) or pleiotropic associations with LDL cholesterol and C-reactive protein (43), blood protein levels (44) and liver enzyme levels (45). We identified a suggestive signal with sE-selectin level also in 3q12.1 near *ST3GAL6* (ST3 beta-galactoside alpha-2,3-sialyltransferase 6), but the association was not significant after multiple correction (p=1.75×10^−08^). Both of the sialyltransferase genes have been implicated in the production of functional E-, P-, and L-selectin ligands in mice (46).

#### Two independent association signals on sICAM-1 level near ICAM1

We replicated the previously reported association for sICAM-1 level in 19p13.2 near *ICAM1* (intracellular adhesion molecule 1) (23, 44). When the primary association test was conditioned for the lead variant rs117960796, another significant association was detected (rs74428614, p=1.14×10^−16^) indicative of more than one independent variant contributing to sICAM-1 level in this locus.

### Vascular endothelial growth factor

In the meta-analyses, we identified a novel locus 4p16.2 with a large effect on VEGF (β=−2.38 SD). This locus harbours genes *EVC* (EvC ciliary complex subunit 1), *EVC2* (EvC ciliary complex subunit 2), and *STK32B* (serine/threonine kinase 32B). Mutations in this locus have been associated previously with Celiac disease (47), coronary heart disease (48), essential tremor (49), and Ellis van Creveld syndrome (50, 51). In addition, we replicated two previously reported loci associating with VEGF levels in 6p21.1 near *VEGFA* (vascular endothelial growth factor A) and in 9p24.2 near *VLDLR* (very-low-density lipoprotein receptor) (8).

### Pro-inflammatory cytokines

#### Locus near DLEU1 shows a large effect on TNFa

We identified a novel variant with a large effect on TNFα levels (β=2.13 SD) in 13q14.3 near *DLEU1* and *DLEU7* (deleted in lymphocytic leukemia 1 and 7) in the meta-analyses. Previous GWAS findings in this locus include associations with reticulocyte related traits (42) and tooth development (52).

#### The HLA locus shows a small effect on IL-1β

A novel variant at 6p22.1 in the human leukocyte antigen locus associating with IL-1β level was identified in the meta-analyses. In the conditional analyses, we observed two independent association signals at this locus (Table 1, Figure S2 J). The same locus and the same lead variant rs6917603 showed also a suggestive effect on IL4 level (Figure S2 L), but the meta-analysed result was not significant after multiple correction (p=5.56×10^−09^). Variants in this region have been associated previously with multiple immune system related traits such as white blood cell count (42), but also with several other traits, including lipid metabolism phenotypes (53), schizophrenia (54) and lung cancer (55).

### Chemokines

We replicated previously reported loci near *CXCL10* (C-X-C motif chemokine ligand 10) and *ACKR1* (atypical chemokine receptor 1) associating with IP10 levels and with MCP1 levels, respectively (8).

### Supplemental genome-wide tests in NFBC1966

Altogether 236 individuals having fever or CRP>10 mg/l were excluded from the supplemental genome-wide tests performed in the NFBC1966 population. The results of the supplementary analyses were congruent with the original findings (Table S4).

### Comparisons of SNP effects on inflammatory markers versus other traits

Elevated circulating concentrations of inflammatory markers is a risk factor for cardiovascular diseases (24, 25). In order to add insights how genetic variants associating with inflammatory phenotypes contribute to other cardiovascular health-related traits, we compared the SNP effects (Z-scores) at each significant locus versus corresponding SNP effects obtained from open-access data (26–28). The list of SNPs used to determine the correlations of the SNP effects on inflammatory phenotypes versus other traits is provided in Table S5. At the *ABO* locus, we observed a negative linear relationship between the SNP effects on sE-selectin and sICAM-1 levels and CAD risk, stroke risk as well as LDL-C and HDL-C levels (Figure 3A-B, Figure S3 A-B). For sVCAM-1 at the same locus, the overall trend was similar, but correlations did not reach statistical significance due to a low number of representative SNPs obtained in clumping (Figure 3C, Figure S3 C). The results at the other loci are provided in Figures S3 D-I.

**Figure 3.**
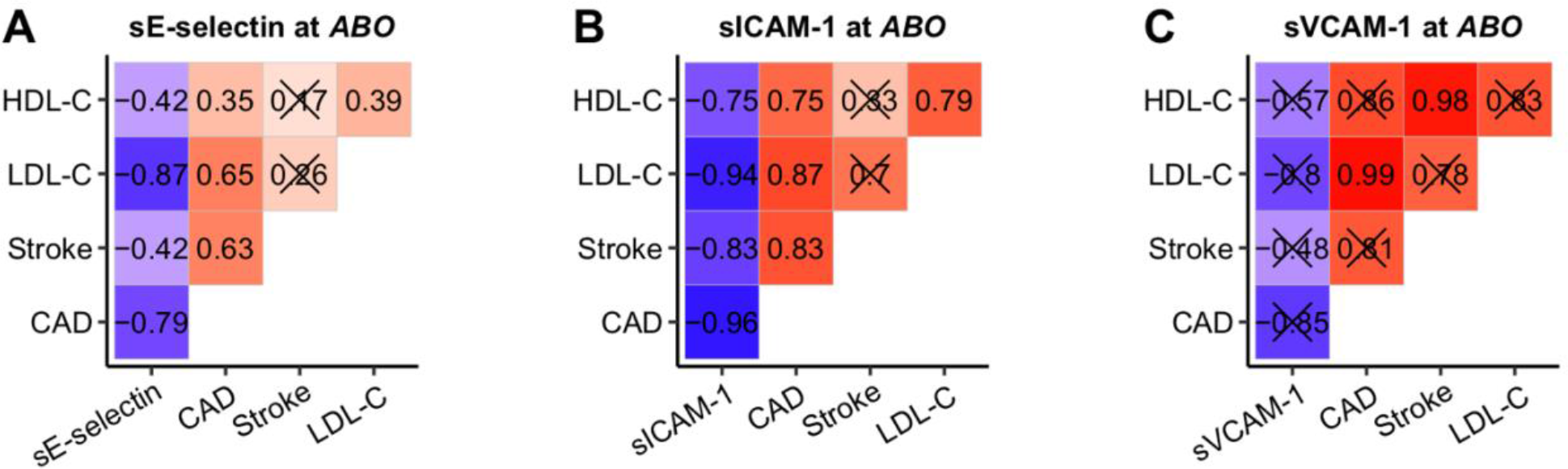
The correlations of the SNP effects on soluble adhesion molecule levels versus other cardiovascular health-related traits at the *ABO* locus. The correspondence of the SNP effects on inflammatory phenotypes versus cardiovascular health-related traits were determined for the SNPs associating with (A) sE-selectin, (B) sICAM-1, and (C) sVCAM-1 levels at the *ABO* locus. SNP effects on coronary artery disease (CAD), stroke, and LDL-C or HDL-C were extracted from CARDIoGRAM (26), Stroke Consortia (27), and a metabolomics GWAS (28) summary statistics, respectively. The correlations (Pearson’s *r*) of the genetic effects were estimated using subsets of representative SNPs extracted from the summary statistics of the present GWAS using a clumping function in PLINK and *r*^*2*^ threshold of 0.2. Prior to clumping, data were filtered to include only the SNPs available in all the three datasets. Correlations with p≥0.05 are marked with a cross. The SNPs used for estimating the correlations are listed in Table S5 and scatter plot representations as well as correlations at the other loci and other inflammatory phenotypes are shown in Figure S3.

## Discussion

The present study examines genetic determinants of 16 circulating inflammatory phenotypes in 5,284 individuals from Northern Finland with a subsequent meta-analysis of 10 phenotypes in three other Finnish populations adding up to a total of 13,577 participants. We report seven novel and replication of six previously published genetic associations.

We detected a novel association for sVCAM-1 concentration at the *ABO* locus. This locus showed robust associations also with sE-selectin and sICAM-1 concentrations congruent with previous studies (23, 30, 31). The present GWAS findings suggested two distinct association signals at the *ABO* locus for the sE-selectin and sICAM-1 levels versus sVCAM-1 level. The results of the supplementary tests supported the perception that genetic variants in this locus may regulate the circulating concentration of adhesion molecules by at least two different mechanisms. The first mechanism comprises the blood type A subtype A1, tagged by rs507666-A, that has a robust lowering effect on sE-selectin and sICAM-1 levels (23, 30). The second mechanism involves the blood type B that seems to have an increasing effect on sVCAM-1 level. Others have suggested that the lowering effect of the A1 subtype on sE-selectin and sICAM-1 could arise from increased glycosyltransferase activity that possibly modifies the shedding of the adhesion molecules from the endothelium and/or their clearance rate from circulation (23, 30, 56). To the best of our knowledge, the underlying mechanism explaining the association between the blood type B and the higher sVCAM-1 concentration remains unknown and warrants research. VCAM-1-mediated adhesion involves interaction with galectin-3, a protein that has a specificity for galactosides (57, 58). As the B antigen holds an additional galactose monomer compared with the A and O antigens, and galectins are known to recognize blood type antigens (59), it raises the speculation that the amount of unbound sVCAM-1 in the circulation could be influenced by a possible competitive binding of galectin-3 with sVCAM-1 and the B antigen.

To evaluate how variants in the *ABO* locus may relate to other health outcomes, we compared the correspondence of genetic effects on soluble adhesion molecule levels versus cardiovascular health-related traits. We observed that the genetic effects on adhesion molecule levels were inclined to show a negative correlation with the genetic effects on LDL-C and HDL-C levels as well as lower risk for CAD and stroke. This denotes that the genotypes at the *ABO* locus associating with increased levels of soluble adhesion molecules tended to associate with lower circulating cholesterol level as well as lower risk of cardiovascular outcomes. This was unexpected since according to previous evidence increased soluble adhesion molecule levels are linked with atherosclerosis progression and vascular outcomes (25, 60, 61). Possible explanations unravelling the negative correlation advocate that soluble adhesion molecules may compete with leukocyte adhesion to the endothelial molecules or that enhanced ectodomain shedding may contribute to the reduced recruitment of leukocytes to the subendothelial space thereby promoting cardioprotective effects (62). Additionally, the observed negative relationship between the genetic effects on soluble adhesion molecule and LDL-C levels suggests that altered cholesterol metabolism could contribute to the CAD risk associated with the *ABO* locus; the genetic effects of the same SNPs on LDL-C versus CAD risk showed a positive correlation (Figure 3). Nevertheless, further studies are warranted to understand the exact mechanisms.

Another novel association with sVCAM-1 level was detected in chr12 near *HSP90B1* encoding heat shock protein gp96, a chaperone that is essential for assembly of 14 of 17 integrin pairs in the hematopoietic system (63). Integrin α4β1, also known as very late antigen (VLA)-4, is an important ligand of VCAM-1 (64). The lead SNP of this locus is in LD with rs117468318 (*r*^*2*^=1 in NFBC1966) that locates in the 5’UTR region of *HSP90B1* and, according to RegulomeDB (32), is likely to affect transcription factor binding suggesting a possible regulatory mechanism for the detected association. If altered transcription of *HSP90B1* had a downstream effect on integrin α4β1 level, this could further modify the level of unbound sVCAM-1 in circulation.

The 3^rd^ novel locus showing association with sVCAM-1 level was identified in chr17 near *ABCA8*. The lead SNP rs112001035 is an eQTL for *ABCA8* in multiple tissue types (65). ABCA8 has been shown to regulate levels of HDL-cholesterol with a mechanism that likely involves interaction with ABCA1 (66). If ABCA8 is involved in regulation of HDL level (66) and if plasma HDL levels contribute to VCAM-1 expression (67, 68), then altered expression of the *ABCA8* could influence circulating levels of sVCAM-1 via modulating HDL particle concentration. However, this hypothesis is not supported by the fact that the effect of the lead SNP on HDL particle concentration is negligible in a large metabolomics GWAS (β=−0.043 SD, p=0.049) (28). There is evidence suggesting that ABCA8 may be involved in sphingolipid metabolism (69) and it has been hypothesized that ABCA8 may be involved in the formation of specific membrane domains during ApoA-I lipidation (66). Thus, one could speculate that the association between the ABCA8 locus and sVCAM-1 level could be related to altered HDL composition rather than absolute particle concentration, which could contribute to altered endothelial homeostasis. However, more evidence is needed to draw conclusions.

We detected a novel effect of rs11220471 in chr11 near *ST3GAL4* on sE-selectin levels in the NFBC1966 population. *ST3GAL4* encodes a member of the glycosyltransferase 29 family of enzymes involved in protein glycosylation. In mice, St3Gal4 is needed for synthesis of functional selectin ligands (46) and it has been shown to participate to chemokine C-C motif ligand 5 (Ccl5)-dependent myeloid cell recruitment to inflamed endothelium (70). The altered levels or structure of selectin ligands due to variation in *ST3GAL4* could contribute to the levels of unbound sE-selectin in circulation, providing a biologically rational mechanism for the detected association.

In the meta-analyses, we detected a novel large effect locus for VEGF in chr4 (β=−2.38 SD) near genes *EVC* (EvC ciliary complex subunit 1), *EVC2* (EvC ciliary complex subunit 2) and *STK32B* (serine/threonine kinase 32B). Mutations in this locus have been associated previously with celiac disease (47), coronary artery disease (48), and Ellis-van Creveld syndrome, a rare recessive disorder characterized with congenital defects such as short ribs, postaxial polydactyly, growth retardation, and ectodermal and cardiac defects (50, 51). The expression level of *STK32B* has been associated with clinicopathological features of oral cavity cell carcinoma including peritumoral inflammatory infiltration, metastatic spread to the cervical lymph nodes, and tumour size (71). STK32B may play a role in the hedgehog signalling pathway, which has been implicated in metastasis and angiogenesis in cancer (71) and downregulated in celiac disease (72). The hedgehog signalling has shown to be involved in the regulation of VEGF expression during developmental angiogenesis in avian embryo (73). Thus, previous literature and our results advocate that STK32B may be involved in the regulation of VEGF levels possibly via hedgehog signalling-related mechanism.

The other novel findings obtained in meta-analysis include a large effect locus on TNFα level in chr13 (β=2.13 SD). The locus in 13q14.3 associating with TNFα locates near *DLEU1* and *DLEU7* (Deleted in Lymphocytic Leukemia 1 and 7). This region is recurrently deleted in tumours and hematopoietic malignancies (74, 75). *DLEU1* is a part of a transcriptionally coregulated gene cluster that modulates the activity of the NF-kB pathway (76) which is also modulated by TNFα (77). It is largely unknown how the DLEU1 and related DLEU2 regulate NF-kB activity (78); our result suggests that TNFα signalling might be involved in this mechanism.

At last, we identified a small effect locus in chr6 harbouring two independent association signals on IL-1β and showing suggestive association also on IL4 level. This association signal is in the region coding the human leukocyte antigen proteins, and further experimental evidence would be needed to identify the exact mechanism how the locus contributes to interleukin levels.

The strengths and limitations of our study should be considered. The sample size of the present study should provide adequate power for detecting genetic associations with circulating markers of systemic inflammation (9). The use of genetically isolated populations, such as inhabitants of Northern Finland, should further enhance the power for locus identification in GWAS settings (79). We were able to perform meta-analyses only for 10 out of the 16 inflammatory phenotypes analysed in the NFBC1966 population and, thus, a replication of the present findings in other populations would be helpful. The inter-assay coefficient of variability measures for sE-selectin and VEGF in particular are notably larger than 15% that is considered to be the limit for acceptable values (Table S1). However, to our consideration, all the findings identified in the present study locate on genome regions with biologically relevant genes. Furthermore, the extremely small p-values (p=4.48×10^−305^ for sE-selectin at the *ABO* locus and p=4.95×10^−96^ for VEGF at the *VEGFA* locus) and the replications of the previously reported loci speak for the data adequacy and adds confidence to the novel associations. Due to mismatches in genotype availability between the present results and open-access data sets or low number of SNPs obtained after exclusions, it was not possible to carry out meaningful comparisons of the genetic effects at all the significant loci.

The present results provide novel information on genetic mechanisms influencing levels of inflammatory phenotypes in circulation. The evident role of the *ABO* locus in the regulation of the soluble adhesion molecule levels in circulation likely encompasses at least two distinct mechanisms influencing sE-selectin, sICAM-1 and sVCAM-1 levels. Our findings provide evidence that increased soluble adhesion molecule concentrations per se may not be a risk factor for cardiovascular outcomes. In particular, genetic variation associating with increased sE-selectin or sICAM-1 levels at the *ABO* locus seem to contribute to lower CAD risk. Furthermore, genetic effects at the *ICAM1* locus providing a direct molecular link to sICAM-1 concentration do not correlate with the genetic effects on CAD risk nor stroke risk. Overall, the present study extends the knowledge about the precise molecular pathways involved in inflammatory load.

## Supporting information

## Acknowledgements

We gratefully acknowledge the contributions of the participants in the Northern Finland Birth Cohort 1966 study. We also thank all the field workers and laboratory personnel for their efforts. Data on coronary artery disease / myocardial infarction have been contributed by CARDIoGRAMplusC4D investigators and have been downloaded from www.CARDIOGRAMPLUSC4D.ORG.

## Abbreviations

GWAS: genome-wide association study
IL1α: interleukin 1-alpha
IL1β: interleukin 1-beta
IL1ra: interleukin 1 receptor antagonist
IL4: interleukin 4
IL6: interleukin 6
IL8: interleukin 8
IL17: interleukin 17
IP10: interferon gamma-induced protein 10
MAF: minor allele frequency
MCP1: monocyte chemoattractant protein 1
NFBC1966: Northern Finland Birth Cohort 1966
PAI-1: plasminogen activator inhibitor 1
sCD40L: soluble CD40 ligand
sE-selectin: soluble E-selectin
sICAM-1: soluble intercellular cell adhesion molecule 1
sVCAM-1: soluble vascular cell adhesion molecule 1
TNFα: tumor necrosis factor alpha
VEGF: vascular endothelial growth factor

