## Supplementary material for "Genome-wide association study identifies seven novel loci associating with circulating cytokines and cell adhesion molecules in Finns"

**Content**

Table S1. Distributions of the unadjusted and untransformed inflammatory phenotypes in the NFBC1966. 1
Table S2. Genomic inflation factors for the 16 studied inflammatory markers. 2
Table S3. Traits associated previously with the novel loci identified in the present study. 3
Table S4. The results of the supplemental GWAS. 7
Table S5. The list of SNPs used to determine the correlations of the SNP effects on inflammatory phenotypes versus other traits. 8
Figure S1. Correlation heatmap of the inflammatory phenotypes in NFBC1966. 12
Figure S2. A-Z) Manhattan. Q-Q. and LocusZoom plots for all the inflammatory phenotypes studied. 13
Figure S3. A-I) SNP effect correlations as scatter plots. 39

**Table S1. Distributions of the unadjusted and untransformed inflammatory phenotypes in the NFBC1966.**

| | | | | | | Inter-assay | Intra-assay |
|---|---|---|---|---|---|---|---|
| **Phenotype** | **N** | **Mean** | **SD** | **Median** | **IQR** | **CV %** | **CV %** |
| sE-selectin (ng/ml) | 5100 | 33.39 | 15.56 | 31.56 | 18.61 | 28.0 | 7.5 |
| sICAM-1 (ng/ml) | 5100 | 145.35 | 60.49 | 134.81 | 53.70 | 15.9 | 8.3 |
| sVCAM-1 (ng/ml) | 5100 | 1644.40 | 396.67 | 1609.73 | 481.99 | 13.9 | 7.9 |
| VEGF (pg/ml) | 4941 | 222.96 | 2840.43 | 62.77 | 164.39 | 20.6 | 7.5 |
| IL17 (pg/ml) | 4974 | 77.68 | 1331.38 | 21.86 | 40.76 | 8.4 | 2.5 |
| IL1α (pg/ml) | 4917 | 77.81 | 690.51 | 7.17 | 41.88 | 11.0 | 4.0 |
| IL1β (pg/ml) | 4971 | 7.48 | 87.60 | 0.72 | 5.34 | 9.5 | 3.0 |
| IL1ra (pg/ml) | 4864 | 85.02 | 1088.30 | 10.78 | 34.15 | 13.0 | 4.6 |
| IL4 (pg/ml) | 4962 | 49.16 | 912.08 | 2.36 | 19.45 | 10.4 | 3.0 |
| IL6 (pg/ml) | 4966 | 17.85 | 124.09 | 2.45 | 8.85 | 7.7 | 2.6 |
| IL8 (pg/ml) | 4974 | 46.20 | 454.39 | 16.33 | 24.99 | 7.6 | 2.2 |
| IP10 (pg/ml) | 4975 | 442.61 | 433.54 | 364.51 | 231.40 | 8.3 | 4.5 |
| MCP1 (pg/ml) | 4975 | 309.68 | 128.31 | 294.46 | 143.97 | 6.7 | 3.1 |
| sCD40L (pg/ml) | 4970 | 567.59 | 975.28 | 395.33 | 339.14 | 14.9 | 3.7 |
| TNFα (pg/ml) | 4971 | 10.99 | 97.63 | 5.94 | 4.06 | 7.2 | 2.6 |
| PAI-1 (ng/ml) | 5100 | 20.99 | 28.43 | 11.58 | 17.73 | 18.6 | 6.8 |

SD, standard deviation; IQR, interquartile range; CV, coefficient of variation.

**Table S2. Genomic inflation factors for the 16 studied inflammatory markers.**
The genomic inflation factors were calculated in R as median(Z^2)/qchisq(0.5,1), where Z is the effect estimate per standard error ratio.

| Marker | Genomic lambda |
|---|---|
| sE-selectin | 1.024 |
| sICAM-1 | 1.022 |
| sVCAM-1 | 1.011 |
| VEGF | 0.995 |
| IL17 | 1.006 |
| IL1$\alpha$ | 1.023 |
| IL1$\beta$ | 1.007 |
| IL1ra | 0.989 |
| IL4 | 0.998 |
| IL6 | 0.992 |
| IL8 | 1.000 |
| IP10 | 0.989 |
| MCP1 | 0.991 |
| sCD40L | 1.006 |
| TNF$\alpha$ | 0.990 |
| PAI-1 | 1.005 |

**Table S3. Traits associated previously with the novel loci identified in the present study.**

If multiple studies reported associations with the same trait or if multiple SNPs were associated with the same trait, the strongest SNP with the smallest P-value is given.

| | Trait | P-value | SNP | PMID |
|---|---|---|---|---|
| ***sE-selectin in NFBC1966; ST3GAL4; 11q24.2; 11:126266665; rs11220471*** | | | | |
| Biobank Engine (inquiry the lead SNP) | na | | | |
| Phenoscanner (inquiry the lead SNP) | Mean platelet volume | 8.81e-06 | rs11220471 | 27863252 |
| GWAS Catalog (inquiry +-500 kb around the lead SNP) | Blood protein levels (Cell adhesion molecule-related/down-regulated by oncogenes, CDON.4541.49.2) | 4.00e-86 | rs562022020 | 29875488, 28240269 |
| | C-reactive protein levels or LDL-cholesterol levels (pleiotropy) | 6.00e-17 | rs11220463 | 27286809 |
| | Cholesterol, total | 2.00e-15 | rs11220462 | 28334899, 24097068, 20686565, 25961943 |
| | Eosinophil percentage of granulocytes | 1.00e-10 | rs8177376 | 27863252 |
| | LDL cholesterol | 2.00e-22 | rs17135399 | 28334899, 24097068, 20686565, 25961943 |
| | Liver enzyme levels (alkaline phosphatase) | 2.00e-09 | rs2236653 | 22001757 |
| | Lung function (FEV1) | 5.00e-10 | rs567508 | 28166213 |
| | Mean platelet volume | 3.00e-30 | rs7949566 | 27863252 |
| | Mumps | 1.00e-08 | rs3862630 | 28928442 |
| | Neutrophil percentage of granulocytes | 3.00e-12 | rs8177376 | 27863252 |
| | Neutrophil percentage of white cells | 5.00e-10 | rs73017399 | 27863252 |
| | Platelet count | 2.00e-18 | rs4937127 | 27863252 |
| | Platelet distribution width | 4.00e-13 | rs7949566 | 27863252 |
| | Primary tooth development (time to first tooth eruption) | 4.00e-08 | rs4937076 | 23704328 |
| | Serum alkaline phosphatase levels | 6.00-41 | rs10893506 | 29403010 |
| ***sVAM-1 in NFBC1966; ABO; 9q34.2; 9:136131322; rs8176746*** | | | | |
| Biobank Engine (inquiry the lead SNP) | Red blood cell (erythrocyte) count | 5.13e-24 | rs8176746 | (https://biobankengine.stanford.edu) |
| | Platelet crit. | 3.15e-18 | rs8176746 | (https://biobankengine.stanford.edu) |
| | Impedance of whole body | 5.60e-13 | rs8176746 | (https://biobankengine.stanford.edu) |
| | Immature reticulocyte fraction | 2.73e-10 | rs8176746 | (https://biobankengine.stanford.edu) |
| | Deep venous thrombosis | 7.89e-15 | rs8176746 | (https://biobankengine.stanford.edu) |
| Phenoscanner (inquiry the lead SNP) | Reed blood cell count + multiple other red bood cell-related traits | 1.95e-53 | rs8176746 | 27863252 |
| | Plasma carcinoembryonic antigen CEA levels | 4.70e-36 | rs8176746 | 23300138 |
| | Activated partial thromboplastin time | 1.77e-22 | rs8176746 | 22703881 |
| | Von Willebrand factor vWF | 1.59e-17 | rs8176746 | 23381943 |
| | Factor XIII antigen | 1.19e-14 | rs8176746 | 23381943 |
| | Blood clot in the leg | 1.21e-14 | rs8176746 | UKBB |
| | TNFa | 1.96e-14 | rs8176746 | 18464913 |
| | Impedance of whole body | 6.25e-13 | rs8176746 | UKBB |
| | Angiotensin convertin enzyme ACE activity | 2.92e-10 | rs8176746 | 20066004 |
| | Low-density lipoprotein | 2.29e-08 | rs8176746 | 24097068 |

**Table S3. Continues.**

| | Trait | P-value | SNP | PMID |
|---|---|---|---|---|
| **sVAM-1 in NFBC1966; *ABO*; 9q34.2; 9:136131322; rs8176746 (continues)** | | | | |
| GWAS Catalog (inquiry +-500 kb around the lead SNP) | ADAMTS13 levels | 1.00e-57 | rs28673647 | 29296746, 25934476 |
| | Angiotensin-converting enzyme activity | 3.00e-08 | rs495828 | 20066004 |
| | Basophil percentage of white cells + multiple other white blood cell-related traits | 5.00e-25 | rs550065584 | 27863252 |
| | Blood protein levels (E-selectin) | 1.00e-96 | rs651007 | 28240269, 19729612 |
| | Cholesterol, total | 4.00e-49 | rs579459 | 28334899, 24097068, 20686565, 25961943 |
| | Coagulation factor levels | 2.00e-138 | rs687289 | 23267103 |
| | Coronary artery disease or stroke | 4.00e-14 | rs579459 | 21378990, 29212778, 24262325, 27997041, 29531354 |
| | Diastolic blood pressure | 2.00e-18 | rs6271 | 27618452 |
| | Elevated serum carcinoembryonic antigen levels | 2.00e-24 | rs8176741 | 24941225 |
| | End-stage coagulation | 2.00e-25 | rs651007 | 23381943 |
| | Endothelial growth factor levels | 2.00e-33 | rs8176693 | 25552591 |
| | GIP levels in response to oral glucose tolerance test (120 min) | 7.00e-14 | rs635634 | 29093273 |
| | Hemoglobin + other corpuscular-related traits | 2.00e-25 | rs10901252 | 27863252, 29403010, 20139978, 28017375 |
| | High serum lipase activity | 1.00e-22 | rs8176693 | 25028398 |
| | Intraocular pressure | 3.00e-11 | rs8176743 | 25173106, 28073927, 29617998 |
| | Iron status biomarkers (ferritin levels) | 1.00e-08 | rs651007 | 25352340 |
| | Lactate dehydrogenase levels | 7.00e-09 | rs116552240 | 29403010 |
| | LDL cholesterol | 2.00e-51 | rs579459 | 28334899, 24097068, 20686565, 25961943 |
| | Liver enzyme levels | 3.00e-123 | rs579459 | 22001757 |
| | Malaria | 4.00e-21 | rs8176719 | 22895189 |
| | Serum alkaline phosphatase levels | 1.00e-56 | rs651007 | 24094242 |
| | Smoking behavior | 4.00e-08 | rs3025343 | 20418890 |
| | Soluble levels of adhesion molecules (P-selectin, ICAM) | 2.00e-41 | rs579459 | 20167578 |
| | Tonsillectomy | 9.00e-09 | rs635634 | 27182965, 28928442 |
| | Tumor biomarkers (CEA) | 7.00e-105 | rs8176749 | 23300138 |
| | Type 2 diabetes | 2.00e-08 | rs635634 | 28566273 |
| | Urinary metabolites (H-NMR features) | 1.00e-28 | rs579459 | 24586186 |
| | Venous thromboembolism | 4.00e-21 | rs8176645 | 28373160, 22672568 |
| **sVAM-1 in NFBC1966; *HPS90B1*; 12q23.3; 12:104448391; rs117238625** | | | | |
| Biobank Engine (inquiry the lead SNP) | na | | | |
| Phenoscanner (inquiry the lead SNP) | na | | | |
| GWAS Catalog (inquiry +-500 kb around the lead SNP) | Adolescent idiopathic scoliosis | 4.00e-08 | rs7138824 | 30019117 |
| | Blood protein levels (Endoplasmin. HSP90B1.6393.63.3) | 8.00e-133 | rs63658260 | 29875488 |
| | Central corneal thickness/structure | 2.00e-17 | rs11553764 | 29760442, 23291589 |
| | Intraocular pressure | 4.00e-13 | rs11553764 | 29617998 |
| | vWF level | 7.00e-10 | rs4981022 | 20231535, 26486471 |
| | Stem cell growth factor beta levels | 1.00e-23 | rs187503377 | 27989323 |

**Table S3. Continues.**

| | Trait | P-value | SNP | PMID |
|---|---|---|---|---|
| **sVAM-1 in NFBC1966; *ABCA8*; 17q24.2; 17:66823805; rs112001035** | | | | |
| Biobank Engine (inquiry the lead SNP) | na | | | |
| Phenoscanner (inquiry the lead SNP) | na | | | |
| GWAS Catalog (inquiry +-500 kb around the lead SNP) | Adolescent idiopathic scoliosis | 7.00e-10 | rs9893173 | 30019117 |
| | Blood protein levels (C-C motif chemokine 22. CCL22.3508.78.3 + others) | 7.00e-22 | rs77542162 | 29875488 |
| | HDL cholesterol | 2.00e-15 | rs1373067 | 28334899, 24097068, 20686565 |
| | Hemoglobin concentration + other corpuscular-related traits | 2.00e-16 | rs77542162 | 27863252 |
| | Intraocular pressure | 8.00e-21 | rs77542162 | 29617998 |
| | LDL cholesterol | 2.00e-18 | rs77542162 | 25961943 |
| | Musician's dystonia | 4.00e-09 | rs11655081 | 24375517 |
| | Phosphorus levels | 1.00e-08 | rs35186465 | 29403010 |
| | Total cholesterol | 2.00e-13 | rs77542162 | 25961943 |
| **IL1b in meta-analysis; HLA locus; 6p22.1; 6:30017071; rs6917603** | | | | |
| Biobank Engine (inquiry the lead SNP) | na | | | |
| Phenoscanner (inquiry the lead SNP) | Lipid metabolism phenotypes | 3.00e-29 | rs6917603 | 22286219 |
| | Gender | 6.80e-20 | rs6917603 | 22362730 |
| | Rheumatoid arthrtitis | 2.80e-13 | rs6917603 | 24390342 |
| GWAS Catalog (inquiry +-500 kb around the lead SNP) | Albumin-globulin ratio | 3.00e-11 | rs114180431 | 29403010 |
| | Allergic disease (asthma. hay fever or eczema) | 2.00e-09 | rs9259819 | 29083406 |
| | Autism spectrum disorder or schizophrenia | 1.00e-22 | rs151267808 | 28540026 |
| | Beta-2 microglobulin plasma levels | 2.00e-22 | rs9260489 | 23417110 |
| | Bipolar disorder lithium response or schizophrenia | 1.00e-17 | rs144373461 | 29121268 |
| | Blood protein levels | 4.00e-22 | rs3132645 | 28240269, 29875488 |
| | Chickenpox | 1.00e-10 | rs10947050 | 28928442 |
| | Colonoscopy-negative controls vs population controls | 2.00e-08 | rs29234 | 29228715 |
| | Crohn's disease | 2.00e-10 | rs9258260 | 22412388 |
| | Cutaneous lupus erythematosus | 2.00e-11 | rs3094067 | 25827949 |
| | Depression, depressive symptoms | 5.00e-13 | rs2517601 | 29662059 |
| | Drug-induced liver injury | 2.00e-10 | rs2523822 | 21570397, 28043905 |
| | General cognitive ability | 1.00e-11 | rs885916 | 29844566 |
| | Graves' disease | 2.00e-20 | rs3893464 | 21900946 |
| | Hand grip strength | 2.00e-08 | rs453325 | 29691431 |
| | Heel bone mineral density | 3.00e-11 | rs9260426 | 28869591 |
| | Height | 2.00e-24 | rs11970475 | 25429064 |
| | Hepatocellular carcinoma in hepatitis B infection | 9.00e-13 | rs1110446 | 29784950 |
| | HIV-1 control | 2.00e-08 | rs7758512 | 20041166 |
| | IgA nephropathy | 2.00e-11 | rs2523946 | 22197929, 26028593 |
| | IgE levels | 1.00e-15 | rs2571391 | 22075330 |

**Table S3. Continues.**

| | Trait | P-value | SNP | PMID |
|---|---|---|---|---|
| **IL1b in meta-analysis; HLA locus; 6p22.1; 6:30017071; rs6917603 (continues)** | | | | |
| GWAS Catalog (continues) | Itch intensity from mosquito bite | 1.00e-10 | rs2516703 | 28199695 |
| | Late-onset myasthenia gravis | 1.00e-11 | rs150881176 | 26562150 |
| | Lipid metabolism phenotypes | 3.00e-29 | rs6917603 | 22286219 |
| | Lung cancer | 4.00e-18 | rs114601353 | 28604730 |
| | Menarche (age at onset) | 3.00e-10 | rs16896742 | 25231870 |
| | Mixed cellularity Hodgkin lymphoma | 3.00e-23 | rs1633096 | 29196614 |
| | Moderate to vigorous physical activity levels | 1.00e-09 | rs3094622 | 29899525 |
| | Multiple sclerosis | 1.00e-17 | rs2523393 | 19525953 |
| | Multiple white or red blood cell-related traits | 2.00e-74 | rs9260620 | 27863252 |
| | Mumps | 2.00e-17 | rs114193679 | 28928442 |
| | Nasopharyngeal carcinoma | 9.00e-17 | rs29232 | 19664746 |
| | Neuroticism | 1.00e-09 | rs2267633 | 29255261 |
| | Non-albumin protein levels | 3.00e-09 | rs142014154 | 29403010 |
| | Prostate cancer | 2.00e-08 | rs115457135 | 25217961, 29892016 |
| | Sarcoidosis (Lofgren's syndrome vs non-Lofgren's syndrome) | 4.00e-23 | rs3130141 | 26651848 |
| | Schizophrenia | 2.00e-19 | rs114200269 | 26198764, 27922604, 22883433, 21926974 |
| | Shingles | 2.00e-22 | rs2523815 | 28928442 |
| | Tonsillectomy | 8.00e-10 | rs10484552 | 28928442 |
| | Tuberculosis | 2.00e-12 | rs181383036 | 28928442 |
| | Type 1 diabetes and autoimmune thyroid diseases | 2.00e-08 | rs2523989 | 25936594 |
| | Vitiligo | 2.00e-66 | rs60131261 | 27723757 |
| **TNFa in meta-analyses; *DLEU1*; 13q14.3; 13:51141997; rs17074575** | | | | |
| Biobank Engine (inquiry the lead SNP) | na | | | |
| Phenoscanner (inquiry the lead SNP) | na | | | |
| GWAS Catalog (inquiry +-500 kb around the lead SNP) | Adolescent idiopathic scoliosis | 1.00e-08 | rs797487 | 30019117 |
| | Aortic root size | 2.00e-15 | rs2762049 | 28394258 |
| | Circulating odd-numbered chain saturated fatty acid levels (C19:0) | 7.00e-09 | rs17363566 | 29738550 |
| | Eosinophil percentage of white cells | 5.00e-10 | rs201798 | 27863252 |
| | Heel bone mineral density | 4.00e-10 | rs1149821 | 28869591 |
| | Height | 1.00e-69 | rs3118905 | 25282103 |
| | Hemoglobin | 2.00e-10 | rs806293 | 29403010 |
| | High light scatter reticulocyte percentage of red cells | 1.00e-13 | rs9535495 | 27863252 |
| | Hip circumference adjusted for BMI | 4.00e-15 | rs558003 | 25673412 |
| | Multiple sclerosis | 3.00e-10 | rs2812197 | 27386562 |
| | Primary biliary cholangitis | 1.00e-10 | rs9591325 | 26394269 |
| | Primary tooth development (number of teeth) | 3.00e-08 | rs9316505 | 23704328 |
| | Prostate-specific antigen levels | 3.00e-22 | rs202346 | 28139693 |
| | Reticulocyte fraction of red cells | 3.00e-11 | rs9535495 | 27863252 |

**Table S4. The results of the supplemental GWAS.**

Individuals who had reported having fever at the time of the blood sampling (N=32) or C-reactive protein (CRP) > 10 mg/l (N=212) were excluded from the supplementary GWAS completed in NFBC1966. The data adjustments (age, sex, BMI, and the first ten genetic principal components) and transformations were performed as in the primary analyses. The results are given for the locus lead SNPs as in the Table 2.

| Marker | Locus | Chr:Position | Candidate gene | Nearest gene(s) | Annotation | dbSNP refrerence | EA | EAF | Beta | P-value | $Beta_{origNFBC1966}$ | $P\text{-}value_{origNFBC1966}$ |
|---|---|---|---|---|---|---|---|---|---|---|---|---|
| sE-Selectin | 9q34.2 | 9:136141870 | *ABO* | *ABO* | Intronic | rs2519093 | T | 0.212 | -0.908 | 8.01E-295 | -0.903 | 4.48E-305 |
| | 11q24.2 | 11:126266665 | *ST3GAL4* | *ST3GAL4* | Intronic | rs11220471 | G | 0.241 | -0.157 | 8.76E-11 | -0.162 | 7.72E-12 |
| sICAM-1 | 9q34.2 | 9:136141870 | *ABO* | *ABO* | Intronic | rs2519093 | T | 0.212 | -0.350 | 2.26E-45 | -0.352 | 7.43E-48 |
| | 19p13.2 | 19:10383403 | *ICAM1* | *ICAM1* | Intronic | rs117960796 | A | 0.012 | -1.769 | 8.75E-41 | -1.669 | 8.03E-40 |
| sVCAM-1 | 9q34.2 | 9:136131322 | *ABO* | *ABO* | Missense | rs8176746 | T | 0.138 | 0.257 | 2.38E-18 | 0.256 | 5.06E-19 |
| | 12q23.3 | 12:104448391 | *HSP90B1* | *GLT8D2* | Intronic | rs117238625 | A | 0.023 | 0.527 | 2.28E-14 | 0.510 | 2.90E-14 |
| | 17q24.2 | 17:66823805 | *ABCA8* | *ABCA8* | Intergenic | rs112001035 | A | 0.062 | -0.315 | 1.88E-12 | -0.324 | 1.04E-13 |
| IL1β | 6p22.1 | 6:30017071 | | *HLA* locus | Intronic | rs6917603 | C | 0.291 | -0.164 | 2.28E-12 | -0.163 | 8.82E-13 |
| IP10 | 4q21.1 | 4:76899176 | *CXCL10* | *SAD1* | Intronic | rs192716315 | C | 0.003 | 1.472 | 6.35E-13 | 1.513 | 1.79E-13 |
| MCP1 | 1q23.2 | 1:159175354 | *ACKR1* | *ACKR1* | Missense | rs12075 | G | 0.507 | 0.028 | 0.174 | 0.031 | 0.123 |
| TNFα | 13q14.3 | 13:51141997 | *DLEU1* | *DLEU1* | Intronic | rs17074575 | G | 0.001 | 2.560 | 6.40E-10 | 2.131 | 2.48E-09 |
| VEGF | 4p16.2 | 4:5636073 | *STK32B* | *EVC2* | Intronic | rs186725382 | A | 0.001 | -2.629 | 9.13E-11 | -2.380 | 4.22E-10 |
| | 6p21.1 | 6:43927050 | *VEGFA* | *C6orf223* | Intergenic | rs7767396 | G | 0.511 | -0.015 | 0.452 | -0.015 | 0.465 |
| | 9p24.2 | 9:2686273 | *VLDLR* | *VLDLR, KCNV2* | Intergenic | rs7030781 | T | 0.392 | 0.036 | 0.097 | 0.042 | 0.044 |

Statistical significance is considered at $P<3.1x10^{-9}$. All positions correspond to human genome build 37. $Beta_{origNFBC1966}$ and $P\text{-}value_{origNFBC1966}$ indicate the test statistics as in the original discovery GWAS in the NFBC1966 population. EA, effect allele; EAF, effect allele frequency.

**Table S5. The list of SNPs used to determine the correlations of the SNP effects on inflammatory phenotypes versus other traits.**

| phenotype | locus | SNP | EA | Z_phenotype | P_phenotype | Z_cad | P_cad | Z_stroke | P_stroke | Z_ldlc | P_ldlc | Z_hdlc | P_hdlc |
|---|---|---|---|---|---|---|---|---|---|---|---|---|---|
| sEselectin | ABO | rs17408598 | C | -5.09 | 3.49E-07 | -1.142 | 0.254 | 0.356 | 0.721 | 1.541 | 0.129 | 1.044 | 0.301 |
| sEselectin | ABO | rs13295430 | A | -6.91 | 4.78E-12 | 0.667 | 0.505 | 0.944 | 0.345 | -0.732 | 0.471 | -0.019 | 0.985 |
| sEselectin | ABO | rs11243956 | T | 4.12 | 3.79E-05 | -0.804 | 0.422 | -1.000 | 0.319 | -0.741 | 0.466 | -0.879 | 0.384 |
| sEselectin | ABO | rs3011259 | T | 4.39 | 1.12E-05 | -0.742 | 0.458 | -0.406 | 0.686 | -1.288 | 0.205 | -0.288 | 0.776 |
| sEselectin | ABO | rs8193004 | T | -4.13 | 3.57E-05 | -1.419 | 0.156 | -0.646 | 0.518 | -0.123 | 0.904 | -1.866 | 0.064 |
| sEselectin | ABO | rs628094 | A | -4.86 | 1.19E-06 | 1.586 | 0.113 | 0.813 | 0.417 | 2.125 | 0.036 | 1.225 | 0.225 |
| sEselectin | ABO | rs7039251 | A | -5.15 | 2.66E-07 | -0.669 | 0.504 | -0.071 | 0.945 | 1.749 | 0.085 | -0.984 | 0.329 |
| sEselectin | ABO | rs2073821 | T | 4.91 | 9.14E-07 | -0.776 | 0.438 | 1.231 | 0.219 | -1.688 | 0.097 | 1.164 | 0.249 |
| sEselectin | ABO | rs2519090 | A | 3.92 | 8.78E-05 | -0.330 | 0.742 | -0.650 | 0.516 | -0.580 | 0.568 | -0.787 | 0.436 |
| sEselectin | ABO | rs598499 | G | 9.96 | 2.18E-23 | -0.102 | 0.918 | 0.478 | 0.633 | -2.256 | 0.026 | 0.302 | 0.765 |
| sEselectin | ABO | rs671050 | C | -5.71 | 1.15E-08 | -0.944 | 0.345 | 1.085 | 0.279 | 0.873 | 0.390 | 0.348 | 0.730 |
| sEselectin | ABO | rs615940 | C | 4.92 | 8.57E-07 | 0.303 | 0.762 | 1.159 | 0.246 | -2.385 | 0.019 | -1.479 | 0.143 |
| sEselectin | ABO | rs2073927 | G | -5.24 | 1.57E-07 | 0.538 | 0.591 | -1.970 | 0.049 | 0.418 | 0.680 | -1.313 | 0.193 |
| sEselectin | ABO | rs11244015 | C | -8.51 | 1.80E-17 | 0.687 | 0.492 | -1.031 | 0.303 | 1.837 | 0.071 | 0.645 | 0.523 |
| sEselectin | ABO | rs7868232 | C | -13.77 | 3.81E-43 | 3.810 | 0.0001 | 1.043 | 0.296 | 1.731 | 0.088 | 1.439 | 0.154 |
| sEselectin | ABO | rs7039497 | A | 5.83 | 5.64E-09 | -1.766 | 0.077 | -0.575 | 0.566 | -1.642 | 0.106 | -0.758 | 0.453 |
| sEselectin | ABO | rs10901242 | T | 7.63 | 2.43E-14 | -3.015 | 0.003 | -2.120 | 0.034 | -0.898 | 0.377 | 0.114 | 0.910 |
| sEselectin | ABO | rs7036819 | T | 4.30 | 1.72E-05 | -0.361 | 0.718 | 0.379 | 0.705 | 0.885 | 0.384 | -1.152 | 0.254 |
| sEselectin | ABO | rs4489379 | T | -6.93 | 4.20E-12 | 1.398 | 0.162 | 0.690 | 0.490 | 0.108 | 0.916 | -0.479 | 0.635 |
| sEselectin | ABO | rs7036324 | A | 7.75 | 9.48E-15 | 0.491 | 0.624 | -0.402 | 0.688 | 0.054 | 0.957 | -1.383 | 0.170 |
| sEselectin | ABO | rs10901250 | A | 6.50 | 7.82E-11 | -0.222 | 0.824 | 0.711 | 0.476 | -2.050 | 0.044 | -1.163 | 0.249 |
| sEselectin | ABO | rs8176707 | T | 7.78 | 7.13E-15 | -0.758 | 0.448 | 1.407 | 0.160 | -1.968 | 0.053 | -0.695 | 0.491 |
| sEselectin | ABO | rs2073827 | C | 17.93 | 6.44E-72 | -2.488 | 0.013 | -2.072 | 0.039 | -2.392 | 0.019 | -0.513 | 0.612 |
| sEselectin | ABO | rs630014 | G | -16.10 | 2.71E-58 | 4.079 | 4.51E-05 | 3.076 | 0.002 | 0.775 | 0.445 | -0.690 | 0.494 |
| sEselectin | ABO | rs651007 | T | -35.34 | 1.71E-273 | 6.441 | 1.19E-10 | 3.314 | 0.001 | 5.478 | 7.14E-08 | 0.749 | 0.458 |
| sEselectin | ABO | rs633862 | T | -18.82 | 4.76E-79 | 2.695 | 0.007 | 1.616 | 0.106 | 3.944 | 0.0001 | 0.386 | 0.702 |
| sEselectin | ABO | rs176690 | T | -7.29 | 3.12E-13 | -0.001 | 0.999 | -0.656 | 0.512 | 2.161 | 0.033 | 1.565 | 0.121 |
| sEselectin | ABO | rs17150482 | C | 13.27 | 3.39E-40 | -0.688 | 0.492 | 0.797 | 0.424 | -2.670 | 0.009 | -0.651 | 0.519 |
| sEselectin | ABO | rs3124747 | G | 13.95 | 3.20E-44 | -2.048 | 0.041 | 0.951 | 0.342 | -1.545 | 0.128 | -1.375 | 0.173 |
| sEselectin | ABO | rs685523 | T | 7.72 | 1.13E-14 | -2.612 | 0.009 | -0.959 | 0.337 | -1.210 | 0.234 | -0.443 | 0.660 |
| sEselectin | ABO | rs652600 | A | -5.92 | 3.32E-09 | 1.200 | 0.230 | 1.669 | 0.095 | 0.656 | 0.518 | -0.319 | 0.752 |
| sEselectin | ABO | rs2301614 | A | 7.02 | 2.17E-12 | -0.305 | 0.760 | 1.633 | 0.102 | -0.792 | 0.436 | 0.038 | 0.970 |
| sEselectin | ABO | rs3025301 | G | 4.26 | 2.09E-05 | -0.316 | 0.752 | -0.309 | 0.759 | -0.513 | 0.613 | -2.402 | 0.017 |
| sEselectin | ABO | rs10993874 | T | -5.59 | 2.32E-08 | 1.581 | 0.114 | 0.133 | 0.894 | -0.738 | 0.468 | 2.097 | 0.038 |
| sEselectin | ABO | rs1541332 | A | -6.68 | 2.32E-11 | 2.243 | 0.025 | -0.348 | 0.727 | 1.130 | 0.266 | 0.544 | 0.590 |
| sEselectin | ABO | rs2283124 | T | -4.49 | 7.26E-06 | -0.610 | 0.542 | -1.429 | 0.154 | 1.068 | 0.293 | -0.750 | 0.458 |
| sEselectin | ABO | rs129932 | A | 5.25 | 1.56E-07 | 1.894 | 0.058 | 1.000 | 0.317 | -0.342 | 0.736 | 0.242 | 0.811 |
| sEselectin | ABO | rs129940 | G | -7.10 | 1.25E-12 | -0.121 | 0.904 | 0.954 | 0.340 | 0.430 | 0.672 | 0.638 | 0.527 |
| sEselectin | ABO | rs622035 | C | -3.94 | 8.16E-05 | -0.216 | 0.829 | -0.100 | 0.920 | -0.589 | 0.562 | 0.255 | 0.801 |
| sEselectin | ABO | rs669443 | T | 4.98 | 6.27E-07 | -0.341 | 0.733 | 0.835 | 0.402 | -1.163 | 0.252 | -2.380 | 0.018 |

**Table S5. (continues)**

| phenotype | locus | SNP | EA | Z_phenotype | P_phenotype | Z_cad | P_cad | Z_stroke | P_stroke | Z_ldlc | P_ldlc | Z_hdlc | P_hdlc |
|---|---|---|---|---|---|---|---|---|---|---|---|---|---|
| sICAM1 | ABO | rs10901243 | C | -6.21 | 5.38E-10 | 3.412 | 0.001 | 1.196 | 0.233 | 1.021 | 0.315 | 0.541 | 0.592 |
| sICAM1 | ABO | rs2073827 | C | 7.55 | 4.42E-14 | -2.488 | 0.013 | -2.072 | 0.039 | -2.392 | 0.019 | -0.513 | 0.612 |
| sICAM1 | ABO | rs630014 | G | -6.47 | 9.57E-11 | 4.079 | 4.51E-05 | 3.076 | 0.002 | 0.775 | 0.445 | -0.690 | 0.494 |
| sICAM1 | ABO | rs651007 | T | -13.28 | 3.22E-40 | 6.441 | 1.19E-10 | 3.314 | 0.001 | 5.478 | 7.14E-08 | 0.749 | 0.458 |
| sICAM1 | ABO | rs633862 | T | -7.49 | 6.70E-14 | 2.695 | 0.007 | 1.616 | 0.106 | 3.944 | 0.0001 | 0.386 | 0.702 |
| sICAM1 | ABO | rs17150482 | C | 5.27 | 1.40E-07 | -0.688 | 0.492 | 0.797 | 0.424 | -2.670 | 0.009 | -0.651 | 0.519 |
| sICAM1 | ABO | rs3124747 | G | 4.31 | 1.62E-05 | -2.048 | 0.041 | 0.951 | 0.342 | -1.545 | 0.128 | -1.375 | 0.173 |
| sICAM1 | ABO | rs652600 | A | -4.69 | 2.68E-06 | 1.200 | 0.230 | 1.669 | 0.095 | 0.656 | 0.518 | -0.319 | 0.752 |
| sVCAM1 | ABO | rs2905078 | A | 4.32 | 1.57E-05 | -0.083 | 0.933 | -0.126 | 0.900 | -0.172 | 0.866 | -0.921 | 0.361 |
| sVCAM1 | ABO | rs4962099 | C | 4.81 | 1.49E-06 | -1.040 | 0.299 | 0.840 | 0.401 | -1.698 | 0.095 | -0.751 | 0.457 |
| sVCAM1 | ABO | rs4489379 | T | -4.88 | 1.07E-06 | 1.398 | 0.162 | 0.690 | 0.490 | 0.108 | 0.916 | -0.479 | 0.635 |
| sVCAM1 | ABO | rs8176746 | T | 8.91 | 5.06E-19 | -1.189 | 0.235 | 1.554 | 0.121 | -2.091 | 0.040 | -0.252 | 0.803 |
| sVCAM1 | ABO | rs579459 | C | -6.16 | 7.26E-10 | 6.447 | 1.14E-10 | 3.828 | 0.0001 | 5.496 | 6.45E-08 | 0.760 | 0.451 |
| sVCAM1 | HSP90B1 | rs4981022 | A | 4.71 | 2.43E-06 | -0.535 | 0.593 | 1.429 | 0.154 | 1.004 | 0.323 | -0.841 | 0.404 |
| sVCAM1 | HSP90B1 | rs2271637 | G | 4.25 | 2.10E-05 | 1.334 | 0.182 | 1.878 | 0.061 | -0.389 | 0.702 | -0.935 | 0.354 |
| sVCAM1 | HSP90B1 | rs4135064 | T | 4.37 | 1.26E-05 | 0.195 | 0.845 | 0.891 | 0.372 | 1.366 | 0.179 | 0.296 | 0.769 |
| sVCAM1 | HSP90B1 | rs1564892 | G | 4.72 | 2.35E-06 | -0.592 | 0.554 | 0.546 | 0.586 | 0.044 | 0.966 | -0.527 | 0.601 |
| sVCAM1 | HSP90B1 | rs6539137 | T | -4.66 | 3.18E-06 | 1.813 | 0.070 | -1.593 | 0.111 | -0.369 | 0.717 | 0.129 | 0.899 |
| sVCAM1 | HSP90B1 | rs1922270 | C | 4.19 | 2.79E-05 | -0.905 | 0.366 | 0.286 | 0.775 | 0.691 | 0.496 | -1.070 | 0.289 |
| sEselectin | ST3GAL4 | rs11220286 | C | 4.02 | 5.73E-05 | 0.832 | 0.406 | 0.727 | 0.467 | 1.464 | 0.150 | 0.492 | 0.626 |
| sEselectin | ST3GAL4 | rs8177399 | T | 4.41 | 1.05E-05 | 2.614 | 0.009 | 0.994 | 0.321 | 2.599 | 0.011 | -0.168 | 0.868 |
| sEselectin | ST3GAL4 | rs1786698 | C | -4.22 | 2.42E-05 | -0.207 | 0.836 | 0.733 | 0.464 | -0.965 | 0.342 | 0.360 | 0.722 |
| sEselectin | ST3GAL4 | rs3862629 | T | 4.20 | 2.68E-05 | 1.499 | 0.134 | -0.879 | 0.379 | 1.102 | 0.278 | -1.513 | 0.134 |
| sEselectin | ST3GAL4 | rs10893505 | A | 4.79 | 1.68E-06 | 2.317 | 0.021 | -2.112 | 0.035 | 1.914 | 0.060 | -0.709 | 0.482 |
| sEselectin | ST3GAL4 | rs7933887 | A | -6.63 | 3.30E-11 | -2.677 | 0.007 | -1.121 | 0.262 | -0.540 | 0.595 | 1.929 | 0.056 |
| sEselectin | ST3GAL4 | rs3802815 | C | -4.08 | 4.58E-05 | -1.399 | 0.162 | -0.452 | 0.653 | -1.178 | 0.246 | -0.290 | 0.774 |
| sEselectin | ST3GAL4 | rs4935969 | T | 5.29 | 1.23E-07 | 0.358 | 0.720 | -0.274 | 0.782 | 0.956 | 0.347 | 0.564 | 0.576 |
| sEselectin | ST3GAL4 | rs11220490 | T | 6.54 | 6.16E-11 | 1.144 | 0.253 | -0.207 | 0.836 | 0.460 | 0.651 | -1.502 | 0.137 |
| sEselectin | ST3GAL4 | rs11605349 | C | 4.89 | 9.95E-07 | 1.368 | 0.171 | -0.270 | 0.787 | 0.990 | 0.330 | -1.117 | 0.269 |

**Table S5. (continues)**

| phenotype | locus | SNP | EA | Z_phenotype | P_phenotype | Z_cad | P_cad | Z_stroke | P_stroke | Z_ldlc | P_ldlc | Z_hdlc | P_hdlc |
|---|---|---|---|---|---|---|---|---|---|---|---|---|---|
| sICAM1 | ICAM1 | rs2303099 | T | 4.69 | 2.73E-06 | -1.292 | 0.197 | -1.240 | 0.216 | 3.324 | 0.001 | 1.051 | 0.298 |
| sICAM1 | ICAM1 | rs16995016 | A | 3.92 | 8.76E-05 | -0.782 | 0.434 | 1.516 | 0.131 | -0.753 | 0.458 | -2.446 | 0.015 |
| sICAM1 | ICAM1 | rs8109578 | A | -4.25 | 2.18E-05 | 1.122 | 0.262 | 1.375 | 0.170 | -2.206 | 0.030 | -1.258 | 0.213 |
| sICAM1 | ICAM1 | rs11085721 | C | 4.06 | 4.81E-05 | -0.399 | 0.690 | -1.120 | 0.262 | 0.609 | 0.549 | 0.019 | 0.985 |
| sICAM1 | ICAM1 | rs1988068 | A | -4.26 | 2.09E-05 | 1.381 | 0.167 | 2.176 | 0.030 | 0.262 | 0.796 | -0.762 | 0.450 |
| sICAM1 | ICAM1 | rs8111930 | G | -7.43 | 1.07E-13 | 1.221 | 0.222 | -0.416 | 0.678 | -1.469 | 0.148 | 0.635 | 0.529 |
| sICAM1 | ICAM1 | rs5030390 | A | -6.15 | 7.88E-10 | -1.326 | 0.185 | -0.297 | 0.766 | -0.307 | 0.763 | 0.385 | 0.703 |
| sICAM1 | ICAM1 | rs1799969 | A | -6.70 | 2.07E-11 | 1.136 | 0.256 | 0.306 | 0.760 | -2.048 | 0.044 | -1.964 | 0.052 |
| sICAM1 | ICAM1 | rs2278442 | A | 4.34 | 1.40E-05 | 0.423 | 0.672 | 0.927 | 0.354 | -0.754 | 0.458 | -0.269 | 0.790 |
| sICAM1 | ICAM1 | rs7254474 | G | 8.46 | 2.68E-17 | -0.134 | 0.893 | -0.154 | 0.879 | -0.467 | 0.645 | 0.990 | 0.327 |
| sICAM1 | ICAM1 | rs17677095 | G | 4.31 | 1.65E-05 | 1.816 | 0.069 | 1.294 | 0.195 | 2.214 | 0.029 | 1.574 | 0.119 |
| sICAM1 | ICAM1 | rs7256672 | G | -6.62 | 3.66E-11 | -0.822 | 0.411 | 0.591 | 0.554 | -4.246 | 2.96E-05 | 0.666 | 0.509 |
| sICAM1 | ICAM1 | rs10416653 | T | -6.84 | 7.96E-12 | -0.466 | 0.641 | 0.415 | 0.678 | -1.741 | 0.087 | 1.960 | 0.052 |
| sICAM1 | ICAM1 | rs9676881 | A | 4.57 | 4.84E-06 | -2.542 | 0.011 | -0.770 | 0.442 | -2.450 | 0.016 | 1.288 | 0.202 |
| sICAM1 | ICAM1 | rs10775614 | A | 3.97 | 7.10E-05 | 4.055 | 5.02E-05 | 2.018 | 0.044 | 2.000 | 0.049 | 2.758 | 0.006 |
| sICAM1 | ICAM1 | rs10418609 | T | 6.20 | 5.73E-10 | -1.961 | 0.050 | -0.599 | 0.549 | 1.246 | 0.220 | -1.527 | 0.130 |
| VEGF | VLDLR | rs1889109 | G | -4.10 | 4.25E-05 | 0.311 | 0.756 | -0.367 | 0.713 | -1.741 | 0.087 | 0.529 | 0.600 |
| VEGF | VLDLR | rs6475925 | G | 4.03 | 5.30E-05 | -0.007 | 0.995 | -2.109 | 0.035 | 0.705 | 0.488 | 0.172 | 0.864 |
| VEGF | VLDLR | rs4741769 | G | -3.91 | 8.78E-05 | -0.166 | 0.868 | 3.963 | 7.17E-05 | 0.711 | 0.484 | 0.548 | 0.587 |
| VEGF | VLDLR | rs2375980 | G | 7.12 | 9.34E-13 | -0.332 | 0.740 | -1.558 | 0.121 | -1.447 | 0.154 | 0.367 | 0.716 |
| VEGF | VLDLR | rs10968136 | C | -4.08 | 4.38E-05 | -0.295 | 0.768 | 2.802 | 0.005 | 1.639 | 0.107 | 0.225 | 0.824 |
| VEGF | VLDLR | rs10757686 | C | 4.25 | 1.94E-05 | -0.384 | 0.701 | -1.283 | 0.199 | -1.225 | 0.228 | 0.449 | 0.656 |
| VEGF | VLDLR | rs7040049 | G | -4.27 | 1.84E-05 | -1.320 | 0.187 | 0.457 | 0.647 | 2.286 | 0.024 | 0.189 | 0.852 |
| VEGF | VEGFA | rs1776721 | G | -7.83 | 4.09E-15 | -1.270 | 0.204 | 1.889 | 0.059 | -0.357 | 0.725 | 1.076 | 0.286 |
| VEGF | VEGFA | rs7767396 | G | 20.76 | 4.95E-96 | 2.079 | 0.038 | -0.294 | 0.767 | -0.651 | 0.522 | 0.463 | 0.647 |
| VEGF | VEGFA | rs17209449 | G | -6.55 | 5.91E-11 | -1.785 | 0.074 | -1.044 | 0.296 | 1.229 | 0.227 | -0.631 | 0.532 |
| VEGF | VEGFA | rs11968152 | C | 6.98 | 3.00E-12 | 0.037 | 0.970 | -0.195 | 0.845 | 0.358 | 0.725 | 0.871 | 0.388 |
| VEGF | VEGFA | rs1950506 | G | 4.53 | 5.65E-06 | -1.320 | 0.187 | -0.347 | 0.730 | -0.060 | 0.953 | 0.172 | 0.865 |
| VEGF | VEGFA | rs1359617 | C | 5.45 | 5.06E-08 | 0.144 | 0.885 | 0.724 | 0.469 | -1.702 | 0.094 | 0.513 | 0.611 |
| VEGF | VEGFA | rs17209421 | C | 4.23 | 2.39E-05 | 0.412 | 0.680 | -2.056 | 0.040 | -2.048 | 0.044 | 1.169 | 0.247 |
| VEGF | VEGFA | rs910604 | G | -5.65 | 1.81E-08 | -1.089 | 0.276 | -0.209 | 0.835 | 0.263 | 0.795 | -0.100 | 0.921 |
| VEGF | VEGFA | rs943072 | G | -6.25 | 3.75E-10 | 0.086 | 0.931 | 0.610 | 0.543 | 0.700 | 0.491 | -0.773 | 0.444 |
| VEGF | VEGFA | rs12199215 | C | 6.52 | 7.25E-11 | -0.656 | 0.512 | -0.295 | 0.770 | 0.986 | 0.332 | 0.952 | 0.345 |
| VEGF | VEGFA | rs1539508 | G | 5.37 | 7.26E-08 | -1.107 | 0.268 | -0.512 | 0.609 | -0.356 | 0.726 | -0.693 | 0.492 |
| VEGF | VEGFA | rs7757024 | C | 4.36 | 1.33E-05 | 0.520 | 0.603 | 0.753 | 0.452 | 0.955 | 0.347 | 0.079 | 0.938 |
| VEGF | VEGFA | rs7763358 | C | 5.30 | 1.19E-07 | -0.423 | 0.672 | 0.801 | 0.424 | 0.907 | 0.372 | 1.217 | 0.228 |
| VEGF | VEGFA | rs6936047 | G | -5.19 | 2.10E-07 | 0.697 | 0.486 | 1.049 | 0.295 | -0.290 | 0.775 | -0.410 | 0.685 |

### Table S5. (continues)

| phenotype | locus | SNP | EA | Z_phenotype | P_phenotype | Z_cad | P_cad | Z_stroke | P_stroke | Z_ldlc | P_ldlc | Z_hdlc | P_hdlc |
|---|---|---|---|---|---|---|---|---|---|---|---|---|---|
| MCP1 | ACKR1 | rs4269772 | C | 5.76 | 8.52E-09 | -1.052 | 0.293 | 0.166 | 0.870 | -0.723 | 0.477 | -2.232 | 0.027 |
| MCP1 | ACKR1 | rs12075 | G | 10.20 | 1.90E-24 | -1.048 | 0.295 | -1.282 | 0.200 | -0.618 | 0.543 | -2.169 | 0.032 |
| MCP1 | ACKR1 | rs12047264 | G | -4.22 | 2.34E-05 | 1.593 | 0.111 | 1.512 | 0.131 | -1.729 | 0.089 | 0.270 | 0.789 |
| MCP1 | ACKR1 | rs7540542 | G | 6.09 | 1.08E-09 | -1.129 | 0.259 | -1.245 | 0.212 | -0.917 | 0.367 | -1.240 | 0.219 |
| MCP1 | ACKR1 | rs863002 | C | 6.24 | 4.49E-10 | -0.963 | 0.336 | -0.182 | 0.854 | -0.689 | 0.498 | -1.973 | 0.051 |
| MCP1 | ACKR1 | rs3027012 | C | 4.47 | 7.37E-06 | -0.653 | 0.514 | -2.416 | 0.016 | -1.386 | 0.172 | -0.958 | 0.342 |
| MCP1 | ACKR1 | rs4656165 | G | 3.96 | 7.66E-05 | -1.337 | 0.181 | 0.805 | 0.422 | 0.464 | 0.648 | -2.440 | 0.016 |
| MCP1 | ACKR1 | rs12727764 | G | 3.99 | 6.32E-05 | -1.757 | 0.079 | -0.614 | 0.539 | 1.768 | 0.082 | -0.031 | 0.975 |

At each significant loci associating with inflammatory phenotypes in the present study, subsets of representative SNPs were extracted using clumping function in PLINK (23). SNP effects on CAD risk, stroke risk, and LDL-C and HDL-C levels were obtained from CARDIoGRAM (20), Stroke Consortia (21), and metabolomics GWAS (22) summary statistics. Prior to clumping, data were filtered to include SNPs available in all the three data sets.

**Figure S1. Correlation heatmap of the inflammatory phenotypes in NFBC1966.**

Pearson's ρ calculated for the adjusted (age, sex, BMI, ten first genetic principal components) and transformed inflammatory phenotypes. Coloring represents the sign of the correlation (red-yellow positive, purple-blue negative) and the circle area is proportional to the absolute value of the correlation. Cells with statistically insignificant values (P≥0.05) are left blank.

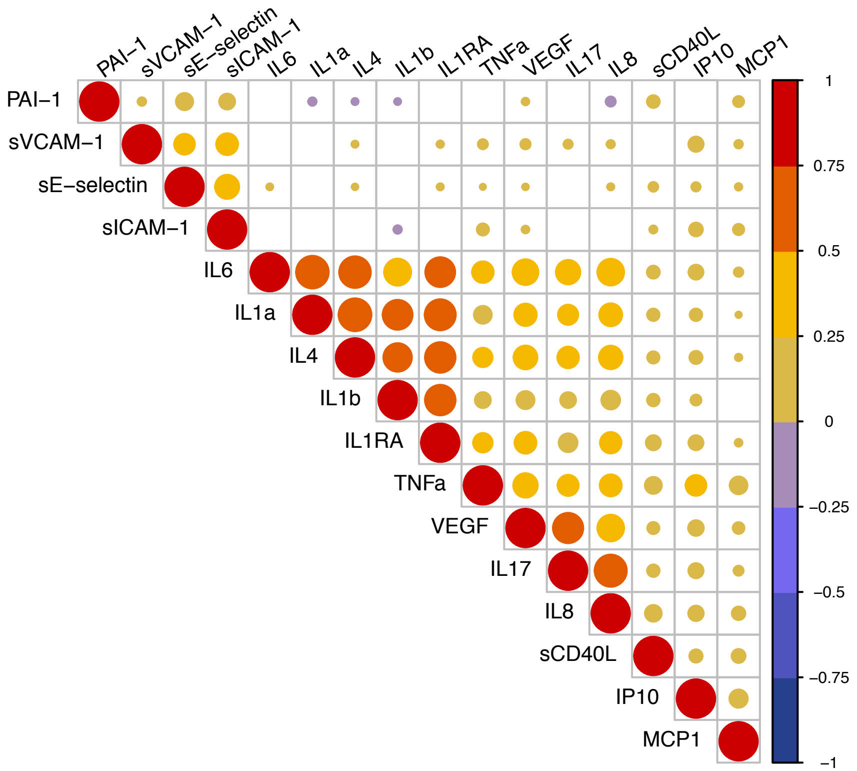

**Figure S2 A) Manhattan, Q-Q, and LocusZoom plots for sE-selectin results in NFBC1966.**

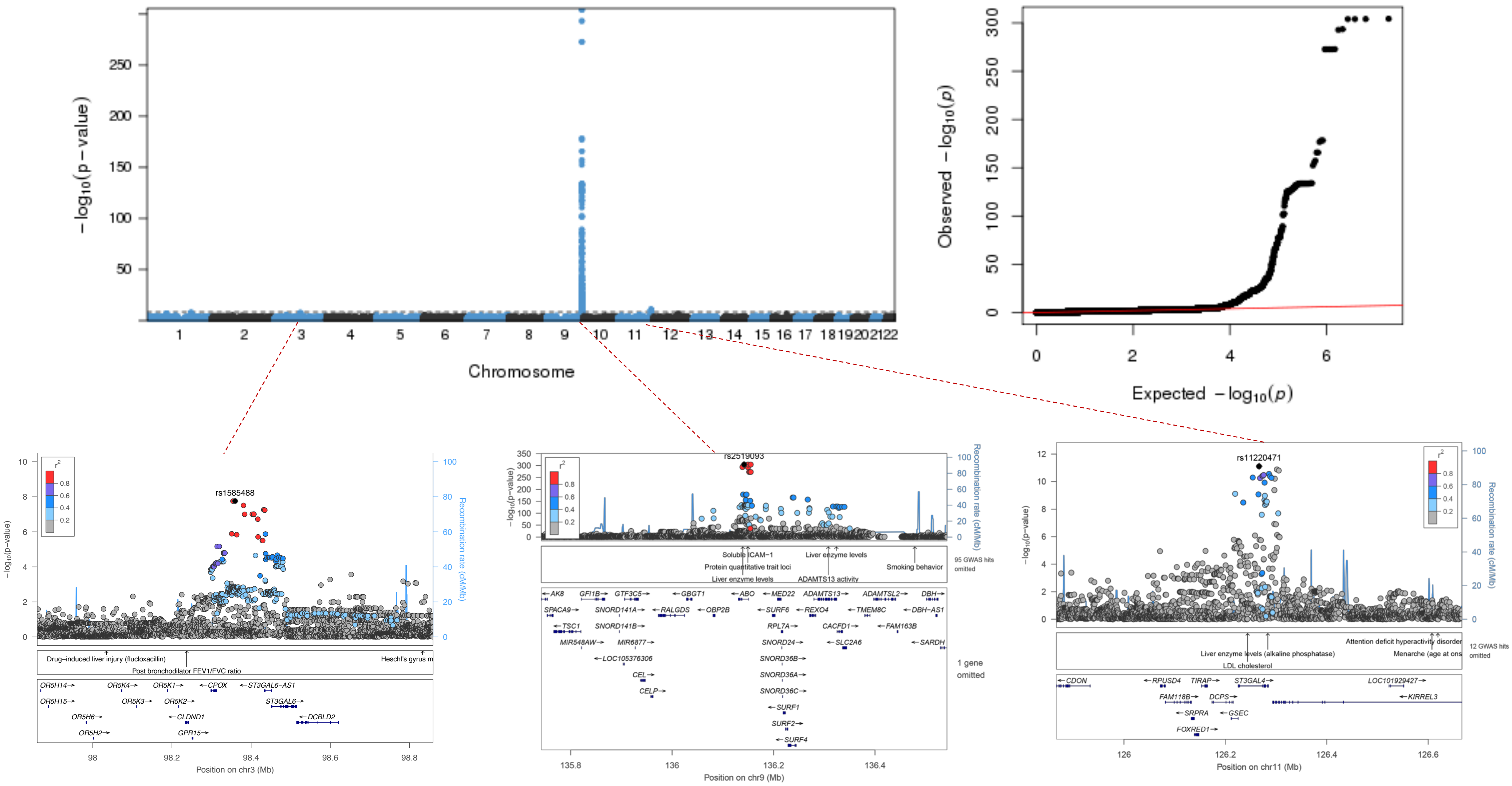

**Figure S2 B) Manhattan, Q-Q, and LocusZoom plots for sICAM-1 results in NFBC1966.**

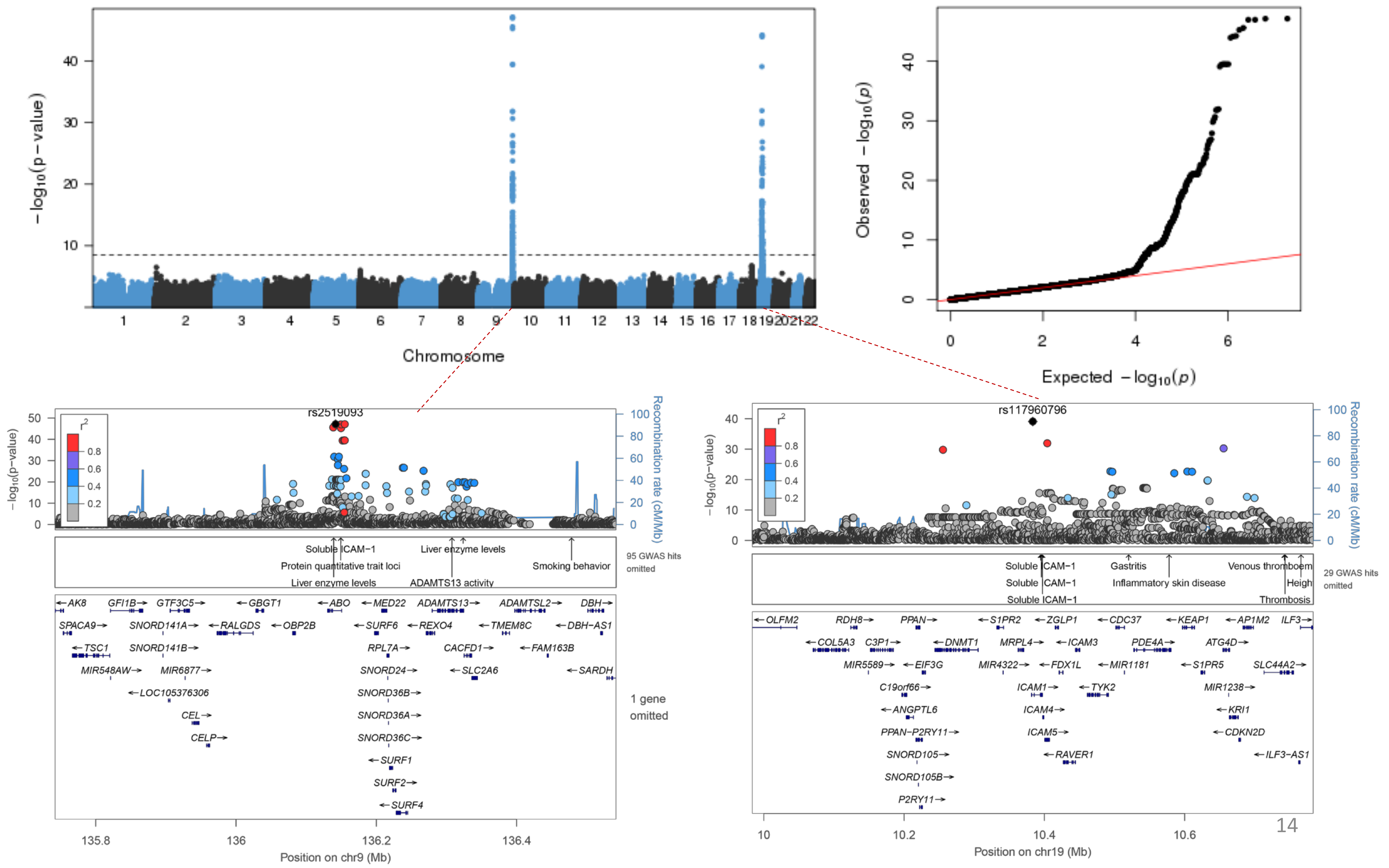

**Figure S2 C) Manhattan, Q-Q, and LocusZoom plots for sVCAM-1 results in NFBC1966.**

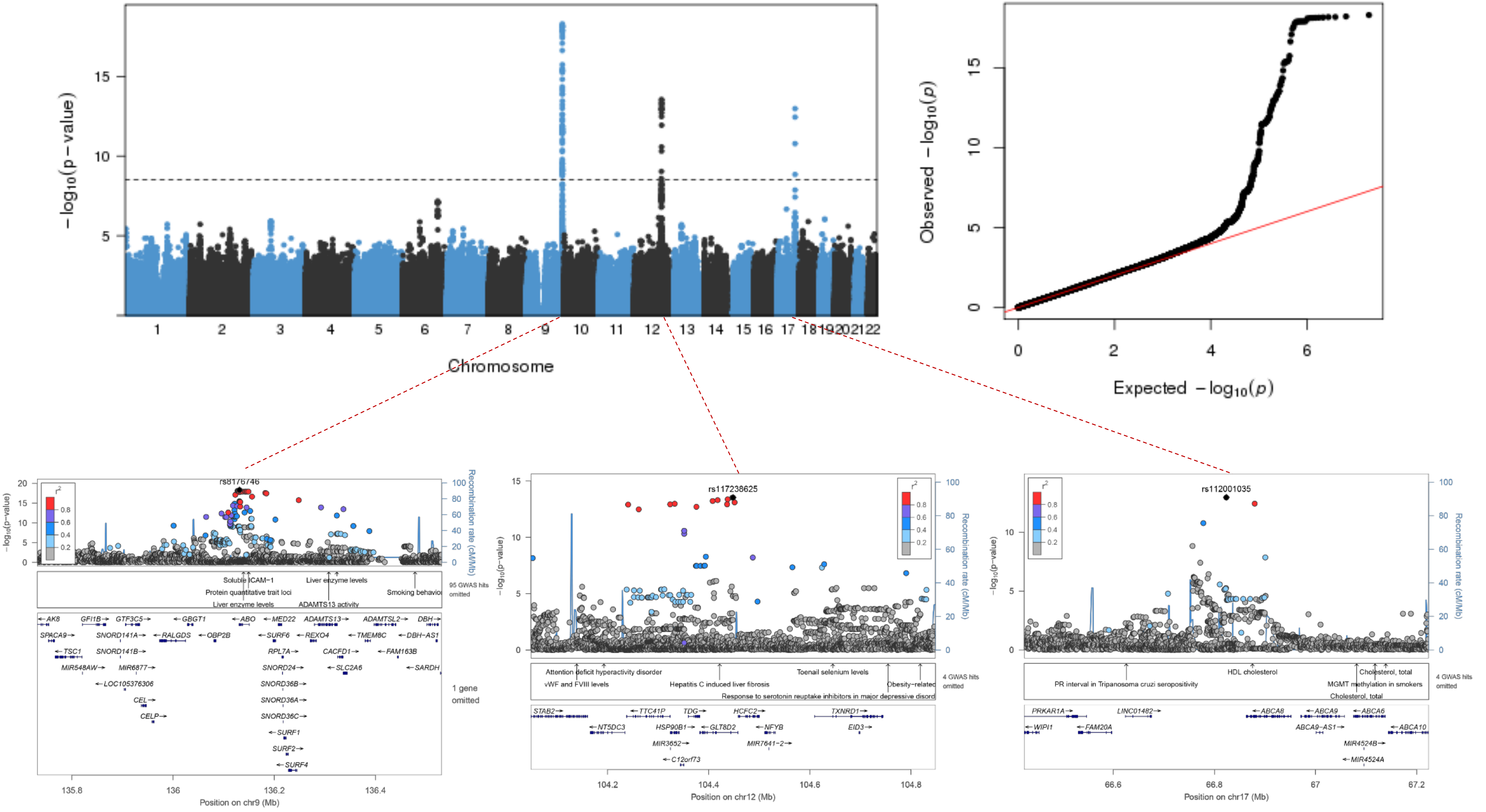

**Figure S2 D) Manhattan, Q-Q, and LocusZoom plots for VEGF results in NFBC1966.**

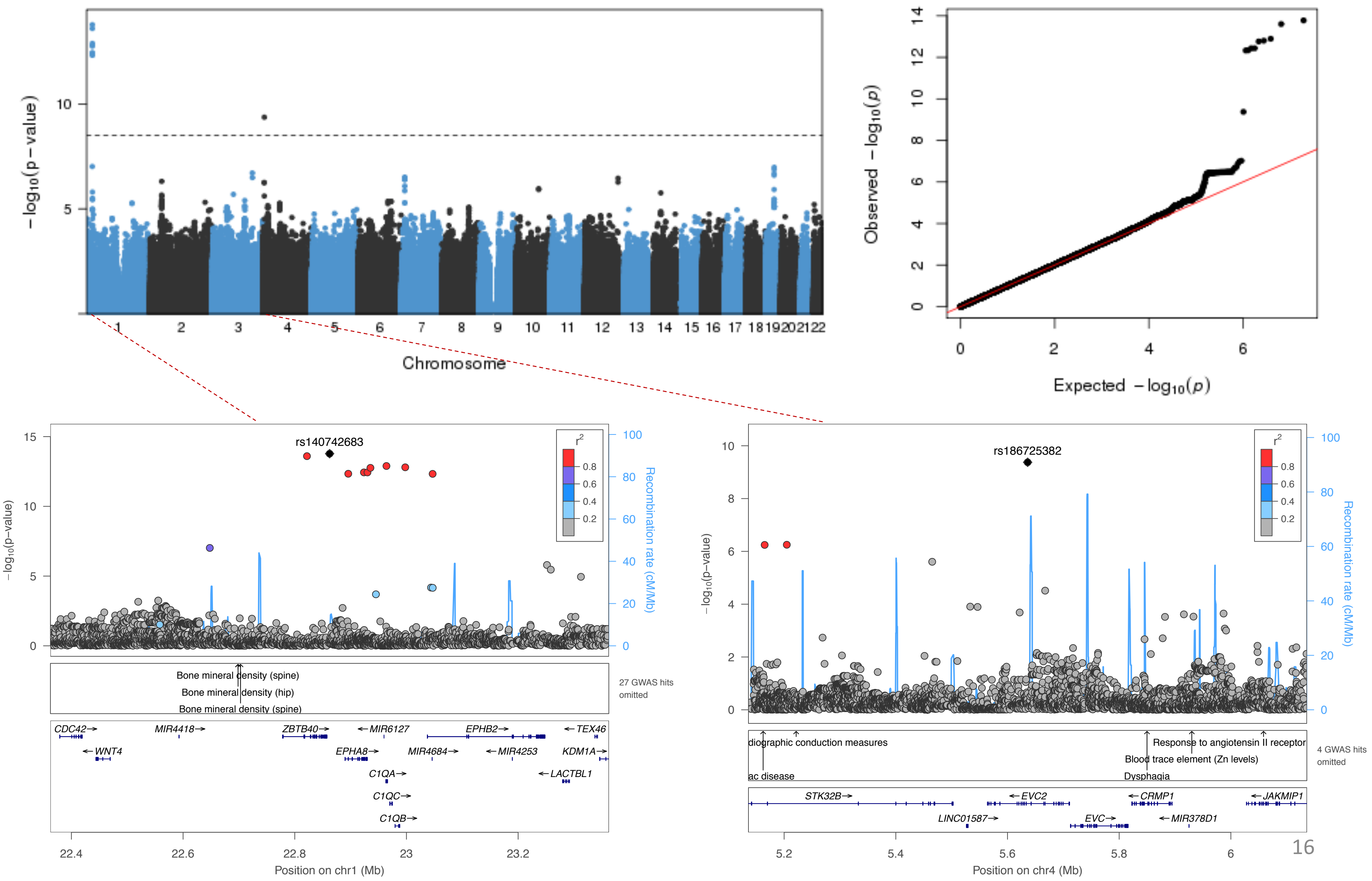

**Figure S2 E) Manhattan, Q-Q, and LocusZoom plots for VEGF results in meta-analyses.**

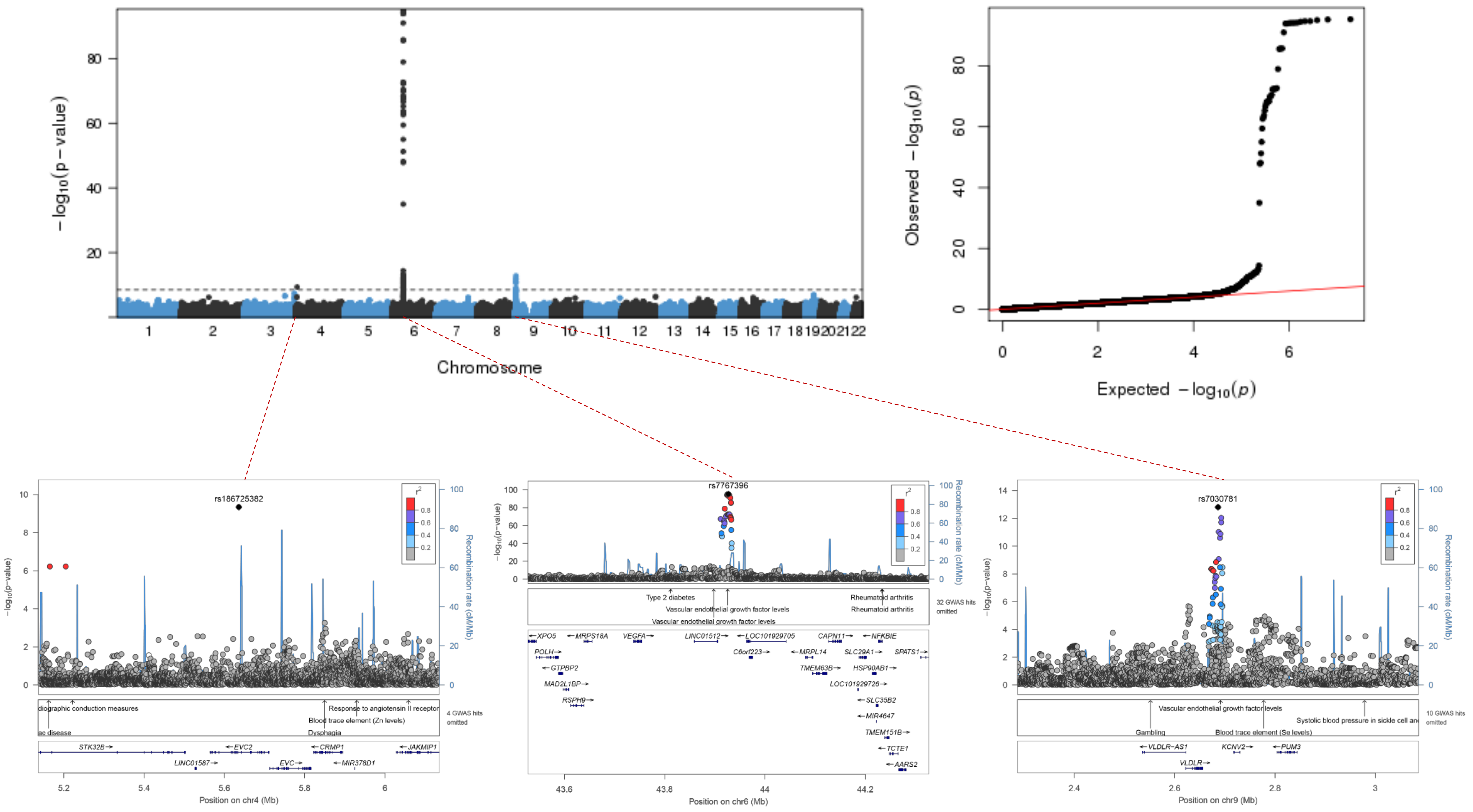

**Figure S2 F) Manhattan, Q-Q, and LocusZoom plots for TNFa results in NFBC1966.**

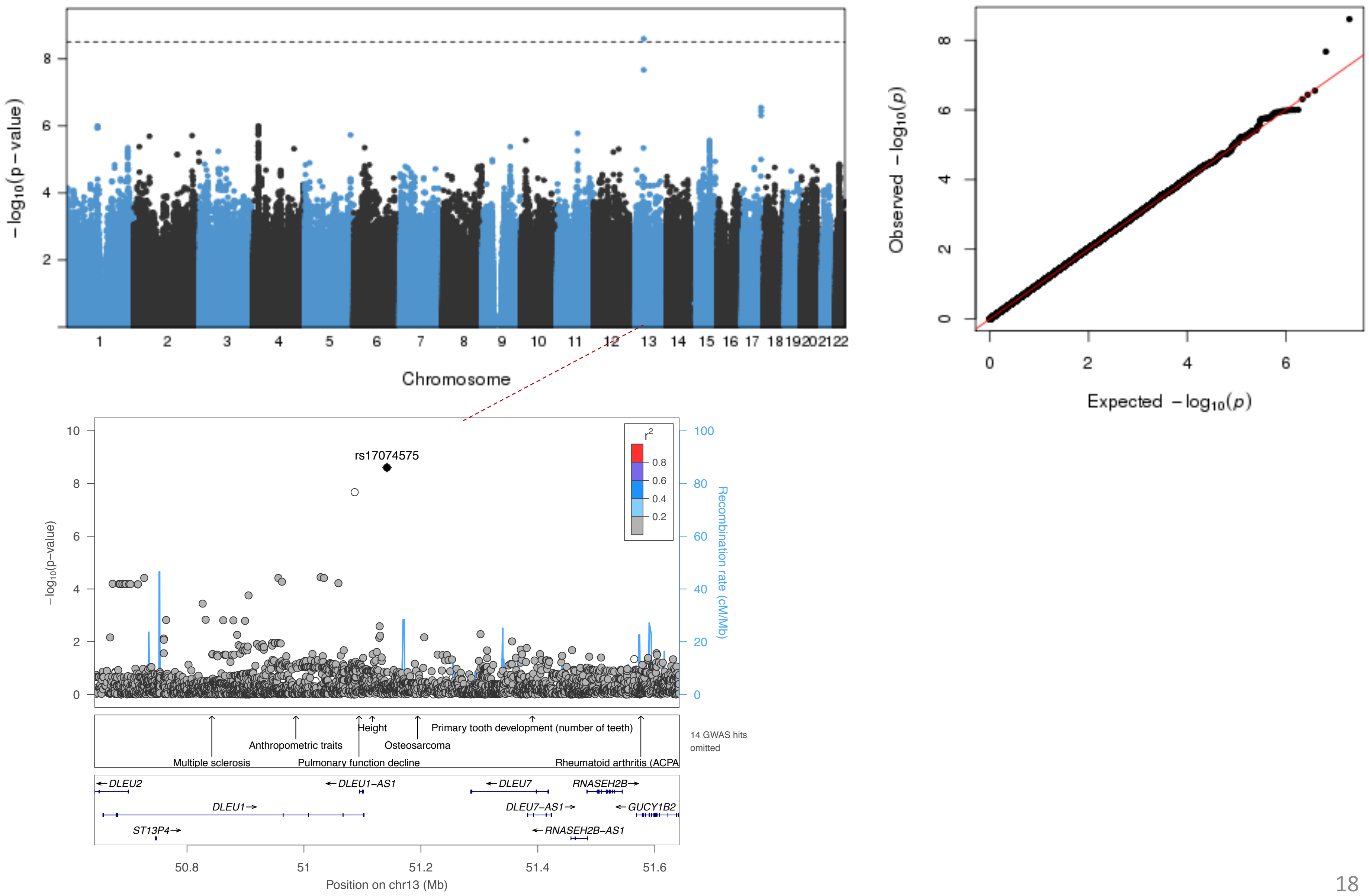

**Figure S2 G) Manhattan, Q-Q, and LocusZoom plots for TNFa results in meta-analyses.**

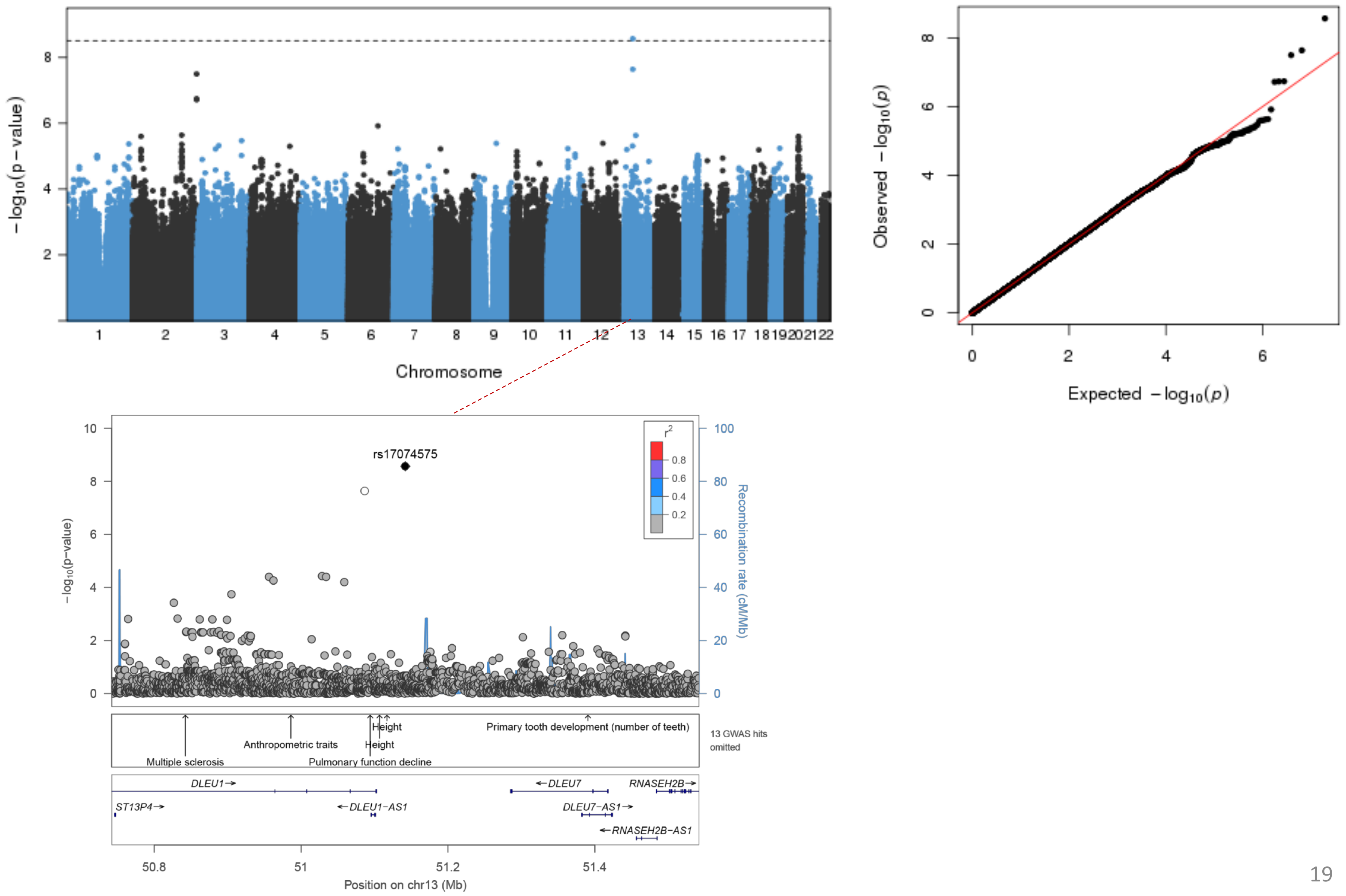

**Figure S2 H) Manhattan, Q-Q, and LocusZoom plots for IL-1a results in NFBC1966.**

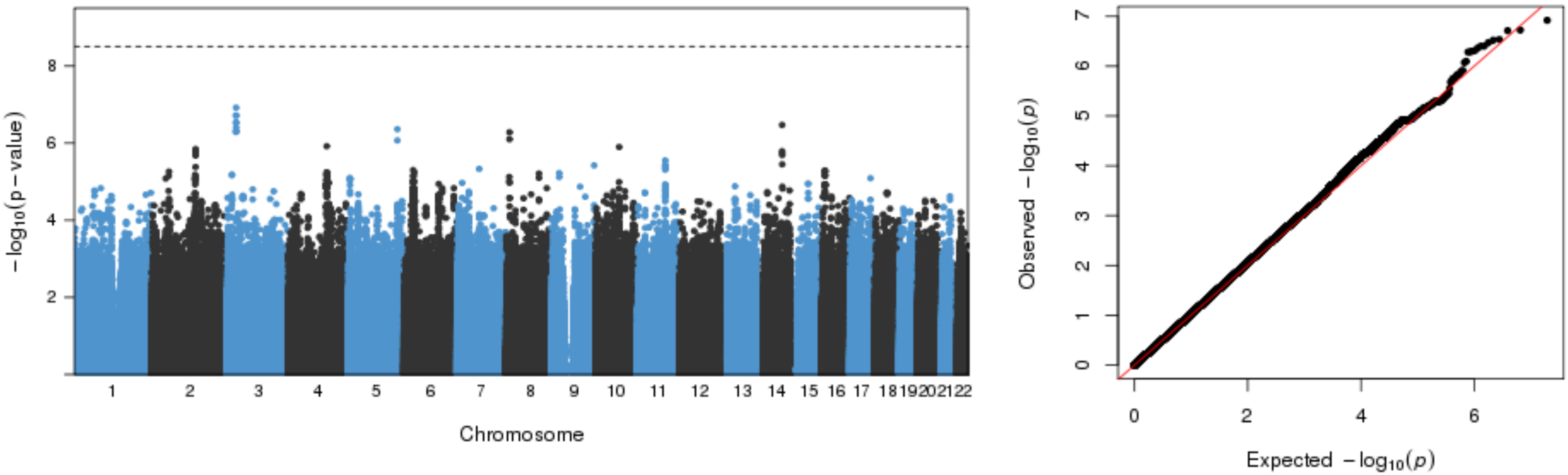


**No significant associtions (p-value limit 3.1x10$^{-9}$).**

**Figure S2 I) Manhattan, Q-Q, and LocusZoom plots for IL-1b results in NFBC1966.**

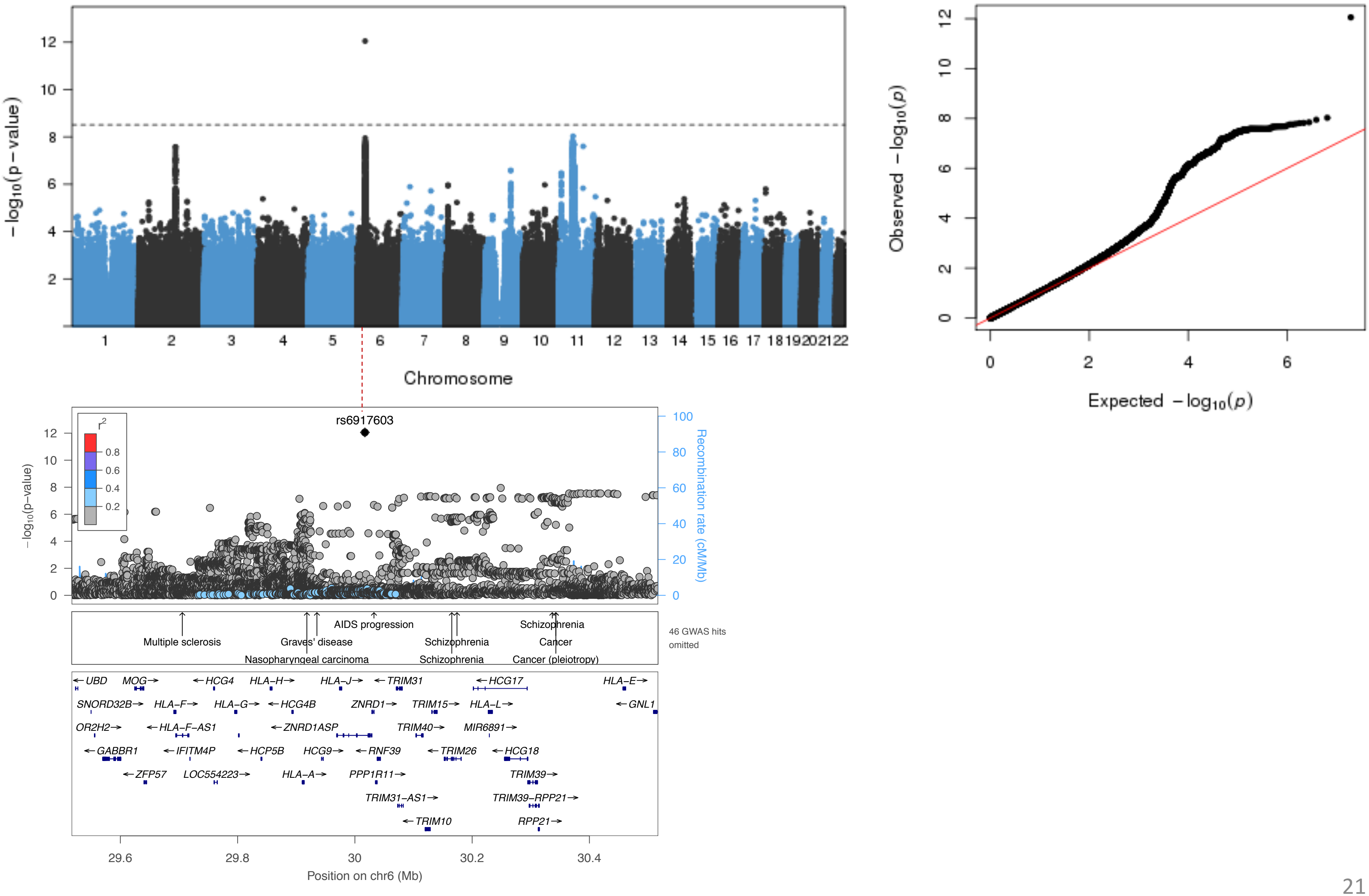

**Figure S2 J) Manhattan, Q-Q, and LocusZoom plots for IL-1b results in meta-analyses.**

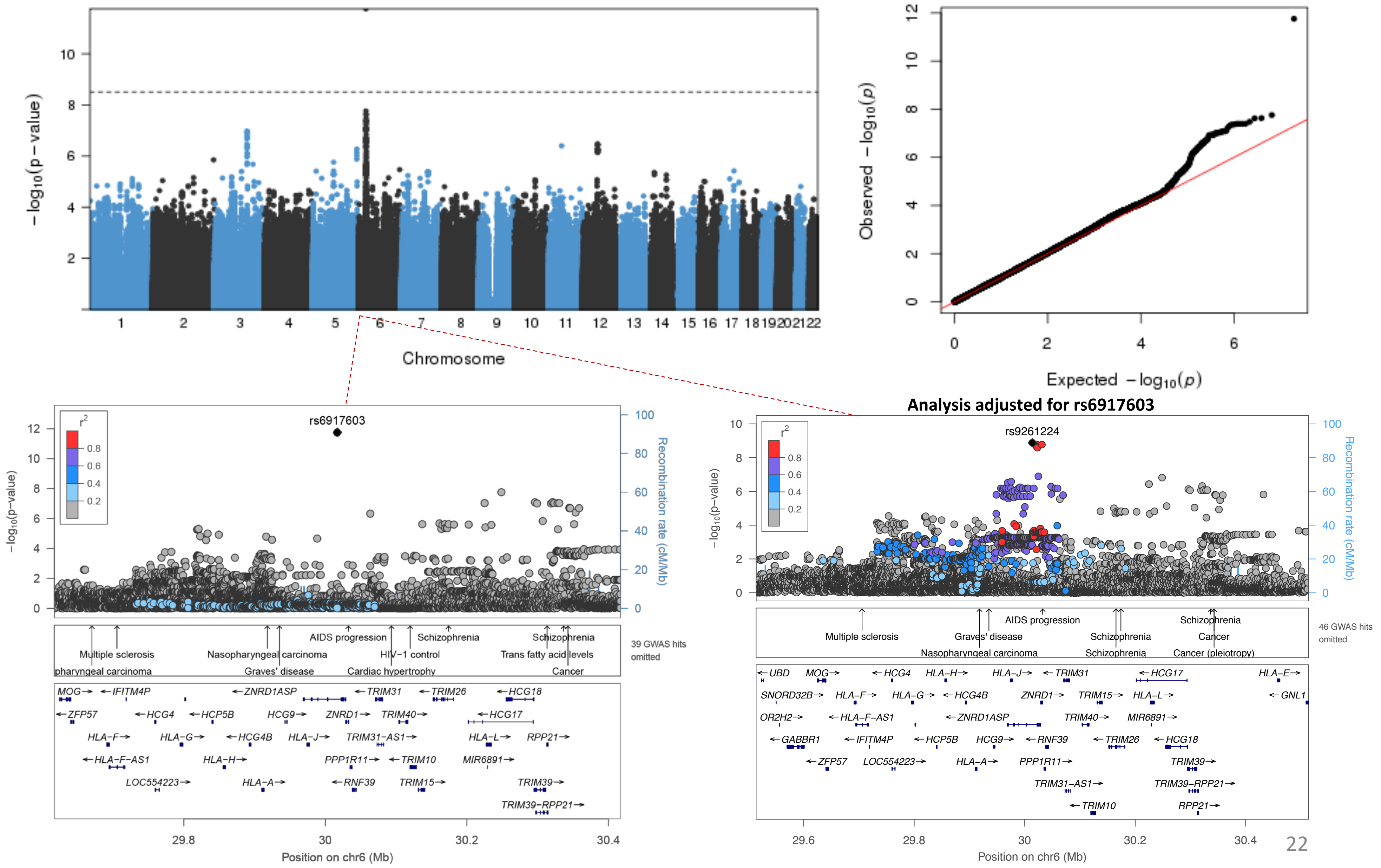

**Figure S2 K) Manhattan, Q-Q, and LocusZoom plots for IL4 results in NFBC1966.**

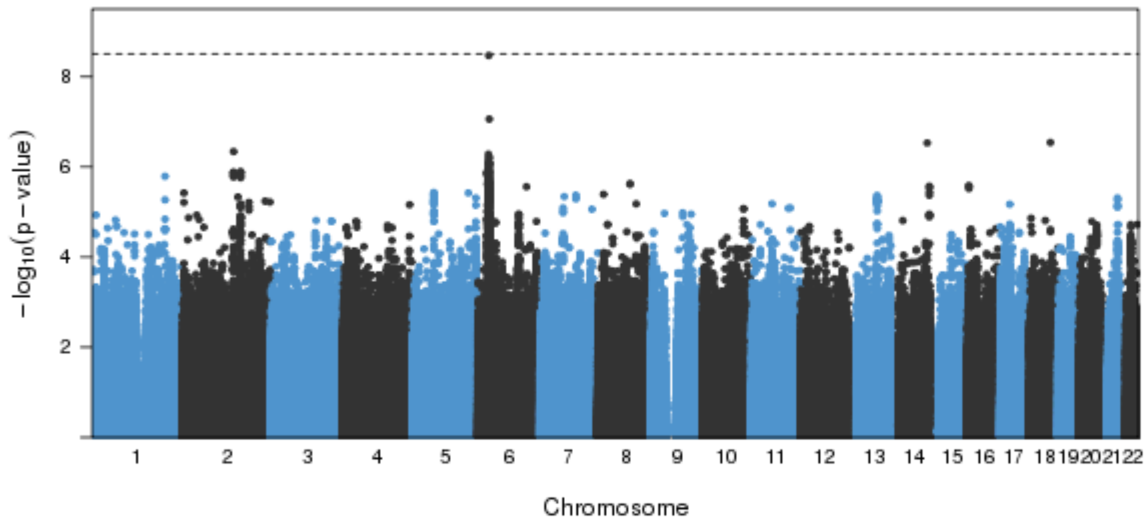


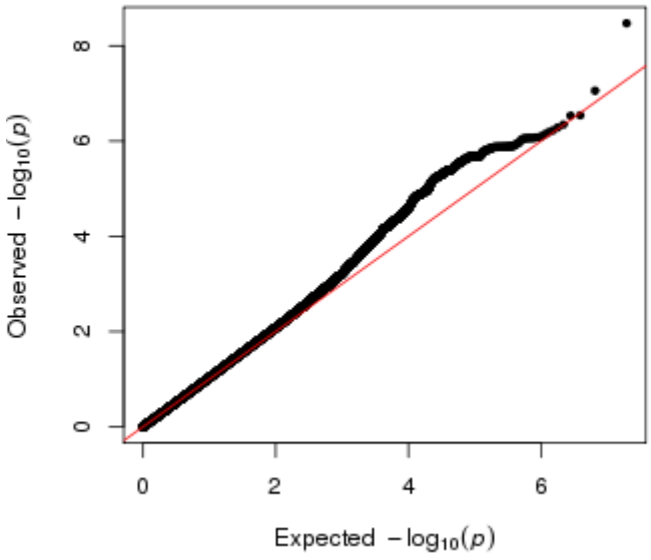


**No significant associtions (p-value limit 3.1x10$^{-9}$).**

**Figure S2 L) Manhattan, Q-Q, and LocusZoom plots for IL4 results in meta-analyses.**

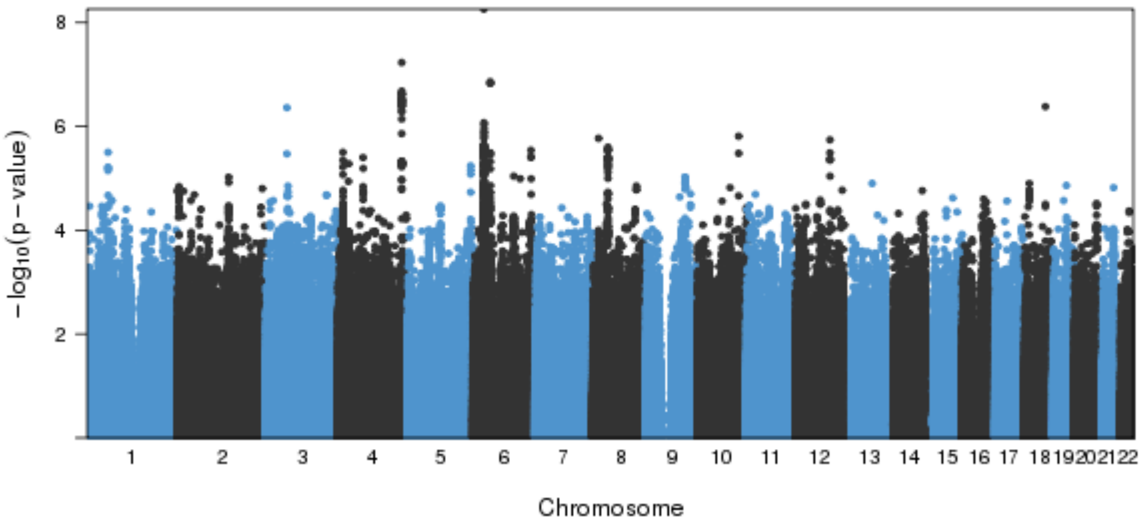


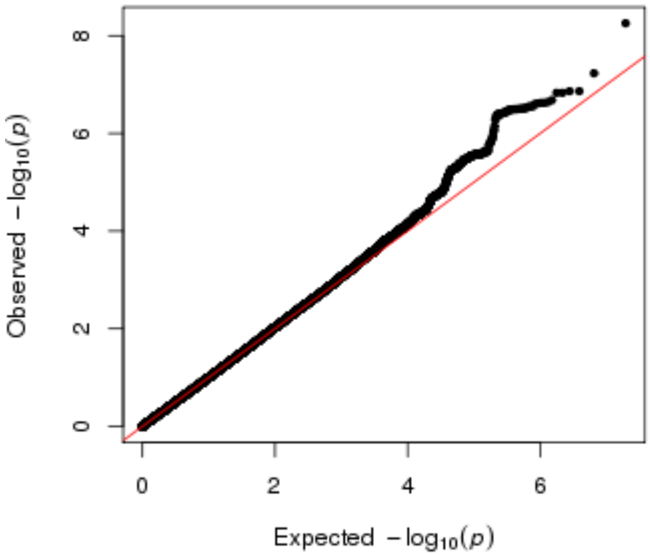


**No significant associtions (p-value limit $3.1x10^{-9}$).**

**Figure S2 M) Manhattan, Q-Q, and LocusZoom plots for IL6 results in NFBC1966.**

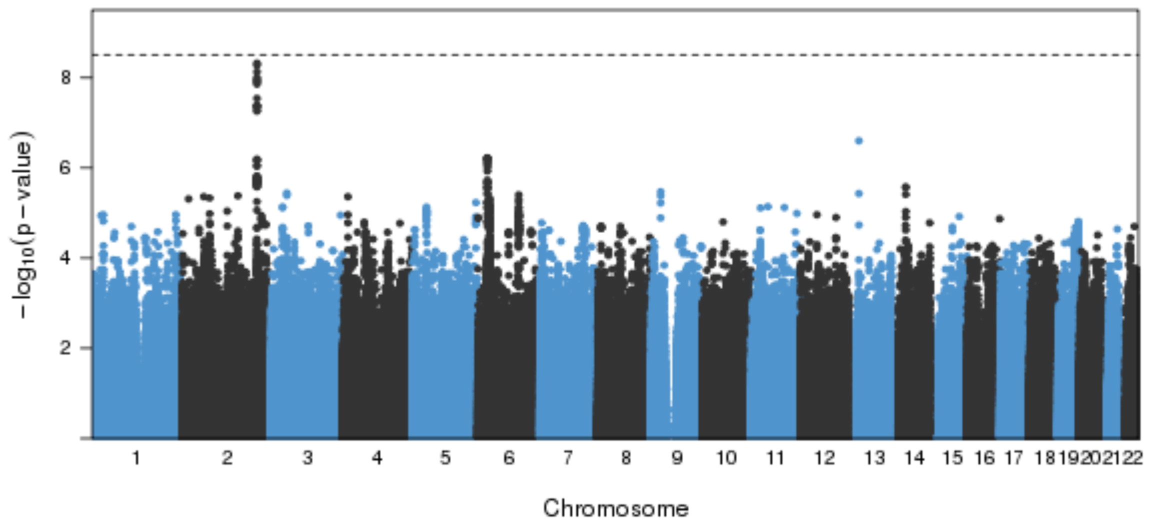


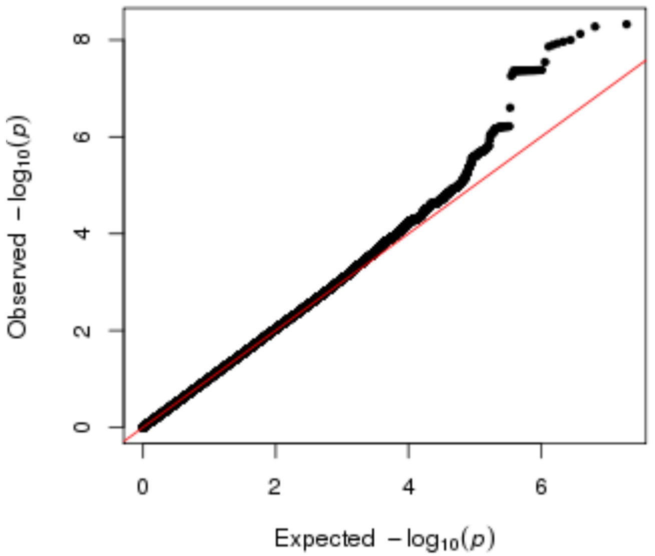


**No significant assocItions (p-value limit $3.1x10^{-9}$).**

**Figure S2 N) Manhattan, Q-Q, and LocusZoom plots for IL6 in meta-analyses.**

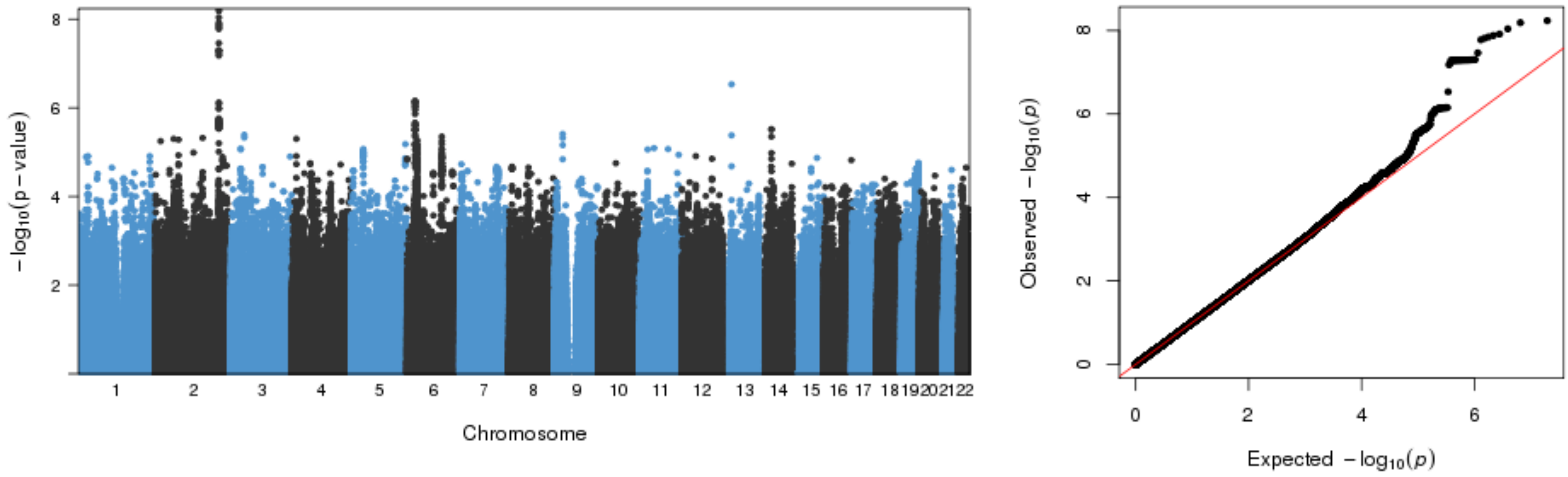


**No significant associtions (p-value limit $3.1x10^{-9}$).**

**Figure S2 O) Manhattan, Q-Q, and LocusZoom plots for IL8 results in NFBC1966.**

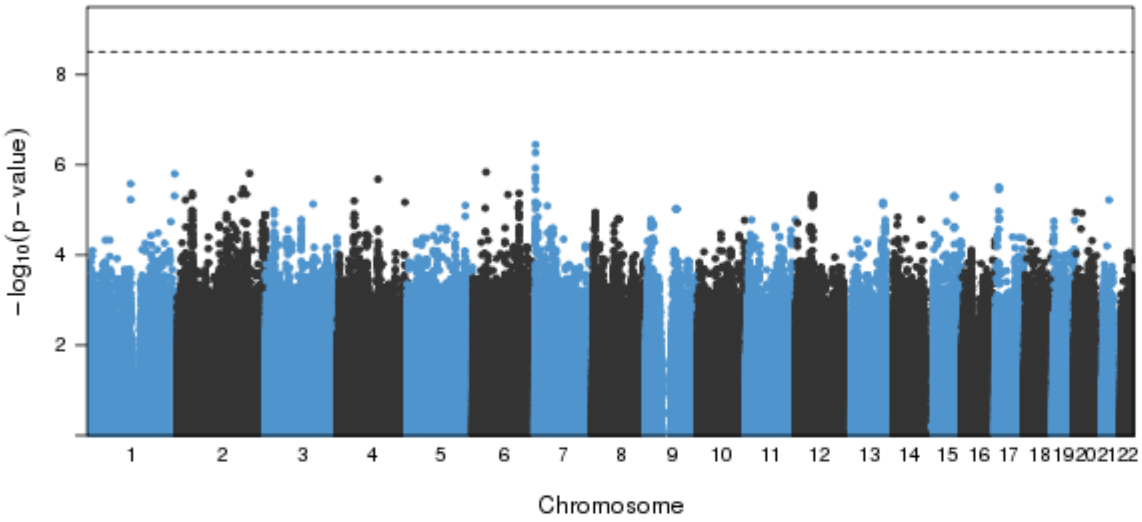


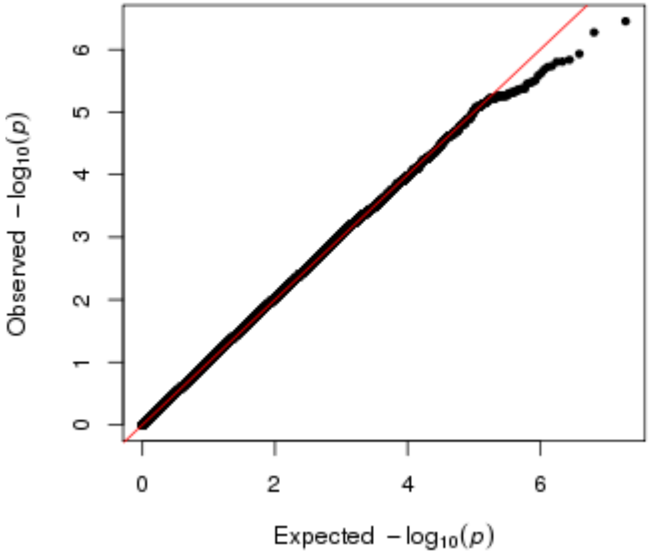


**No significant associtions (p-value limit 3.1x10$^{-9}$).**

**Figure S2 P) Manhattan, Q-Q, and LocusZoom plots for IL8 results in meta-analyses.**

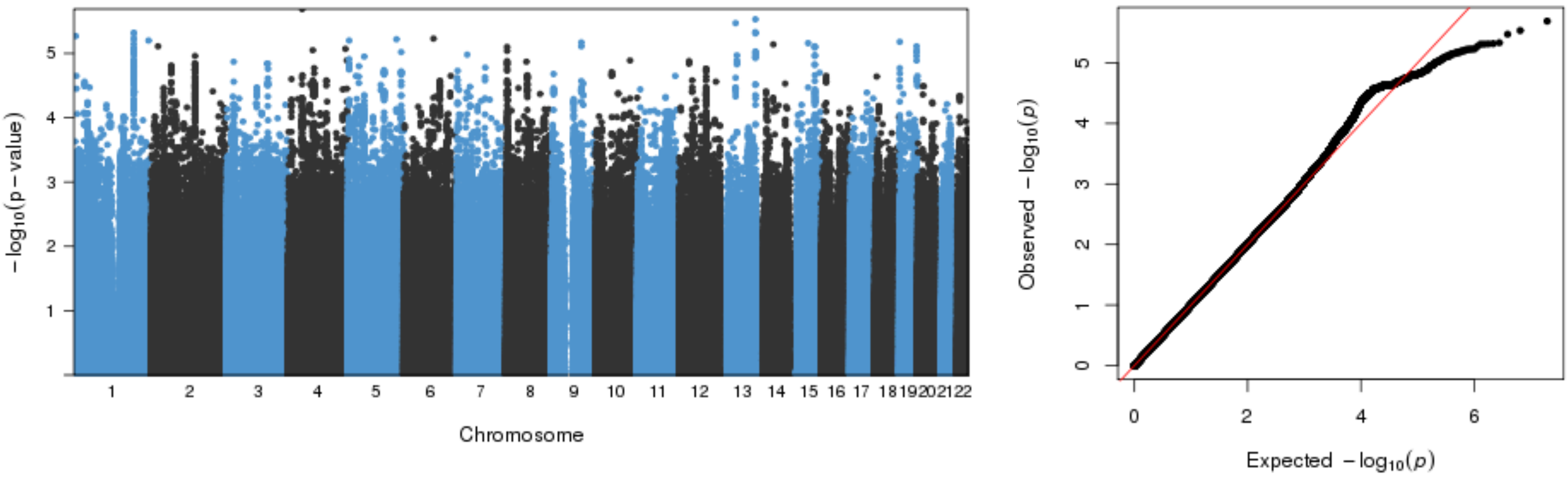


**No significant associtions (p-value limit 3.1x10$^{-9}$).**

**Figure S2 Q) Manhattan, Q-Q, and LocusZoom plots for IL17 results in NFBC1966.**

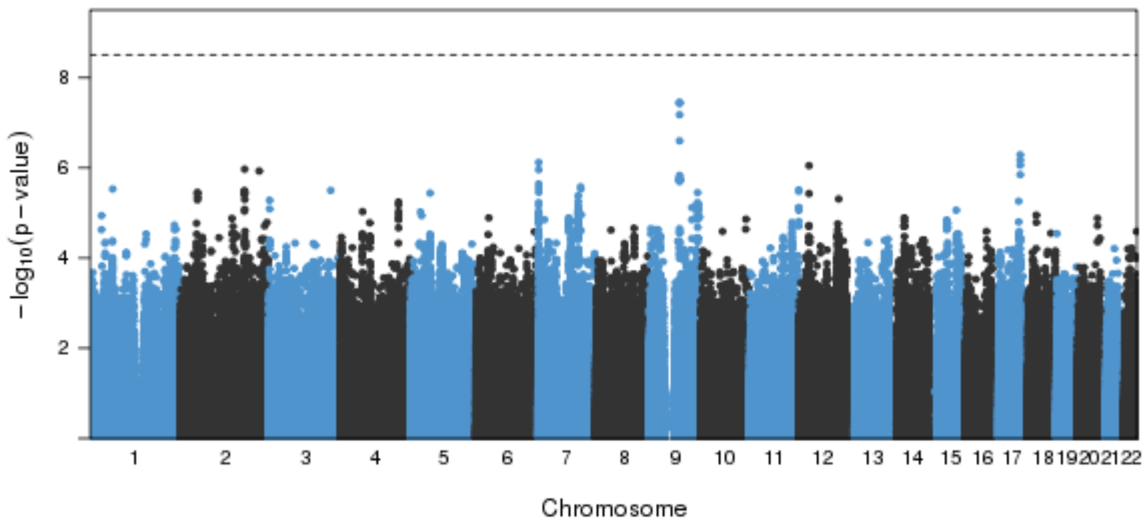


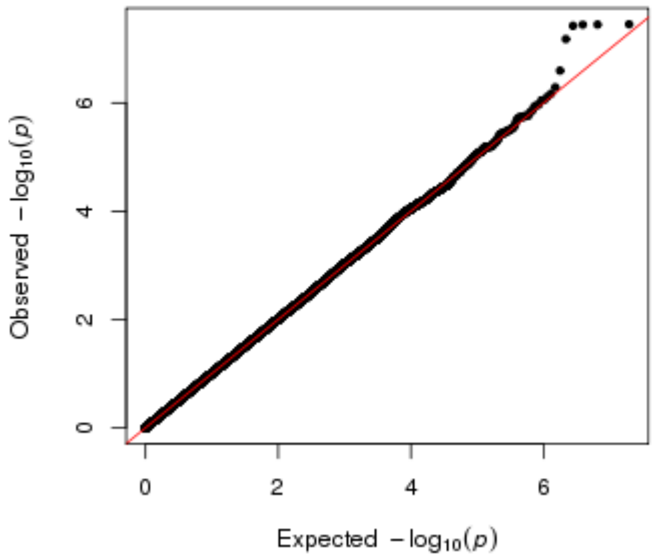


**No significant associtions (p-value limit $3.1x10^{-9}$).**

**Figure S2 R) Manhattan, Q-Q, and LocusZoom plots for IL17 results in meta-analyses.**

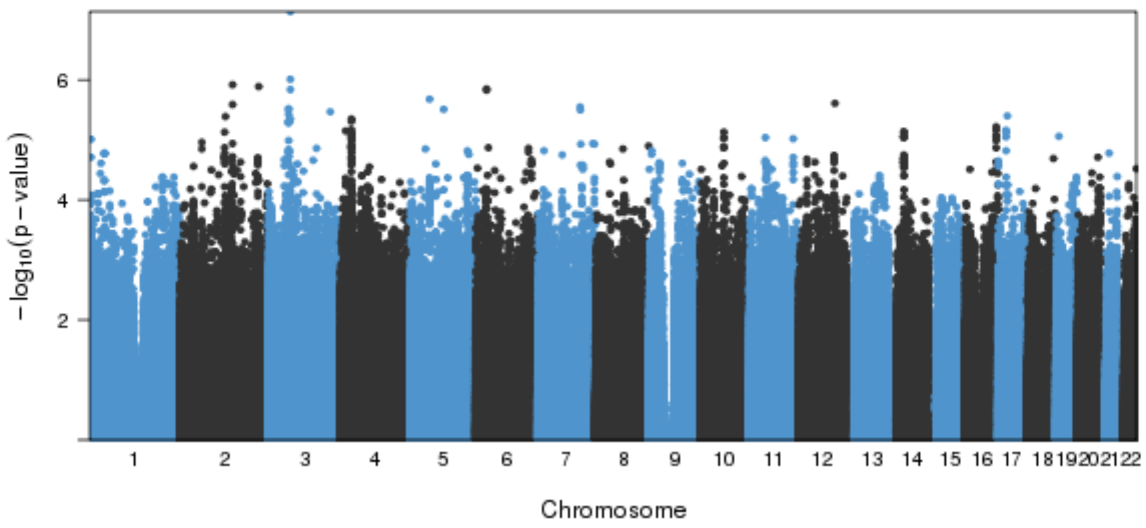


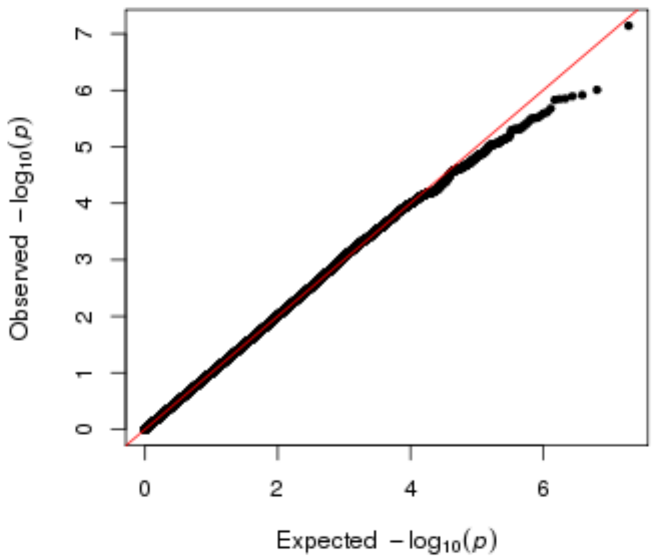


**No significant assocItions (p-value limit $3.1x10^{-9}$).**

**Figure S2 S) Manhattan, Q-Q, and LocusZoom plots for IL1ra results in NFBC1966.**

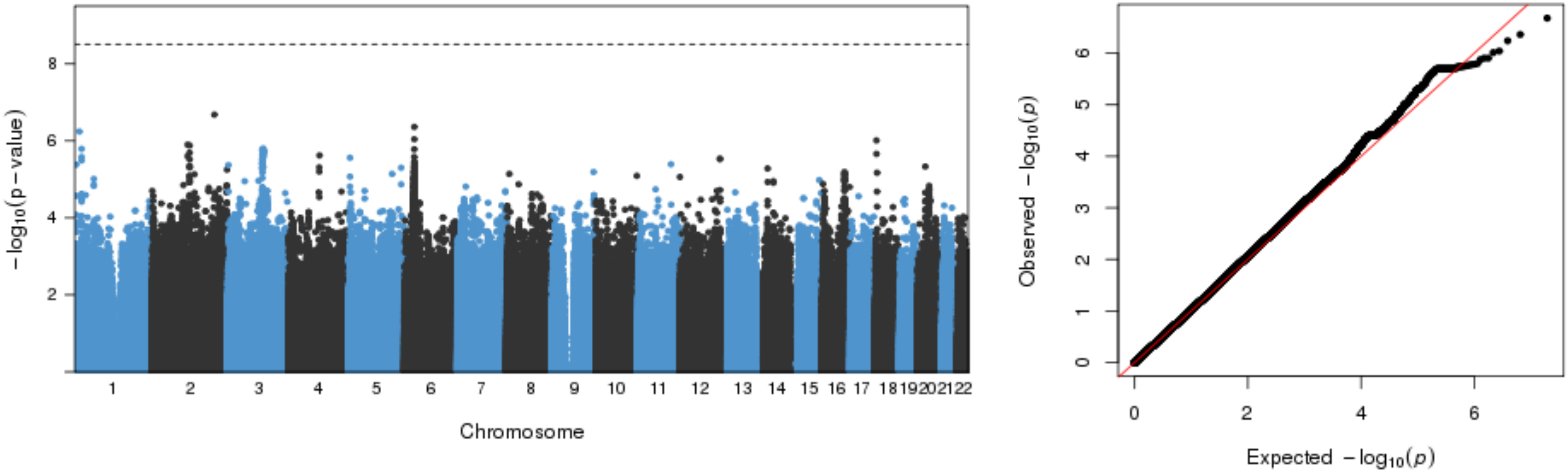


**No significant assocition s (p-value limit 3.1x10$^{-9}$).**

**Figure S2 T) Manhattan, Q-Q, and LocusZoom plots for IL1ra results in meta-analyses.**

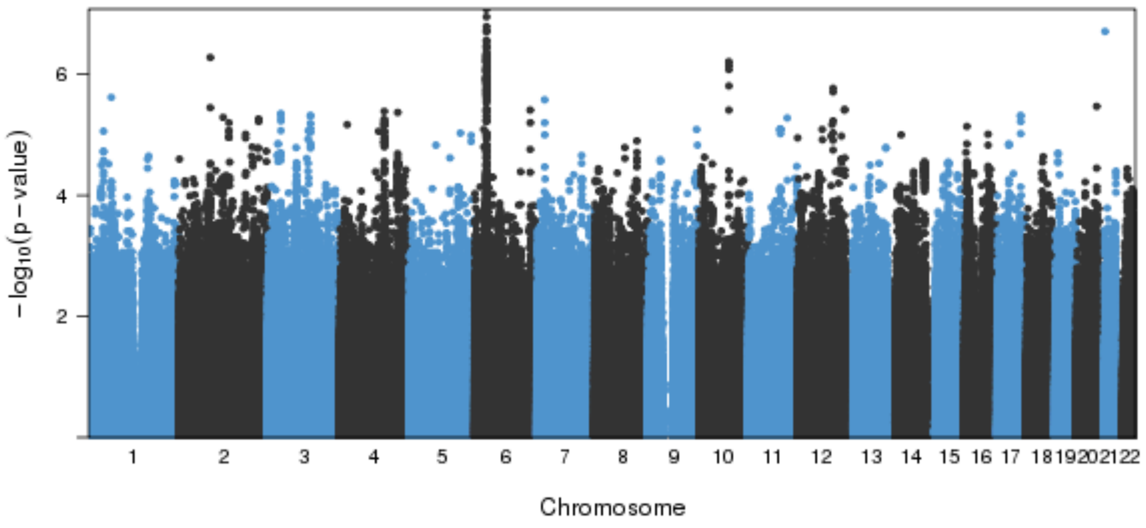


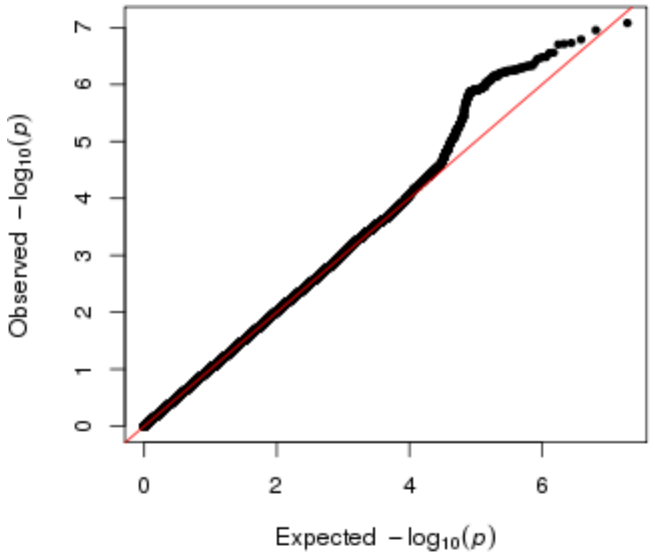


**No significant associtions (p-value limit 3.1x10$^{-9}$).**

**Figure S2 U) Manhattan, Q-Q, and LocusZoom plots for IP10 results in NFBC1966.**

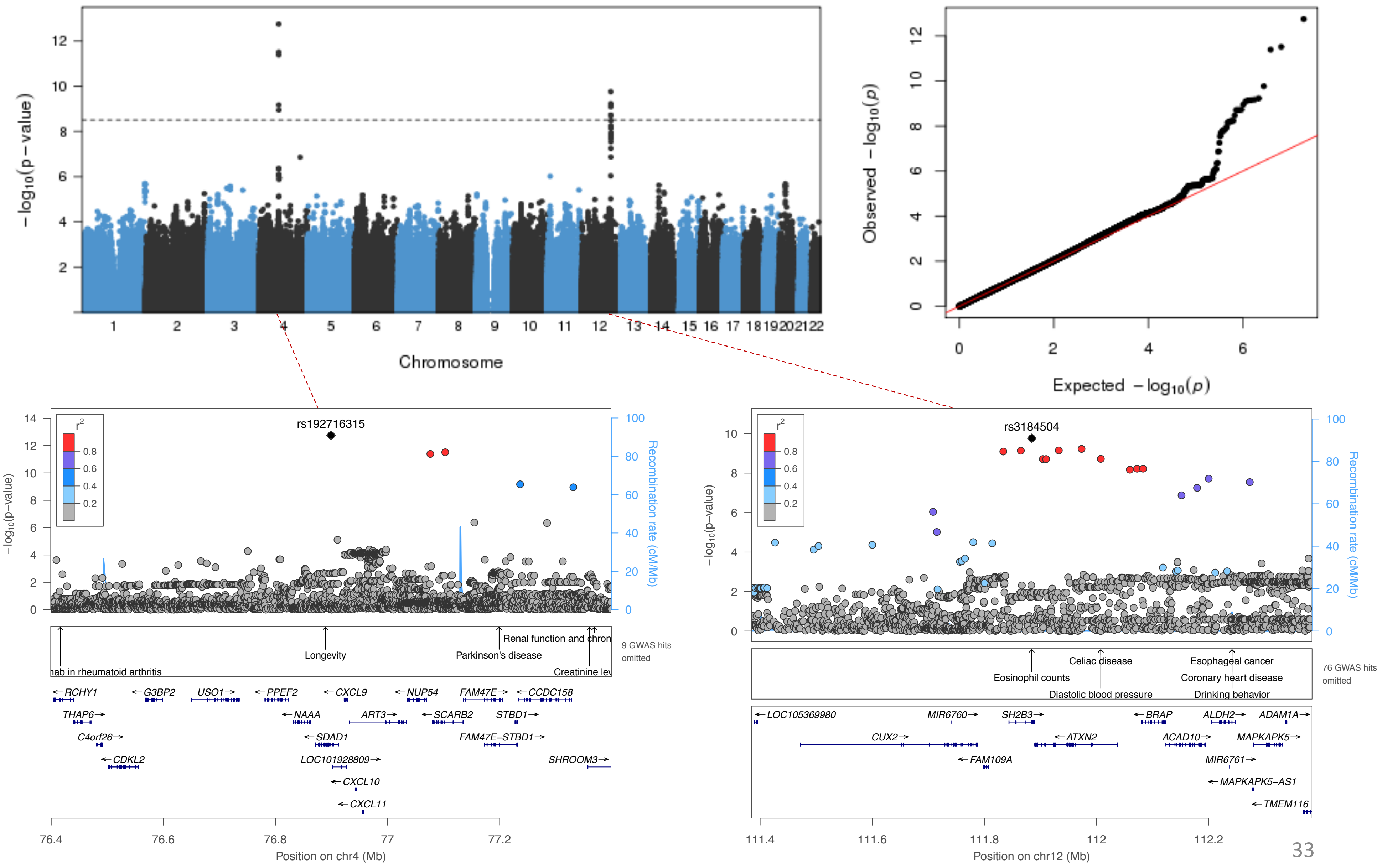

**Figure S2 V) Manhattan, Q-Q, and LocusZoom plots for IP10 results in meta-analyses.**

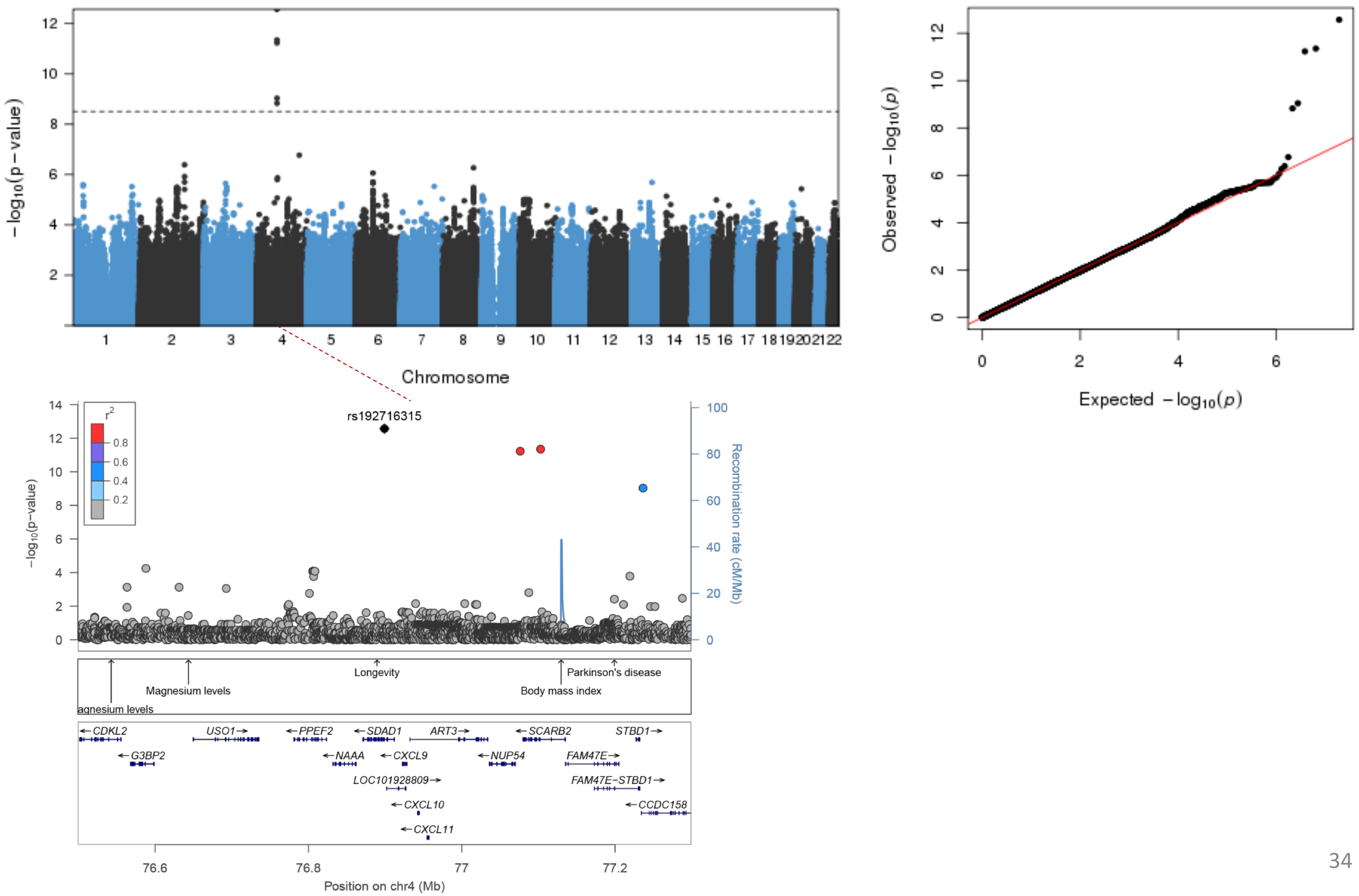

**Figure S2 W) Manhattan, Q-Q, and LocusZoom plots for MCP1 results in NFBC1966.**

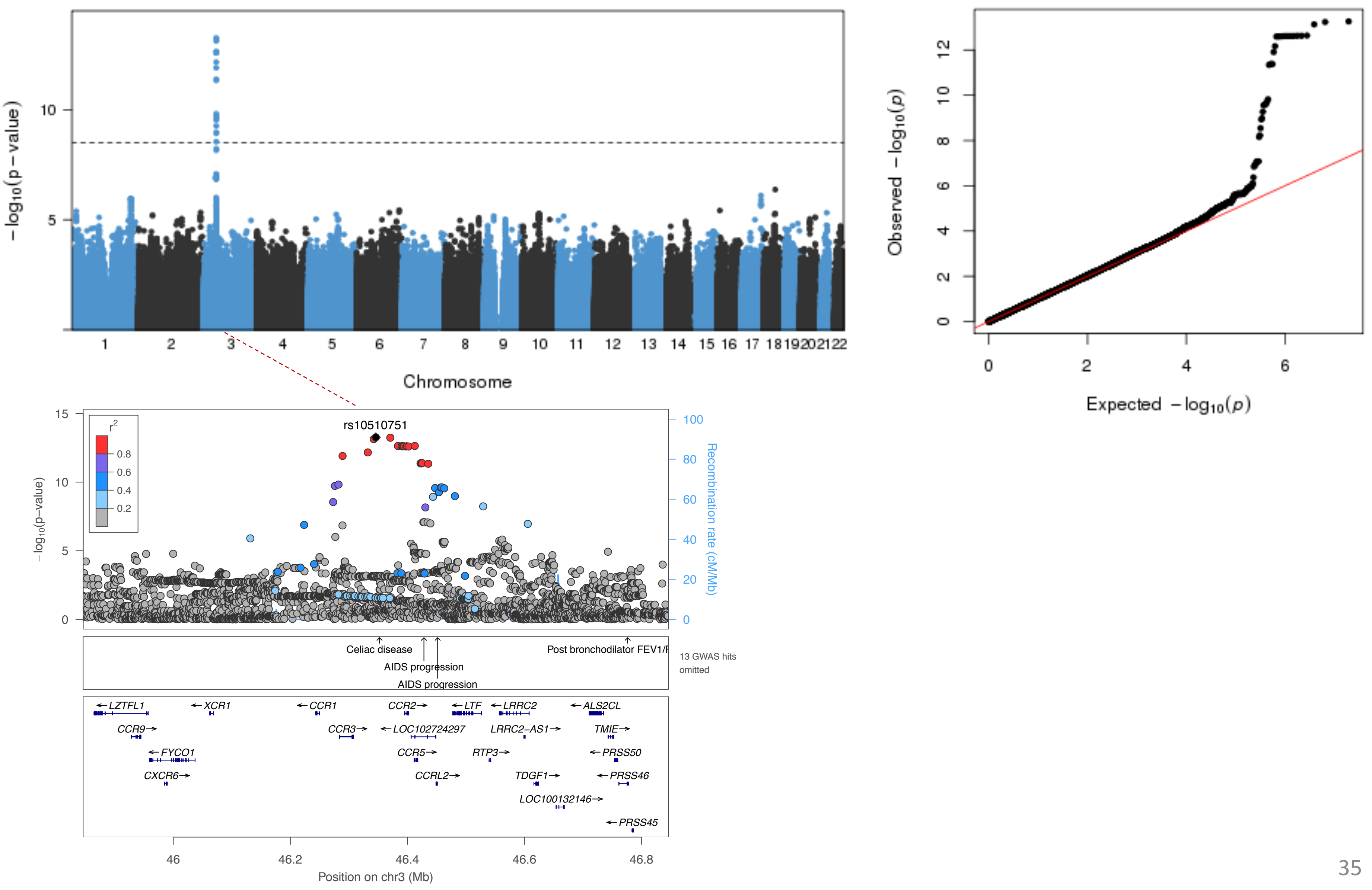

**Figure S2 X) Manhattan, Q-Q, and LocusZoom plots for MCP1 results in meta-analyses.**

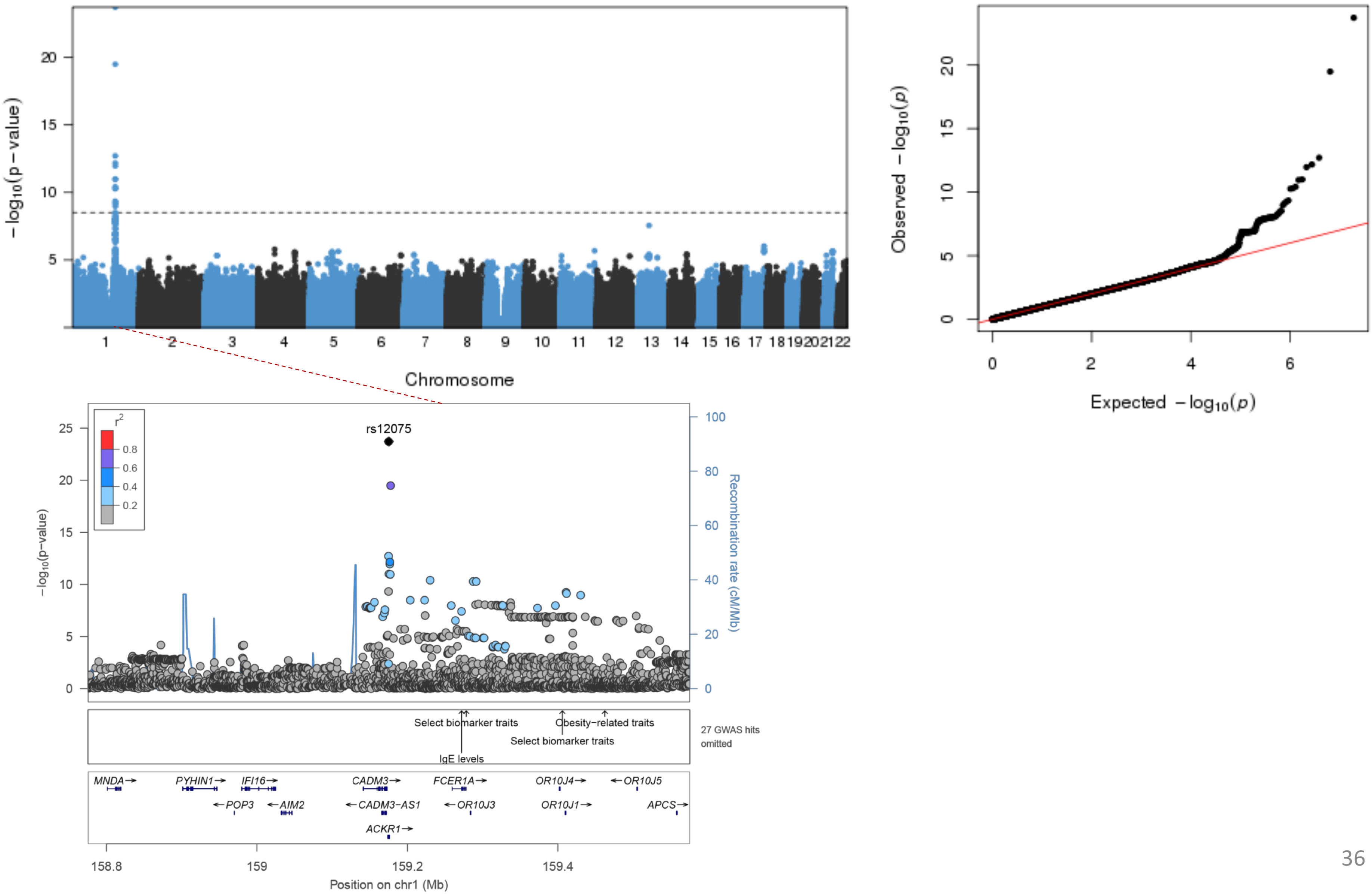

**Figure S2 Y) Manhattan, Q-Q, and LocusZoom plots for sCD40L results in NFBC1966.**

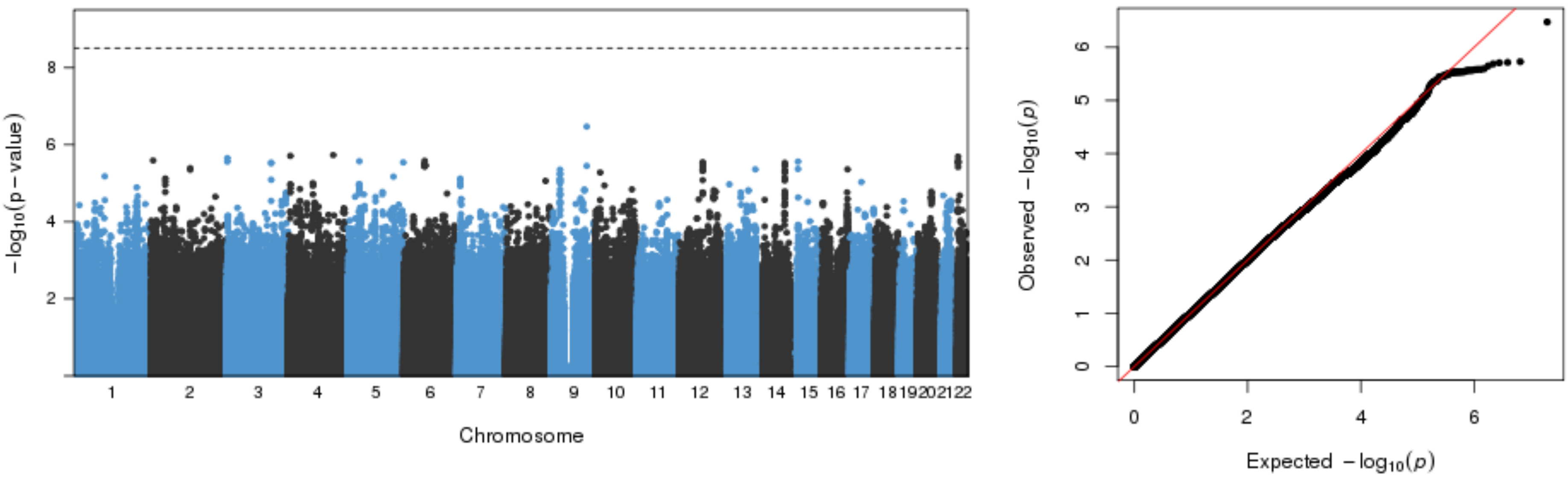


**No significant associtions (p-value limit $3.1x10^{-9}$).**

**Figure S2 Z) Manhattan, Q-Q, and LocusZoom plots for PAI-1 results in NFBC1966.**

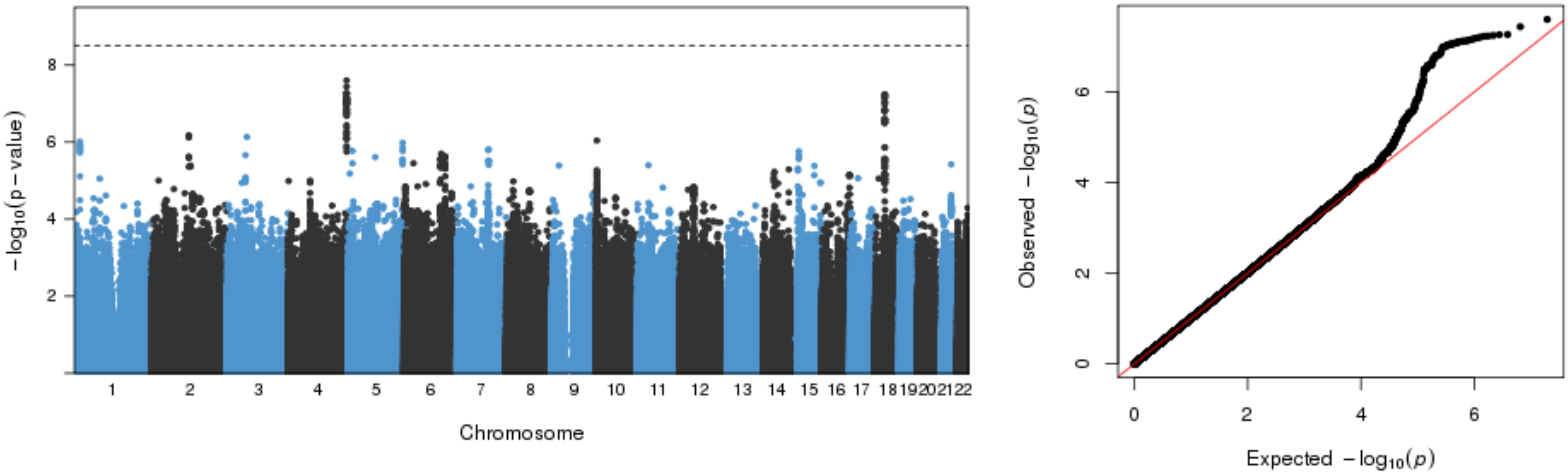


**No significant associtions (p-value limit 3.1x10$^{-9}$).**

**Figure S3 A). SNP effect correlations as scatter plots; sE-selectin at *ABO.***

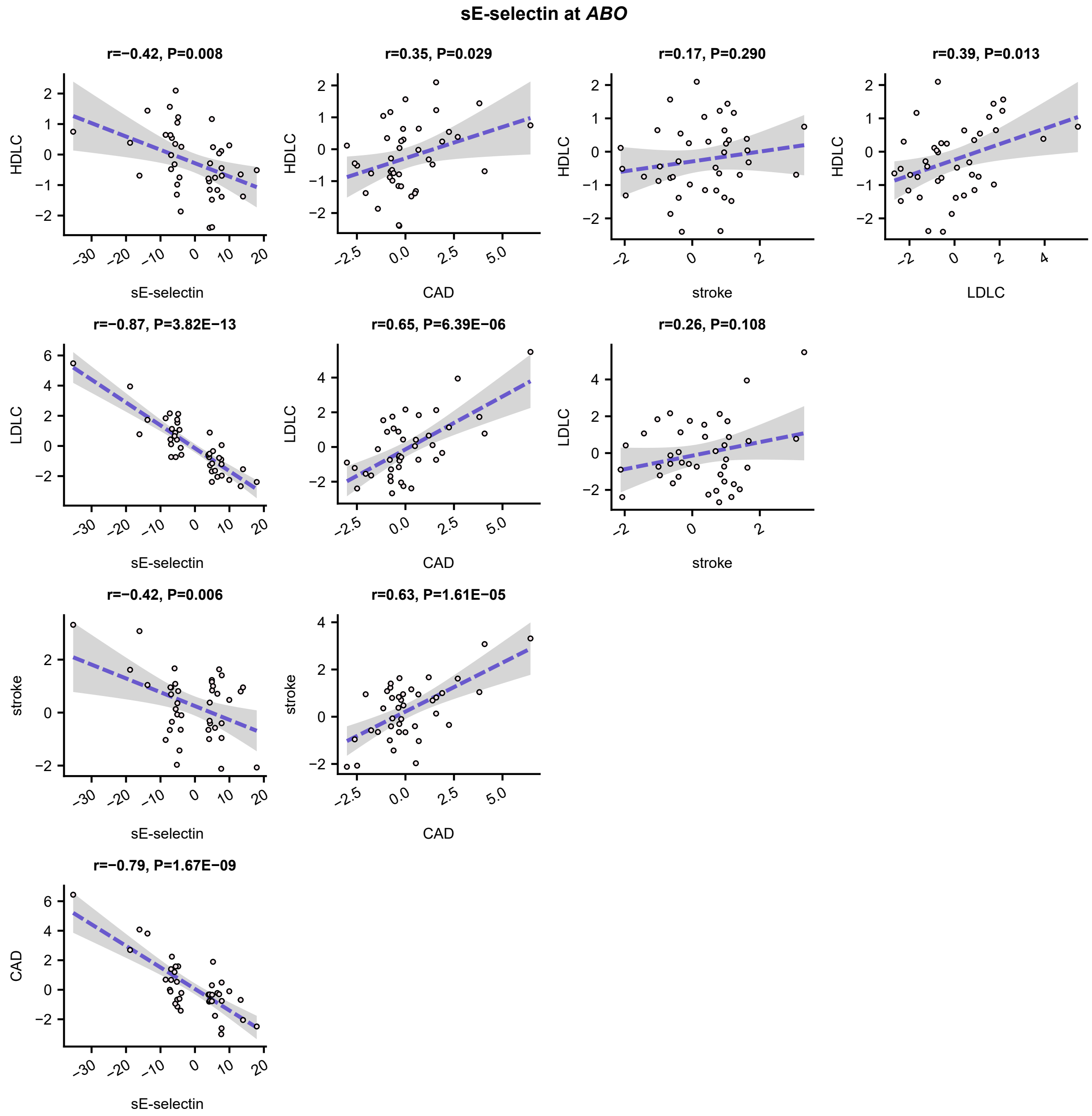

**Figure S3 B). SNP effect correlations as scatter plots; sICAM-1 at *ABO.***

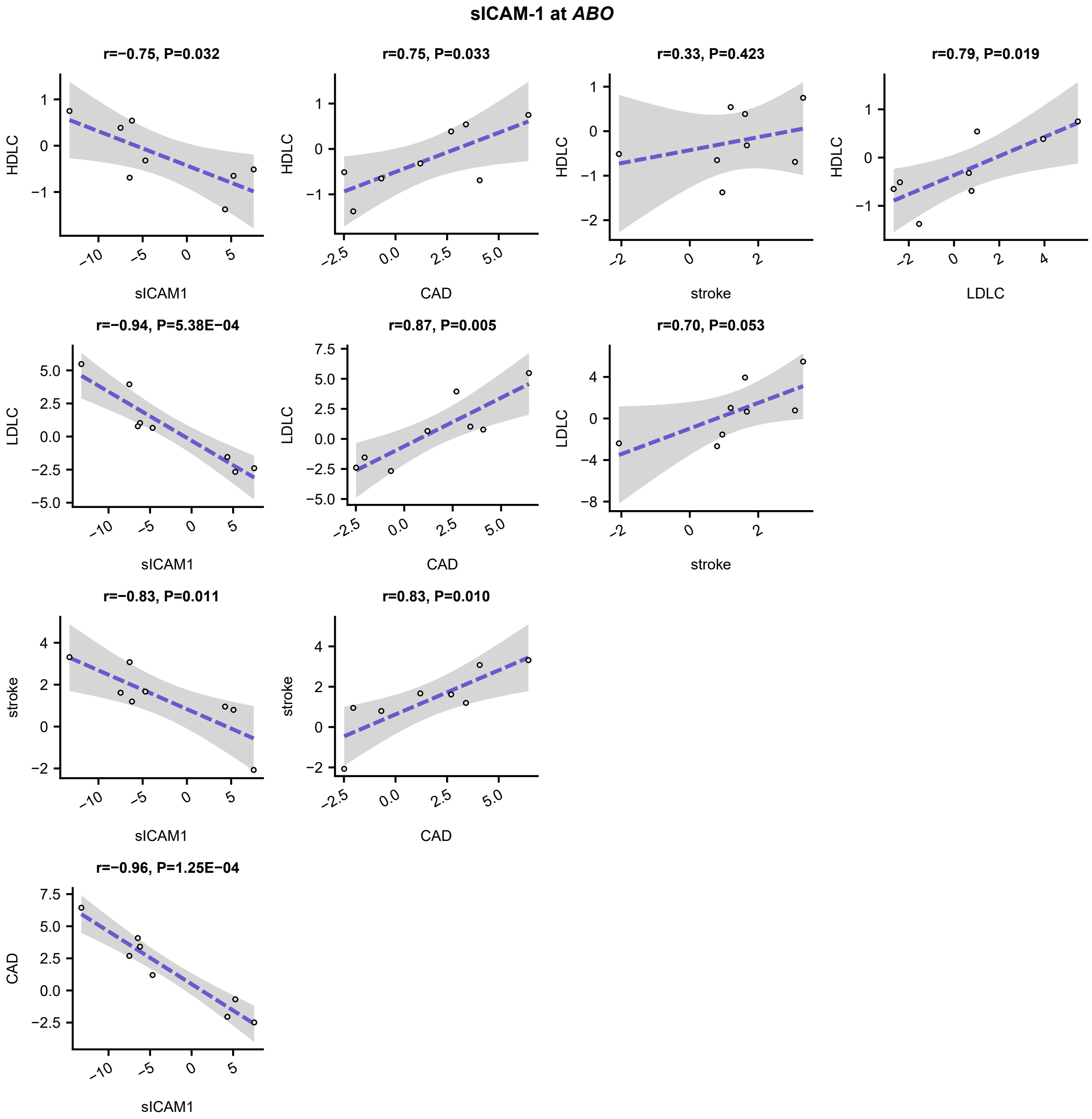

**Figure S3 C). SNP effect correlations as scatter plots; sVCAM-1 at *ABO.***

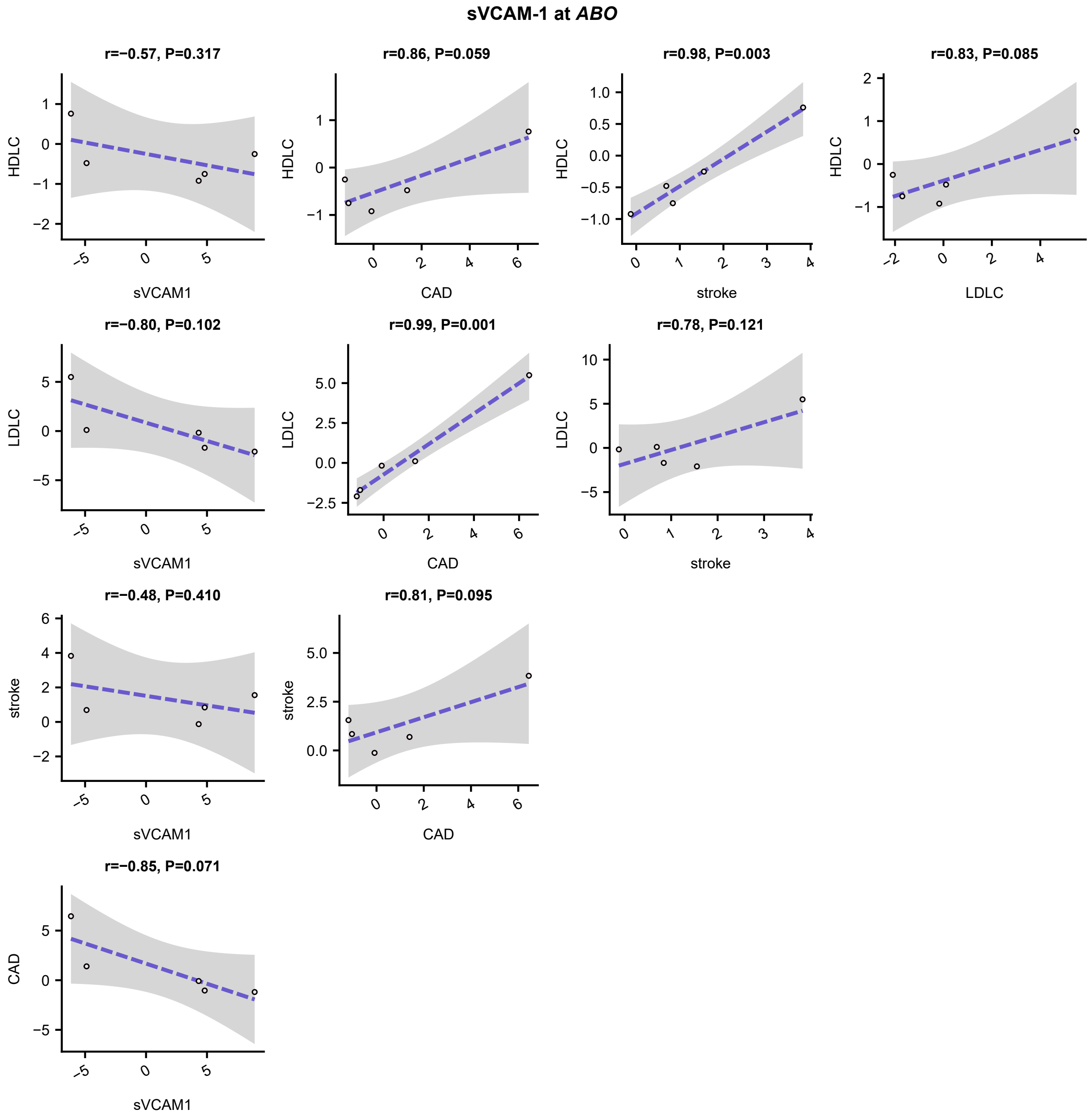

**Figure S3 D). SNP effect correlations as scatter plots; sVCAM-1 at *HSP90B1.***

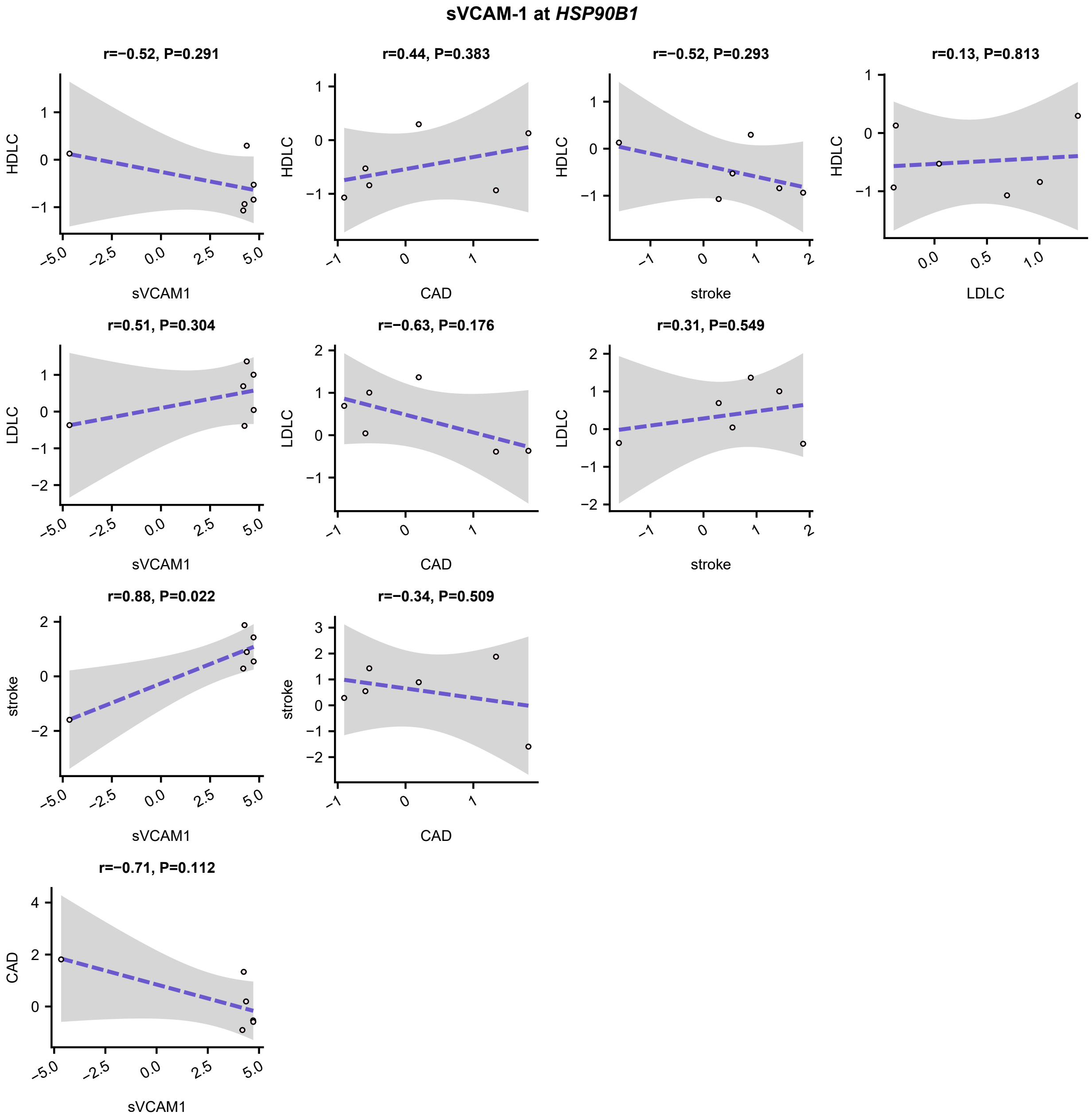

**Figure S3 E). SNP effect correlations as scatter plots; sE-selectin at *ST3GAL4.***

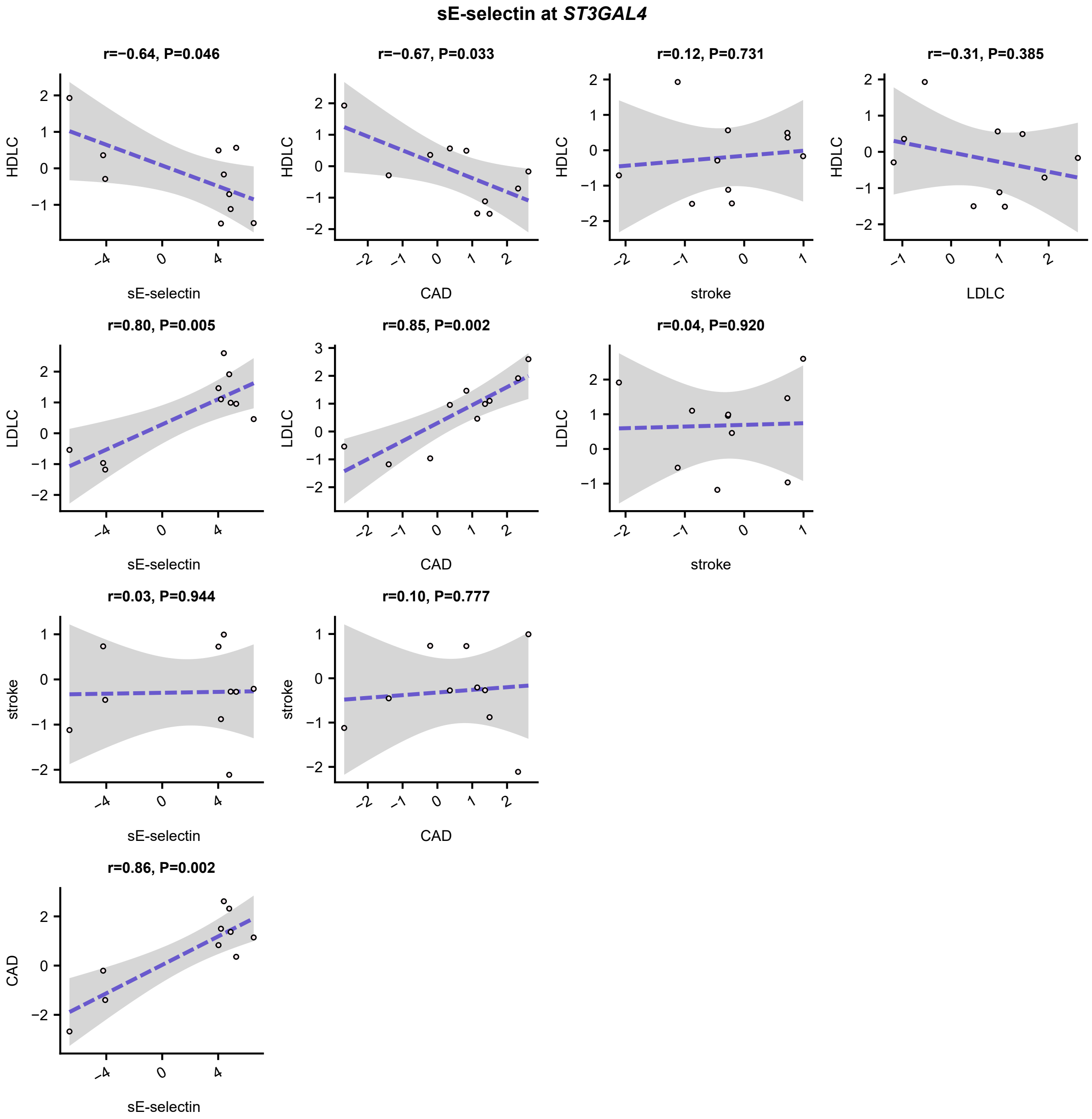

**Figure S3 F). SNP effect correlations as scatter plots; sICAM-1 at *ICAM1*.**

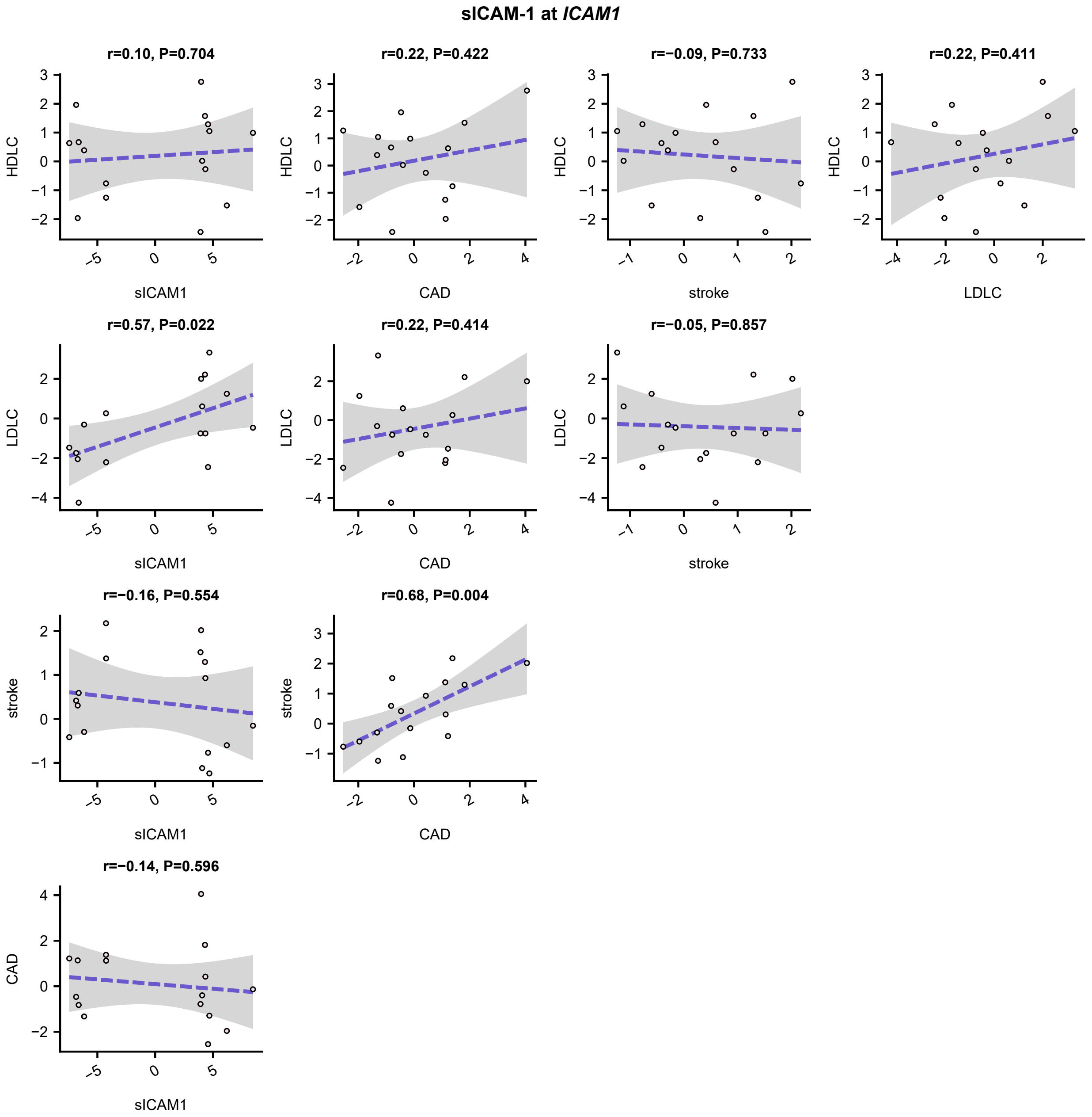

**Figure S3 G). SNP effect correlations as scatter plots; VEGF at *VLDLR*.**

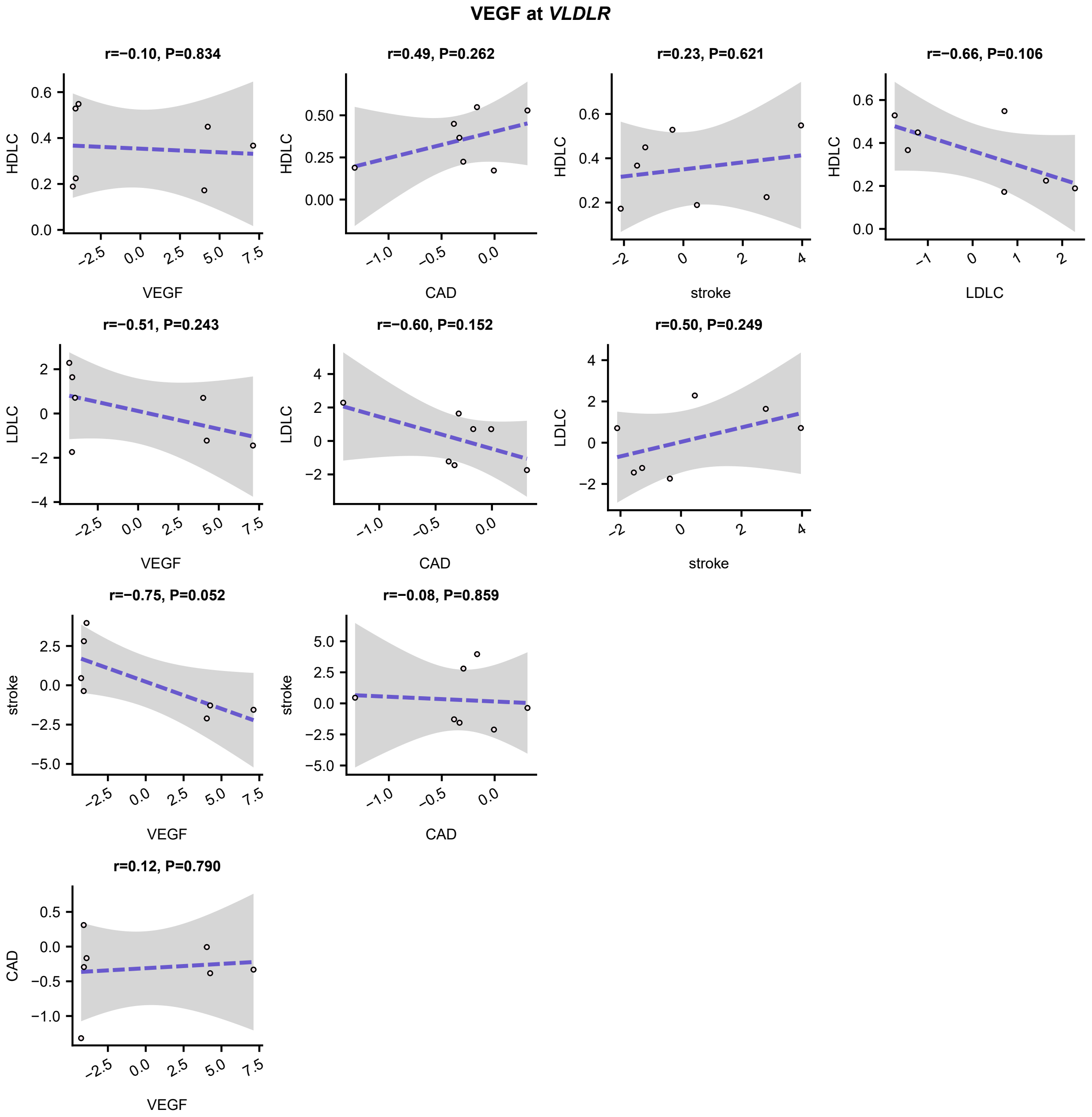

**Figure S3 H). SNP effect correlations as scatter plots; VEGF at *VEGFA.***

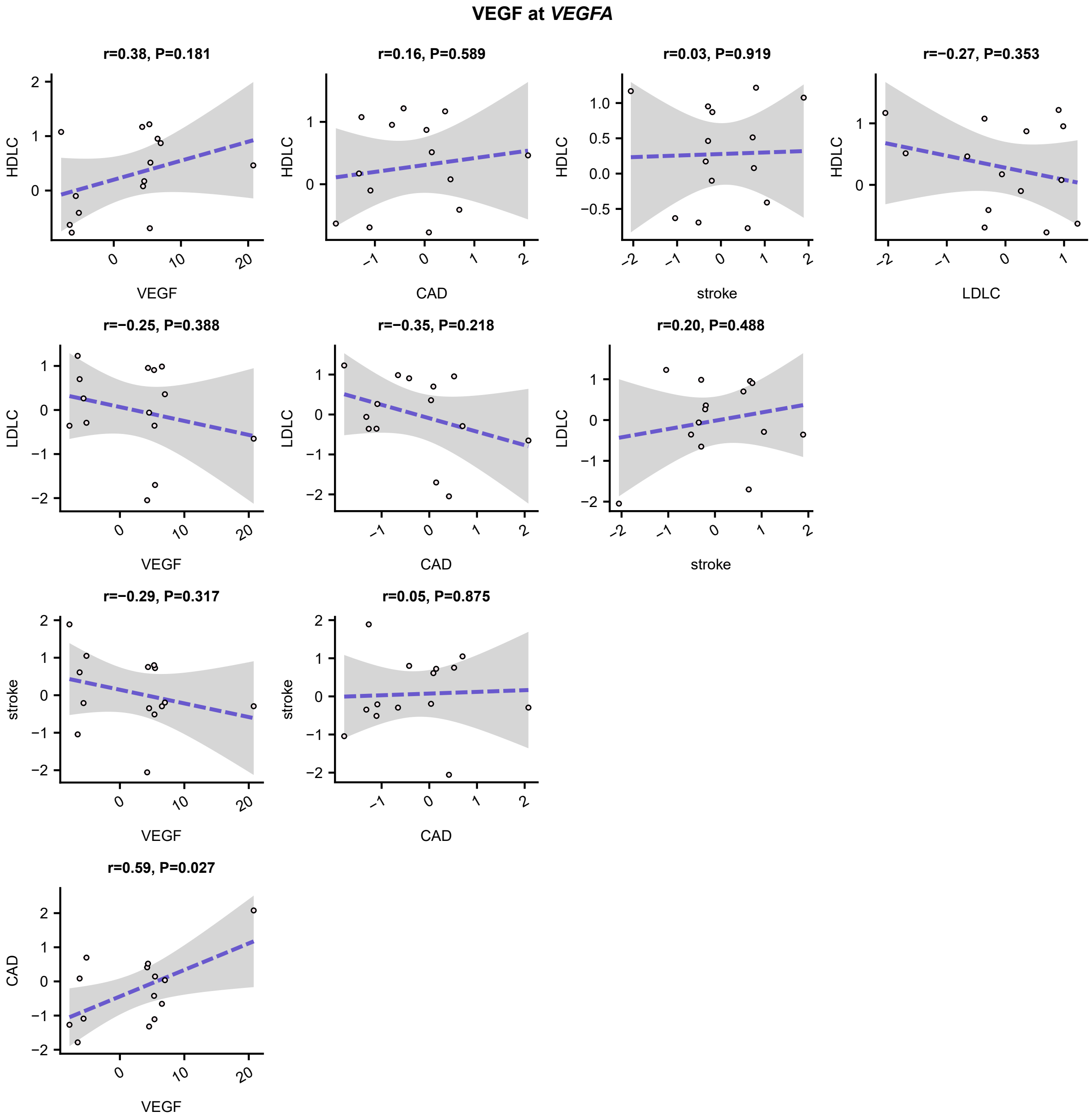

**Figure S3 I). SNP effect correlations as scatter plots; MCP1 at *ACKR1*.**

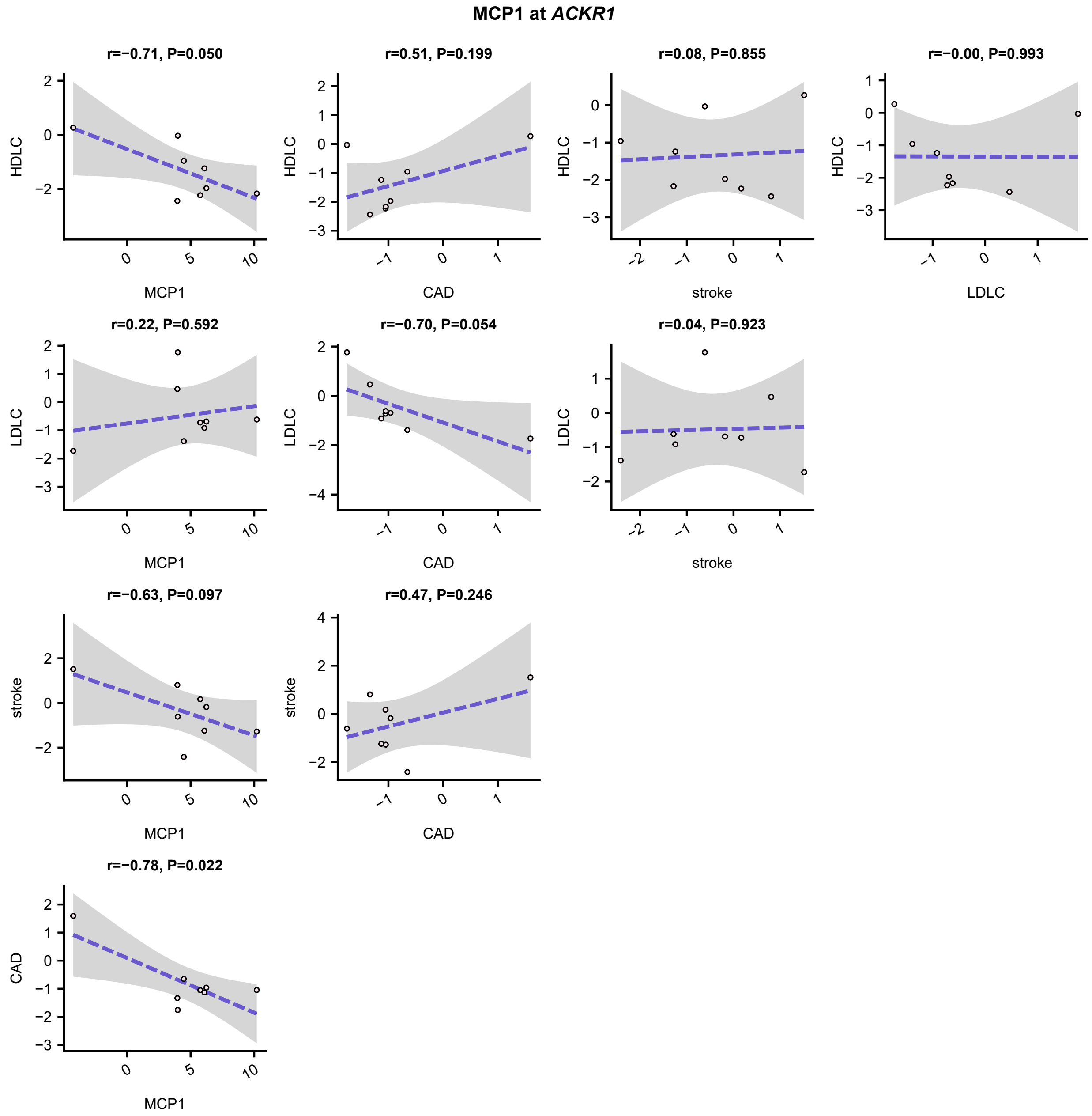
